# Functional interrogation of twenty type 2 diabetes-associated genes using isogenic hESC-derived β-like cells

**DOI:** 10.1101/2023.05.07.539774

**Authors:** Dongxiang Xue, Narisu Narisu, D. Leland Taylor, Meili Zhang, Caleb Grenko, Henry J. Taylor, Tingfen Yan, Xuming Tang, Neelam Sinha, Jiajun Zhu, J. Jeya Vandana, Angie Chi Nok Chong, Angela Lee, Erin C. Mansell, Amy J. Swift, Michael R. Erdos, Ting Zhou, Lori L. Bonnycastle, Aaron Zhong, Shuibing Chen, Francis S. Collins

## Abstract

Genetic studies have identified numerous loci associated with type 2 diabetes (T2D), but the functional role of many loci has remained unexplored. In this study, we engineered isogenic knockout human embryonic stem cell (hESC) lines for 20 genes associated with T2D risk. We systematically examined β-cell differentiation, insulin production and secretion, and survival. We performed RNA-seq and ATAC-seq on hESC-β cells from each knockout line. Analyses of T2D GWAS signals overlapping with HNF4A-dependent ATAC peaks identified a specific SNP as a likely causal variant. In addition, we performed integrative association analyses and identified four genes (*CP, RNASE1, PCSK1N* and *GSTA2*) associated with insulin production, and two genes (*TAGLN3* and *DHRS2*) associated with sensitivity to lipotoxicity. Finally, we leveraged deep ATAC-seq read coverage to assess allele-specific imbalance at variants heterozygous in the parental hESC line, to identify a single likely functional variant at each of 23 T2D GWAS signals.

## Introduction

Type 2 diabetes (T2D) is a major contributor to the global burden of disease, representing the 9th leading cause of death and costing over $700 billion in health expenditures annually.^1^ It is characterized by impaired insulin secretion in pancreatic islet β cells and reduced insulin response in insulin-sensitive tissues.^2^ Although the risk factors leading to insulin dysregulation are diverse,^3^ large-scale genetic studies have demonstrated the role of genetics in T2D susceptibility.^4^ The most recent, diverse, and largest studies have identified >290 distinct signals associated with T2D risk.^5–7^ However, despite success in identifying T2D-associated genetic effects, the challenge of understanding the molecular and cellular mechanisms driving these associations remains difficult, as many of the identified signals lie in non-coding regions of the genome, obscuring the identity of the underlying effector gene(s).^8^ Of the signals where a candidate effector gene is known, few of these genes have been investigated through detailed functional studies in model systems of disease relevant tissues such as pancreatic islets.^9, 10^ As the catalog of effector genes underlying T2D genetic associations accelerates in growth (e.g., through rare variant association studies^11, 12^ and improved molecular and computational identification techniques^13–16)^, the T2D research community needs efficient model systems to probe the molecular and cellular consequences of perturbed candidate effector genes.

Recent advances in stem cell and CRISPR-based technologies provide an opportunity to develop and apply such model systems in relevant tissues. Given the pathophysiology of T2D, much current genetic evidence supports the central role of pancreatic β cell development and dysfunction in T2D disease progression.^13, 17^ Robust protocols to differentiate human pluripotent stem cells (hPSCs), including both human embryonic stem cells (hESCs) and induced pluripotent stem cells (iPSCs), into insulin producing β-like cells have enabled *in vitro* model systems to study β-cell development.^18–20^ Coupled with the advent of flexible, facile gene editing technologies such as CRISPR-Cas9^21^, genetically engineered hPSCs promise to be an effective toolkit to investigate the effect of T2D-implicated genes on β-cell dysfunction. Indeed, recent studies, including ones from our group, have implemented this model system to generate isogenic hPSC-derived pancreatic β-like cells using different cell lines and characterize the effect of T2D-implicated genes, one at a time using a subset of assays we used here —including *NEUROG3*, *PDX1*, *GLIS3*, *ARX*, *GATA6*, *SIX2, ABCC8*, *KCNQ1*, *KCNJ11, CDKAL1*, and *MT1E*—on β cell differentiation, function, and survival.^22–31^ However, these efforts studied only a limited set of genes; the polygenic nature of T2D demands for larger studies to probe candidate effector genes more comprehensively.

In this study, we employ an efficient CRISPR-based platform to generate isogenic knockout (KO) hESCs across 20 genes with varying risk for T2D. We then differentiate KO hESCs as well as two wildtype (*WT*) control hESCs into insulin-producing β-like cells (**Figure 1A, Table 1**). We assess the effect of each KO across five different cellular phenotypes, including β cell differentiation efficiency, measures of insulin production and secretion, and β cell survival after lipotoxic exposure. All but one of the 20 gene KOs altered cellular phenotypes when compared to *WT* hESCs. To understand the molecular mechanisms driving these differences, we generate gene expression and chromatin accessibility profiles of purified insulin-expressing β-like cells and characterize the transcriptional and epigenetic alterations caused by the loss of T2D-associated genes. We test for an association of these transcriptional and epigenetic changes with cellular traits and find 21 genes associated with insulin content and 35 genes associated with β cell survival. We select 9 gene-cellular phenotype associations and demonstrate a causal effect in 6/9 using a human β cell line. Finally, we used the extensive chromatin accessibility data across all 22 hESC lines to test for allelic imbalance at variants that are in 99% credible sets for T2D (i.e., variants likely to be causal for T2D) and are heterozygous in the parental hESC line. At 23 T2D signals, we identify a single variant in the credible set with evidence of allelic imbalance, adding support to the role of these variants as the causal variant.

**Figure 1.**
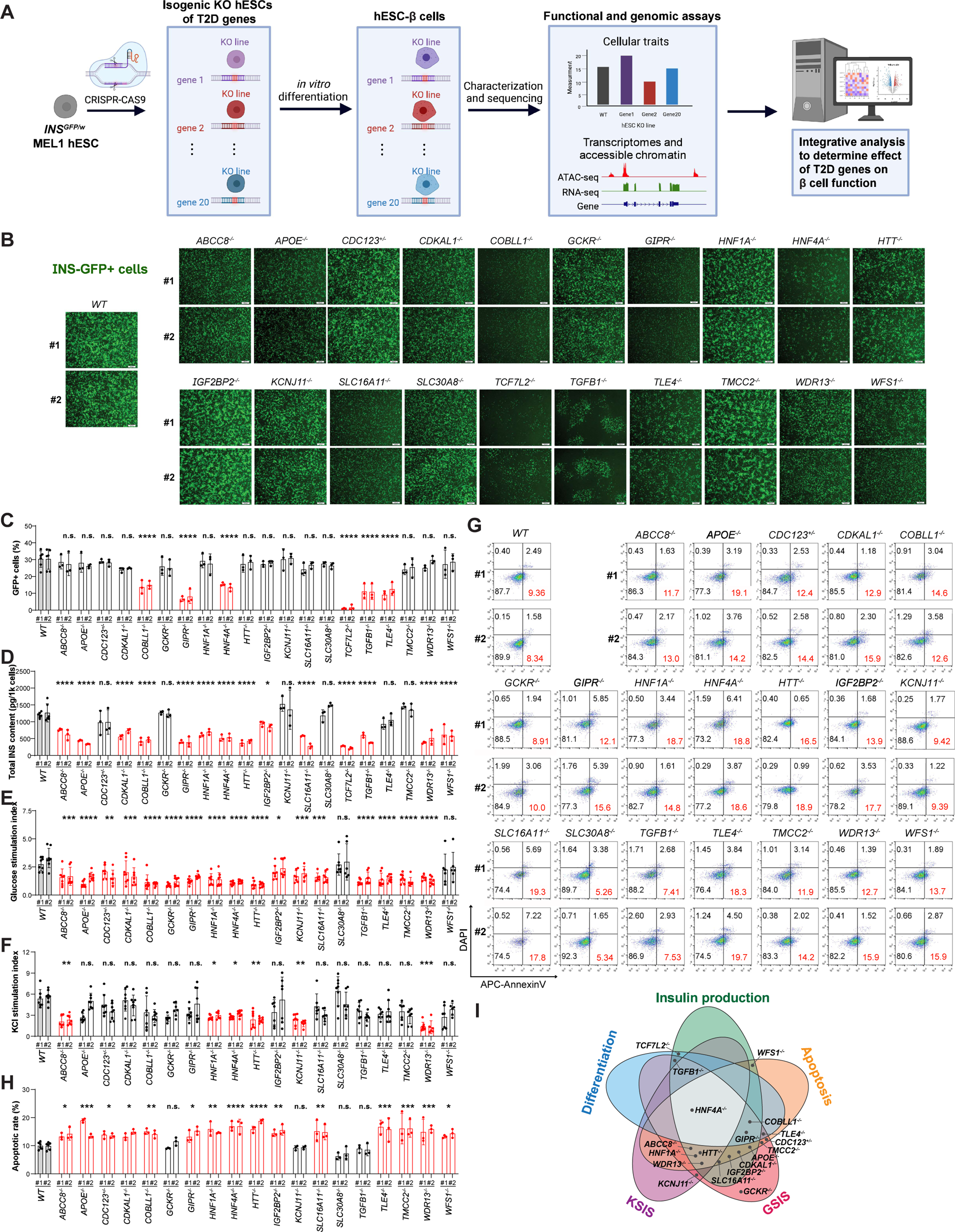
Isogenic hESC lines to evaluate the impact of loss of T2D associated genes in human pancreatic β cell generation, function, and survival. **(A)** Schematic illustration of the experimental design. Isogenic knockout (KO) hESCs were generated by individual KO of 20 T2D-associated genes in *INS^GFP/w^* MEL1 hESCs using CRISPR-Cas9 technology. The isogenic hESCs were differentiated to pancreatic β-like cells to evaluate the roles of T2D-associated genes in human pancreatic β cell generation, function, and survival. Integrative analysis of functional and genetic traits of 20 KO hESC-β cells was conducted to identify the cellular consequences of each knockout, and to search for possible hub genes controlling β cell failure. The figure was generated with BioRender.com with publication licenses. **(B)** Representative images at day 16 of differentiated cells derived from *WT* and isogenic KO *INS*^GFP/w^ hESC lines, which contain a GFP transgene driven by an insulin promoter. Thus, GFP+ cells are expected to represent insulin-producing β cells. Knockout lines with significant defects in β cell differentiation exhibit lower GFP signals, and biological replicates appear highly concordant. (Scale bar = 200 μm). **(C)** Quantification of the percentage of INS-GFP+ cells in differentiated cells (day 24) derived from *WT* and isogenic KO *INS*^GFP/w^ hESC lines. Y-axis represents percent INS-GFP+ cells which were determined by flow cytometry analysis (**Figure S2B**). Red bars represent a reduction in differentiation efficiency (*P*-value<0.05; Dunnett’s test). N=3-6 biological replicates per clone. **(D)** ELISA analysis of total intracellular insulin content of the purified (i.e. sorted GFP+) β-like cells derived from *WT* and isogenic KO hESCs. Red bars represent a reduction in total insulin content (*P*-value<0.05; Dunnett’s test). N=3-6 biological replicates per clone. **(E and F)** Static glucose-stimulated insulin secretion (GSIS) (**e**) and static KCl-stimulated insulin secretion (KSIS) (**F**) of day 30 hESC-islet cells derived from *WT* and isogenic KO hESCs. Stimulation index represents the fold change of the percent of insulin secreted upon 20 mM glucose stimulation divided by the percent of insulin secreted upon 2 mM glucose stimulation. N=6-8 biological replicates per clone. **(G and H)** Representative flow cytometry analysis (**G**) and the quantification of the percentage of Annexin V+DAPI-cells (**H**) in isogenic hESC-derived INS-GFP+ cells after treatment with 1 mM palmitate for 3 days. The gating strategy for analysis of apoptotic rate in hESC-derived β cells is shown in **Figure S2D**. N=3-6 biological replicates per clone. **(i)** Summary of the knockout impact of T2D-associated gene on five measured cellular traits of hESC-β cells. For panels **1C**-**1F** and **1H**, data was shown as mean ± SD of two independent clones (#1 and #2) for each hESC line. Number of biological replicates was listed in **Table S2**. *P*-values were calculated by one-way ANOVA followed by Dunnet’s test. The n.s. indicates a non-significant difference (P-value >0.05) and * symbol illustrates the significant difference of each KO line compared to the *WT* line. * P < 0.05, ** P < 0.01, ***P < 0.001, ****P < 0.0001.

**Table 1.** 20 T2D associated genes selected for creation of isogenic KO hESCs. The list was developed based on evidence from T2D genetics studies and expression in primary human islet β cells and hESC-derived β cells.

| Gene | Gene description | T2D evidence | Expression (TPM) |  |
| --- | --- | --- | --- | --- |
| | | | hESC-β cells | human islet $\beta$ cells |
| *ABCC8* | ATP-binding cassette sub-family C member 8 | T2D knowledge portal effector gene<sup>88,89,90,91</sup>; T2D association from exome-sequencing<sup>12</sup> | 53.75 | 31.27 |
| *APOE* | Apolipoprotein E | T2D knowledge portal effector gene<sup>88,89</sup> | 667.30 | 0.62 |
| *CDC123* | Cell division cycle protein 123 homolog | T2D knowledge portal effector<sup>5,91</sup>; T2D association from exome-sequencing<sup>12</sup> | 69.85 | 12.08 |
| *CDKAL1* | CDK5 Regulatory Subunit Associated Protein 1 Like 1 | T2D knowledge portal effector gene<sup>89,91</sup> | 79.59 | 2.14 |
| *COBLL1* | Cordon-bleu WH2 repeat protein like 1 | T2D knowledge portal effector gene<sup>89,91</sup>; T2D association from exome-sequencing<sup>12</sup> | 9.39 | 1.07 |
| *GCKR* | Glucokinase regulatory protein | T2D knowledge portal effector gene<sup>88,89,91</sup>; T2D association from exome-sequencing<sup>12</sup> | 51.29 | 0.01 |
| *GIPR* | Gastric inhibitory polypeptide receptor | T2D knowledge portal effector gene<sup>88,89,91</sup> | 90.23 | 3.23 |
| *HNF1A* | Hepatocyte nuclear factor 1-alpha | T2D knowledge portal effector gene<sup>88,89,90,91</sup> | 48.05 | 0.31 |
| *HNF4A* | Hepatocyte nuclear factor 4-alpha | T2D knowledge portal effector gene<sup>88,89,90,91</sup> | 102.97 | 0.66 |
| *HTT* | Huntingtin | T2D knowledge portal effector gene<sup>90</sup> | 28.73 | 0.66 |
| *IGF2BP2* | Insulin-like growth factor 2 mRNA-binding protein 2 | T2D knowledge portal effector gene<sup>5,88,89,90,91</sup> | 103.04 | 1.22 |
| *KCNJ11* | ATP-sensitive inward rectifier potassium channel 11; Kir6.2 | T2D knowledge portal effector gene<sup>5,88,89,90,91</sup>; T2D association from exome-sequencing<sup>12</sup> | 51.03 | 2.50 |
| *SLC16A11* | Monocarboxylate transporter 11 | T2D knowledge portal effector gene<sup>88,89,90</sup>; T2D association from exome-sequencing<sup>12</sup> | 29.52 | 0.67 |
| *SLC30A8* | Zinc transporter 8 | T2D knowledge portal effector gene<sup>88,89,90,91</sup>; T2D association from exome-sequencing<sup>12</sup> | 73.95 | 71.32 |
| *TCF7L2* | transcription factor 7 like 2 | T2D knowledge portal effector gene<sup>5,88-90</sup> | 18.96 | 3.42 |
| *TGFB1* | Transforming growth factor β-1 | T2D association from exome-sequencing<sup>12</sup> | 58.31 | 5.49 |
| *TLE4* | Transducin-like enhancer protein 4 | Nearest gene to T2D association<sup>8</sup> | 10.26 | 1.96 |
| *TMCC2* | Transmembrane and coiled-coil domains protein 2 | T2D knowledge portal effector gene<sup>90</sup>; T2D association from exome-sequencing<sup>12</sup> | 7.93 | 0.89 |
| *WDR13* | WD repeat-containing protein 13 | T2D association from exome-sequencing<sup>12</sup> | 32.36 | 12.38 |
| *WFS1* | Wolframin | T2D knowledge portal effector gene<sup>5,90,91</sup>; T2D association from exome-sequencing<sup>12</sup> | 163.18 | 7.38 |

## Results

### Generation and functional characterization of isogenic T2D-KO hESC lines

For the isogenic knockout lines, we selected 20 candidate effector genes with various degrees of evidence from T2D genetic studies (**Table 1**). To ensure that the genes played some role in islet β cells and could be studied using hESC-derived β-like cells, we only selected genes that were expressed in primary human islet β cells and hESC-derived β cells.

We generated isogenic knockout lines using an *INS^GFP/w^* MEL1 hESC reporter line^32^ that carries a GFP transgene driven by the human insulin promoter, enabling the isolation of insulin expressing cells by fluorescence-activated cell sorting (FACS; Methods). Briefly, we electroporated *INS^GFP/w^* MEL1 cells with a vector expressing Cas9 and a sgRNA targeting the first or second common exon of each T2D gene (**Table S1**). After subcloning, we expanded the isogenic hESC clones and selected two isogenic biological replicate clones for each T2D gene (labeled as #1 and #2) carrying either homozygous or compound heterozygous frameshift mutations, which we confirmed using Sanger DNA sequencing (**Figure S1**; Methods). For all target genes, we documented loss of function (LoF) mutations on both copies of the chromosome apart from *CDC123*, which was heterozygous LoF (*CDC123* is a cell division cycle protein; homozygous LoF would impair the cell division^33, 34^). In addition, as controls for subsequent analyses, we selected two wild type (*WT*) clones—one of which was exposed to Cas9 without the targeting sgRNA, the other of which was the unexposed *INS^GFP/w^*MEL1 cells. For all of the selected lines, we used immunocytochemistry staining to confirm that each clone retained typical hESC colony morphology and expressed pluripotency markers, including OCT4, SSEA4, NANOG, and TRA-1-81 (**Figure S2A**).

We differentiated the 22 hESC lines (2 *WT* lines, 20 KO lines with 2 biological replicate clones of each KO line; **Table S2**) into pancreatic β-like cells (hESC-β cells; Methods). We performed live cell imaging (Methods) and confirmed the presence of INS-GFP+ cells derived from the two *WT* lines (**Figure 1B**). For the KO lines, we observed variable representation of INS-GFP+ cells (**Figure 1B**), suggesting that some of the T2D-associated genes may affect differentiation efficiency. We further quantified the percent of GFP+ cells using flow cytometry and statistically tested for differences between the *WT* lines and each KO line grouped by the T2D KO gene (Methods). We found that the *COBLL1^−/−^, GIPR^−/−^, HNF4A^−/−^, TCF7L2^−/−^, TGFB1^−/−^*, and *TLE4^−/−^* lines showed impaired differentiation efficiency (*P*-value<0.05, Dunnett’s test; **Figure 1C, Figure S2B**). Notably, for the *TCF7L2^−/−^* line, the effect on differentiation efficiency was so severe that we dropped this line from some of the subsequent functional experiments that required many β-like cells (e.g., insulin secretion assays; Methods).

Next, we measured three insulin-related cellular traits (Methods): (i) total intracellular insulin content of the purified INS-GFP+ cells; (ii) insulin secretion after stimulation with 20 mM glucose (glucose stimulated insulin secretion [GSIS] index), and (iii) insulin secretion after stimulation with 30 mM KCl (KCl stimulated insulin secretion [KSIS] index). First, we measured the total intracellular insulin content in purified INS-GFP+ hESC-β cells using enzyme-linked immunosorbent assays (ELISAs) and tested for differences across all lines (Methods; **Figure 1D**). We detected decreased total intracellular insulin content in *ABCC8^−/−^*, *APOE^−/−^*, *CDKAL1^−/−^*, *COBLL1^−/−^*, *GIPR^−/−^*, *HNF1A^−/−^*, *HNF4A^−/−^*, *HTT^−/−^*, *IGF2BP2^−/−^*, *SLC16A11^−/−^*, *TCF7L2^−/−^*, *TGFB1^−/−^*, *WDR13^−/−^*, and *WFS1^−/−^* (*P*-value<0.05, Dunnett’s test). Next, we differentiated all of the lines apart from *TCF7L2^−/−^* into islet-like organoids (hESC-islets) and assessed GSIS, a measure independent of total insulin calculated by dividing the amount of insulin secreted upon 20 mM glucose stimulation by the amount of insulin secreted upon 2 mM glucose stimulation. Multiple KO hESC-islets exhibited impaired response to high glucose (*P*-value<0.05, Dunnett’s test; **Figure 1E**), including *ABCC8^−/−^*, *APOE^−/−^*, *CDC123*^+/−^, *CDKAL1^−/−^*, *COBLL1^−/−^*, *GCKR^−/−^*, *GIPR^−/−^*, *HNF1A^−/−^*, *HNF4A^−/−^*, *HTT^−/−^*, *IGF2BP2^−/−^*, *KCNJ11^−/−^*, *SLC16A11^−/−^*, *TGFB1^−/−^*, *TLE4^−/−^*, *TMCC2^−/−^* and *WDR13^−/−^*. Finally, we assessed KSIS in hESC-islets by dividing the amount of insulin secreted upon 30 mM KCl stimulation by the amount of insulin secreted under the control condition (upon 2 mM glucose stimulation). We found that loss of *ABCC8, HNF1A, HNF4A, HTT, KCNJ11* and *WDR13* resulted in decreased insulin secretion in response to 30 mM KCl (*P*-value<0.05, Dunnett’s test; **Figure 1F**).

As a final cellular phenotype, we evaluated the apoptotic rate of isogenic *WT* and mutant hESC-derived INS-GFP+ cells by quantifying the percent of AnnexinV+DAPI-cells (**Figure S2D**). Under regular cell culture conditions, we did not observe differences in β cell survival between *WT* and KO hESC-derived population (**Figure S2E**). Given the importance of pancreatic β cell death induced by lipid accumulation and stress in the context of T2D,^35, 36^ we assayed the apoptotic rate of INS-GFP+ hESC-β cells after exposing cells to 1 mM palmitate for 3 days to simulate lipotoxicity. Comparing the KO lines to *WT* lines, we observed increased palmitate-induced β cell apoptosis in *ABCC8^−/−^*, *APOE^−/−^*, *CDC123^+/−^*, *CDKAL1^−/−^*, *COBLL1^−/−^*, *GIPR^−/−^*, *HNF1A^−/−^*, *HNF4A^−/−^*, *HTT^−/−^*, *IGF2BP2^−/−^*, *SLC16A11^−/−^*, *TLE4^−/−^*, *TMCC2^−/−^*, *WDR13^−/−^*, and *WFS1^−/−^* (*P*-value<0.05, Dunnett’s test; **Figure 1G-H**).

In summary, of the 20 T2D genes knocked out in the isogenic KO lines, we found 6 caused defective differentiation, 14 decreased total intracellular insulin content, 17 impaired response to glucose stimulation, 6 impaired response to potassium stimulation, and 15 increased sensitivity to lipotoxicity. Interestingly, 5 genes—*ABCC8, HNF1A, HNF4A, HTT*, and *WDR13*—showed effects across all three β cell functional traits pertaining to insulin (total insulin content, GSIS, and KSIS; **Figure 1I**) and also β cell susceptibility to lipotoxicity (apoptotic rate; **Figure 1I**). Overall, 19 of the 20 T2D-associated genes showed an effect in at least one of the cellular assays considered, supporting our hypothesis that these 20 genes may affect T2D risk in part by perturbing the generation, function, and survival of pancreatic β cells. Strikingly, the loss of *HNF4A* affected all five cellular traits, suggesting a role of *HNF4A* in the development and function of pancreatic β cells (**Figure 1I**).

### Knockout of T2D genes results in large-scale transcriptomic and chromatin accessibility changes

We selected one clone for each KO line along with the two *WT* lines (22 total lines), differentiated each with replication, purified INS-GFP+ hESC-β cells, and performed RNA-seq as well as ATAC-seq (**Table S2**; Methods). After quality control procedures, we quantified the expression of 20,068 genes and 208,945 accessible chromatin regions across all lines (Methods). We evaluated the reproducibility of signals across replicates and found that the expression/accessibility measurements were highly reproducible (minimum Pearson’s *r*≥0.95).

For each KO line, we compared gene expression and chromatin accessibility signals to that of the *WT* lines (Methods). Focusing on the differentially expressed genes (DEGs; false discovery rate [FDR]<5% and |fold change|>1.5), the range of the number of DEGs varied widely, from 295 genes in *SLC30A8^−/−^* hESC-β cells to 5,969 genes in *HNF4A^−/−^* hESC-β cells (**Figure 2A**). We found 171 out of 257 genes previously reported as T2D effector genes (Methods) were DEGs in at least one line, including *PPARG* (*HNF4A*^−/−^, *HNF1A*^−/−^), *PAX4 (IGF2BP2^−/−^*), and *NEUROG3 (TCF7L2^−/−^*; **Figure S3A**). In addition, we estimated the enrichment of DEGs in genes binned by their expression specificity in primary islet β cells compared to other islet cell types (derived from a previous single cell RNA-seq study^37^; Methods). We observed that genes with expression profiles highly specific to β cells were enriched (FDR<5%) in DEGs for 17/20 of the KO lines (**Figure 2B**), underscoring the relevance of hESC-β cells as a model for primary islet β cells. These β cell-specific genes (**Figure S3B**) also included many well-characterized T2D genes such as *G6PC2*^38^ and *NKX6-1*^39^.

**Figure 2.**
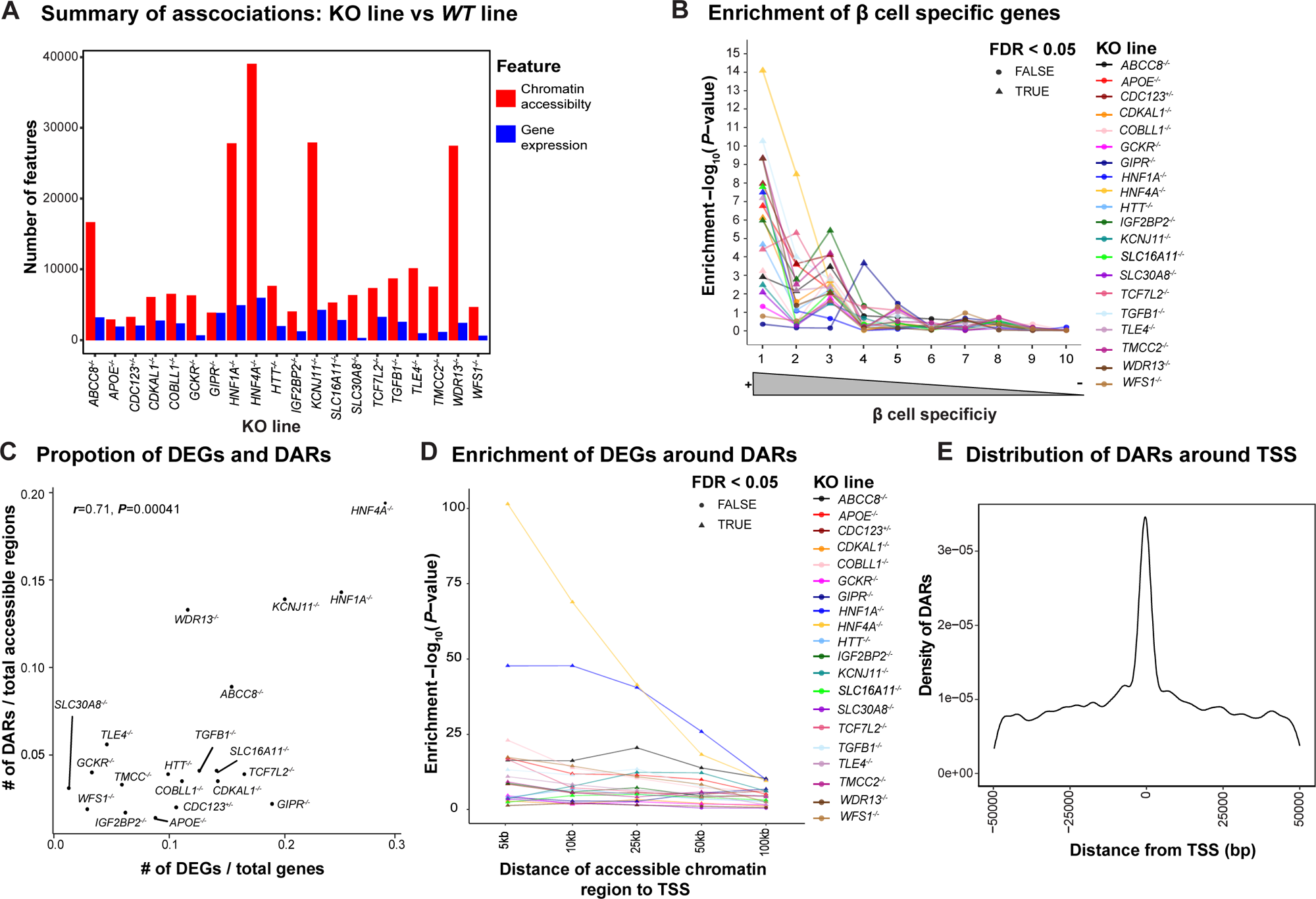
Loss of T2D associated genes results in large scale transcriptomic and epigenetic changes in hESC-β cells. **(A)** Summary of differential gene expression and differential chromatin accessibility in INS-GFP+ β-like cells derived from isogenic *WT* or KO hESC lines. Here x-axis= KO lines, and y-axis=number of features (differentially expressed genes in blue, differentially accessible chromatin regions in red (FDR<0.05 and |fold change|>1.5). **(B)** Enrichment of DEGs in β cell specific genes. We grouped all genes expressed in human pancreatic islets into 10 bins using single cell RNA-seq data (Bonnycastle *et al*. 2019; Methods). The genes in bin 1 are expressed highly specifically in β cells, while those in bin 10 are expressed ubiquitously across different cell types. x-axis= bins of genes with decreasing order of the specificity score (left to right), y-axis=−log_10_(*P*-value, hypergeometric test). The DEG set in *HNF4A^−/−^* KO versus *WT* hESC-derived INS-GFP+ cells have the highest enrichment of β cell specific genes. Each line represents one KO line. Triangle indicates an enrichment of DEGs in β cell specific genes in the specified bin at FDR< 0.05. **(C)** Correlation of DEGs and DARs. x-axis=fraction of DEGs in total number of genes, and y-axis=fraction of DARs in total number of accessible regions. Pearson’s r and associated *P*-value are shown on top. Each dot represents one KO line. **(D)** Enrichment of DEGs around DARs in varying sizes of windows. x-axis=distance of DAR from transcription start sites (TSS) of DEG (5, 10, 25, 50, and 100kb), y-axis=−log_10_(Enrichment *P*-value, Fisher’s exact test test). Each line represents one KO line. Triangle indicates an enrichment of DEGs around DARs in the specified window at FDR< 0.05. **(E)** Distribution of accessible chromatin regions associated with nearby gene expression (Methods). The associated accessible regions are the most abundant around TSS of a nearby gene. x-axis=distance of ATAC-seq peaks from nearby TSSs in base pairs, y-axis=occurrences of ATAC-seq peaks. Number of associated ATAC-seq peaks taper off when the distance gets > 50kb.

By comparing ATAC-seq data from KO lines to *WT* lines (Methods), we also identified differentially accessible chromatin regions (DARs; FDR<5% and |fold change|>1.5). As with the DEG results, both *HNF4A^−/−^* and *HNF1A^−/−^* exhibited the greatest number of DARs, with 39,013 and 27,773 DARs identified respectively (**Figure 2A**). Indeed, across all KO lines, the proportion of DARs identified was strongly correlated with the proportion of DEGs identified (Pearson’s *r*=0.71, *P*-value=4.1×10^−4^; **Figure 2C**). Since accessible chromatin regions often mark regulatory elements that modulate nearby gene expression,^13^ we asked if DEGs were enriched nearby DARs (<100 kilobases [kb] at various window sizes; Methods). Indeed, we found that DEGs were enriched near DARs up to 100 kb away (FDR<5%) in all but two KO lines, *SLC30A8^−/−^* and *WDR13^−/−^* (**Figure 2D**). At closer distances (<=25 kb), all lines showed substantial enrichment (FDR<5%). Finally, we considered the distance between each DAR and the nearest transcription start site (TSS) and observed that a large proportion of DARs (>21.7%) occur within 25 kb of a TSS (**Figure 2E**), as would be expected if DARs marked regulatory elements.

To identify potential regulatory elements in hESC-β cells, we fit a regression to link accessible chromatin regions to nearby genes (<50kb) by jointly modeling ATAC-seq and RNA-seq signals across all 22 lines (KO+WT; Methods). We identified 1,150 associations (FDR<5%) spanning 726 genes and 1,035 accessible chromatin regions. While most genes were associated with a single chromatin region, we found that a few genes were associated with as many as 10 open chromatin peaks (**Figure S3C**). The same trend held true for accessible chromatin regions (**Figure S3D**). Notably, we found chromatin regions associated with T2D effector genes (e.g., *TMEM176A/B*; **Figure S3E**) and genes important in β cell differentiation (e.g., *NKX6-1*; **Figure S3E**).

### HNF4A regulates diabetes relevant genes and HNF4A binding sites are perturbed by genetic variants associated with T2D

Across all KO lines, *HNF4A^−/−^* affected all five functional readouts compared to *WT* lines (e.g., hESC-β cell differentiation efficiency, insulin content, insulin secretion index, apoptotic rate; **Figure 1I**), induced the greatest number of transcriptional and epigenomic changes compared to the *WT* lines (5,969 DEGs and 39,013 DARs respectively; **Figure 2A, 3A, 3B**), and resulted in DEGs with the greatest enrichment in β cell specific expression patterns (**Figure 2B**). We therefore further explored the role of *HNF4A* in the context of (i) hESC-β cell gene regulation and (ii) the genetics of T2D. First, we sought to understand the genes regulated by *HNF4A* and performed gene set enrichment of the differential gene expression results (Methods). We found that genes down-regulated in *HNF4A^−/−^* compared to *WT* were enriched (FDR<5%) in genes related to glucose metabolism (e.g., fructose and mannose metabolism) and maturity onset diabetes of the young (MODY; e.g., *PKLR, PAX4, HNF1A, NKX6-1, PDX1, NEUROG3, FOXA3*; **Figure 3C**), while up-regulated genes were enriched processes not clearly relevant to diabetes (e.g., “cell cycle”, “DNA replication”, “basal cell carcinoma”; **Figure S4A**). Such results suggest that HNF4A generally up-regulates diabetes relevant genes, consistent with previous reports of HNF4A as a transcriptional activator.^40, 41^

**Figure 3.**
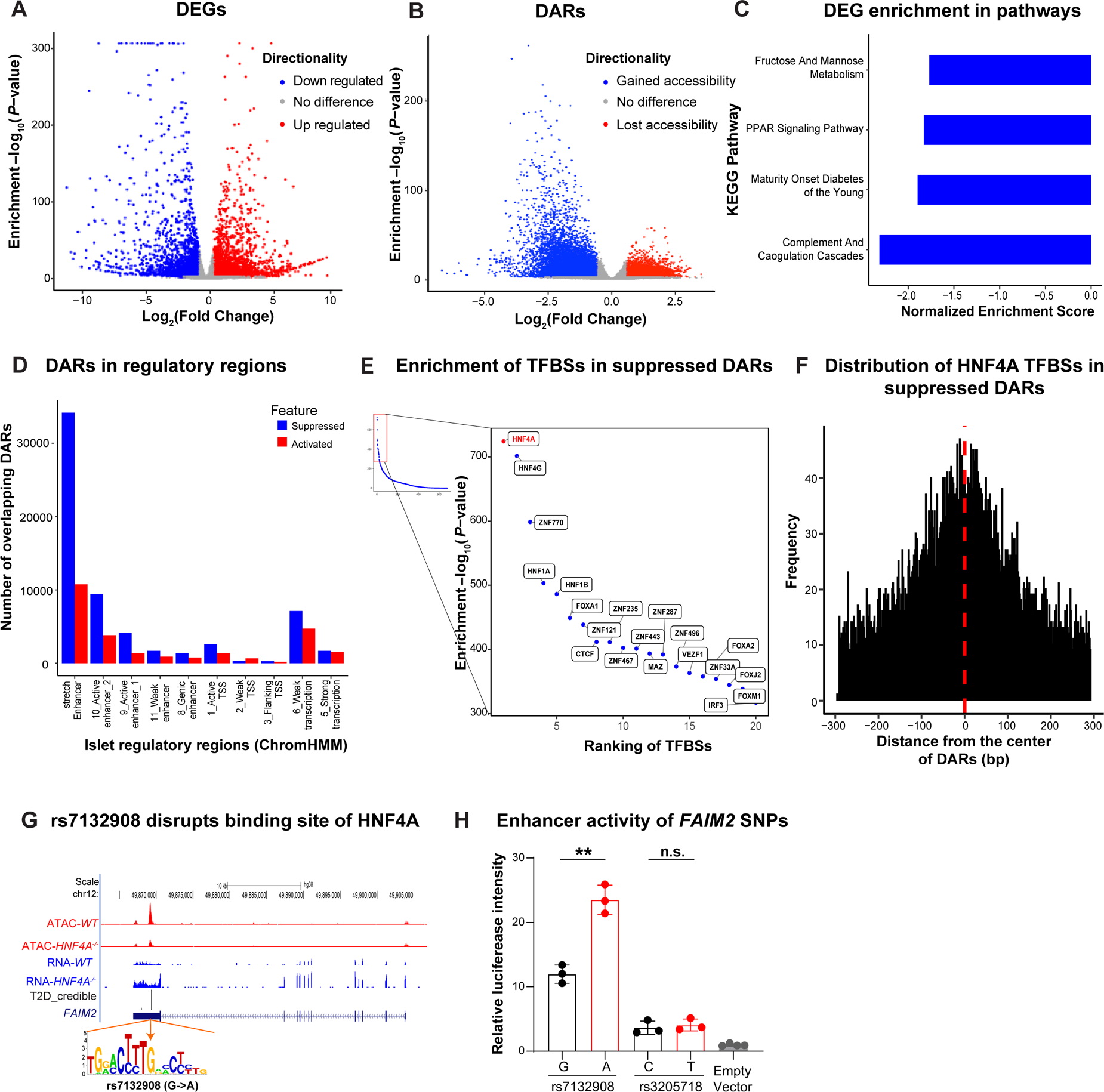
Fine mapping analysis of transcriptomic and epigenomic alterations in *HNF4A^−/−^* hESC-β cells prioritize a causal variant rs7132908 at a T2D risk locus. **(A)** DEGs of the INS-GFP+ cells derived from the *HNF4A^−/−^*versus *WT* hESC. Blue dots and red dots represent down- and up-regulated genes in *HNF4A*^−/−^ hESC-β cells, respectively. x-axis=Log_2_(fold change), y-axis=−log_10_(*P*-value). Genes associated at FDR<0.05 and |fold change| > 1.5 are highlighted. **(B)** DARs of the purified INS-GFP+ cells derived from the *HNF4A^−/−^* hESC-β versus *WT* hESC-β cells. Blue dots and red dots represent suppressed- and activated DARs in *HNF4A*^−/−^ hESC-β cells, respectively. x-axis=Log_2_(fold change), y-axis=−log_10_(*P*-value). Chromatin accessible regions associated at FDR<0.05 and |fold change|>1.5 are highlighted. **(C)** KEGG pathways enriched with down-regulated genes in *HNF4A^−/−^* versus *WT* hESC (FDR < 0.05). x-axis=normalized enrichment scores per GSEA (Methods), y=KEGG pathways. **(D)** Overlap of DARs in the INS-GFP+ cells derived from the *HNF4A^−/−^*versus *WT* hESC with islet regulatory features defined by ChromHMM (Varshney *et al*. 2017). Suppressed DARs: blue, activated DARs: red. x-axis=islet regulatory features, ordered by stretch enhancer, enhancers, promoters and transcriptional regions (left to right). y-axis=counts of overlapping DARs adjusted by total number of respective regulatory regions (number of DARs overlapping a regulatory feature * 10,000/total number of regulatory regions). **(E)** Enrichment of HNF4A transcription factor binding site (TFBSs) motifs in suppressed DARs of the *HNF4A*^−/−^ hESC-β cells. Right panel shows the top 20 most enriched TFBSs. x-axis=ranking of enriched TFBSs (enrichment in decreasing order from left to right), y-axis=−log10(*P-*value) by MEME simple enrichment analysis (SEA) (Methods). Red dot highlights HNF4A binding site is the most enriched. **(F)** Relative distance of HNF4A TFBSs from the center of suppressed DARs in the *HNF4A*^−/−^ hESC-β cells in comparison to *WT* hESC-β cells. TFBS motif abundance was generated by scanning 150bp flanking regions around centers of all suppressed DARs using MEME fimo (Methods). x-axis=distance of the TFBS motif from the center of DARs, y-axis= frequency of the motif occurrences. **(G)** T2D credible set of SNPs at a locus on chromosome 12 (fmap.FAIM2.chr12:50263148) near *FAIM2*. rs7132908 overlaps a DAR and the A allele disrupts an HNF4A binding site. Top panel shows ATAC-seq (red) and RNA-seq (blue) read pileups in *WT* and *HNF4A*^−/−^ hESC-β cells. “T2D_credible” shows two T2D credible set SNPs (height of the bar represent PPA) of at the locus overlapping 3’ UTR of *FAIM2*. **(H)** Luciferase analysis to assess the functionality of the two credible set SNPs and an empty vector (negative control) in EndoC-βH1 cell. Both alleles of rs7132908 have enhancer activity, but the G allele (which matches the HNF4A consensus sequence) has lower activity. Thus, HNF4A, coded by a gene highly expressed in EndoC-βH1 cells (expression TPM=68^92^), is likely serving as a repressor in this instance. The second credible set SNP rs3205718 appears to have no differential activity for either allele and thus much less likely to be the causative SNP. x-axis=alleles tested for rs7132908 and rs3205718, y-axis=relative luciferase intensity. Data was shown as mean ± SD. There are 3 biological replicates for both rs7132908 and rs3205718 group and 4 biological replicates for empty vector control group. Unpaired Student’s t-test: ** *P* < 0.01.

Next, we expanded our characterization of the regulatory patterns of *HNF4A^−/−^* DARs. We observed that 58% of the 39,013 DARs in the line were suppressed while 42% were activated (Figure 3B). Among all the KO lines, *HNF4A^−/−^*showed the largest percentage of DEGs around DARs, with 79% of DEGs having a DAR within 50kb (*P*-value=3.1×10^−10^, Fisher’s exact test; **Figure 2D**; Methods). We hypothesized that such results may indicate that *HNF4A^−/−^* DARs occur in regulatory elements that may drive the observed changes in gene expression. We tested the enrichment of DARs in islet regulatory regions^42^(Methods). We observed strong enrichment (FDR<0.05) for diminished accessible regions (suppressed DARs in *HNF4A^−/−^* KO) in multiple islet regulatory regions (**Figure 3D**)—notably islet stretch enhancers, which generally regulate tissue/cell-type specific gene expression^43^ - while we did not find such an enrichment for activated DARs. We scanned the *HNF4A^−/−^* DARs using binding site motifs for 677 transcription factors expressed in *WT* hESC-β cells (Methods) and found that the suppressed DARs were most enriched in the HNF4A binding motif (FDR<5%; **Figure 3E**), while the activated DARs were most enriched in the FOXA1 binding motif (FDR<5%; **Figure S4B**). Selecting the HNF4A binding motif, we calculated the distance of the predicted binding site to the center of the accessible chromatin region (Methods). For the suppressed DARs the HNF4A binding motif most often occurred at the center of the region while for activated DARs there was no trend (**Figure 3F** and **Figure S4C**). These results suggest that many of the suppressed DARs might reflect direct changes due to binding of HNF4A while many activated DARs reflect indirect effects resulting from the *HNF4A* knockout.

Given the wide-spread diabetes relevant effects of *HNF4A^−/−^*on hESC-β cells, we investigated if predicted HNF4A binding sites may be perturbed by candidate causal variants within 99% credible sets for genetic associations with T2D^5^ (Methods). We focused on binding sites in suppressed DARs in *HNF4A^−/−^* compared to the *WT* lines and identified 2 variants, representing 2 T2D signals, predicted to affect HNF4A TF footprint (**Table S3**). We selected the T2D genetic association signal at *FAIM2*^5^ for experimental follow-up that contained two credible SNPs, rs7132908 (posterior probability of association [PPA]=0.92) and rs3205718 (PPA=0.07). Of these two SNPs, only rs7132908 overlaps an HNF4A footprint, where the T2D risk allele, “A”, is predicted to disrupt HNF4A binding (**Figure 3G**). This HNF4A footprint occurred in a DAR, with decreased accessibility in *HNF4A^−/−^*compared to *WT*, and was not associated with expression of *FAIM2* or any other nearby gene (FDR>5%), making the effector gene unknown at this signal. We performed allele-specific luciferase assays for both variants in EndoC-βH1, a human pancreatic β cell line (Methods). We observed a change in luciferase activity of the alleles for rs7132908 (*P*-value=0.002, unpaired Student’s t-test) but not for rs3205718 (*P*-value=0.618, unpaired Student’s t-test; **Figure 3H**); the “A” allele of rs7132908 was associated with increased luciferase activity. Combined, these data suggest that rs7132908 is likely the causal variant at this T2D signal and that the “A” allele of rs7132908 increases T2D risk by decreasing HNF4A binding and increasing the strength of an enhancer. In this situation it thus appears that HNF4A is acting as a repressor.

### Association between gene expression and cellular traits identifies genes controlling insulin content and β cell survival

By comparing *WT* lines to KO lines spanning 20 genes, we identified downstream effects of T2D-relevant genes on cellular traits (e.g., insulin content, β cell survival, GSIS; **Figure 1C-H**), gene expression (**Figure 2A**), and chromatin accessibility (**Figure 2A**). In addition, the availability of cellular traits paired with omics measurements (RNA-seq and ATAC-seq) across gene perturbations created a dataset where one could begin to map regulatory networks for cellular traits, identifying genes and chromatin regions associated with each cellular trait. Across all cell lines with paired cellular trait and omics data, we jointly modeled gene expression and chromatin accessibility with each cellular trait (Methods). We identified 21 genes associated with insulin content and 35 genes associated with β cell survival after palmitate exposure (FDR<5% and |effect size|>1.5, **Table S4**). We did not find any chromatin regions associated with a cellular trait (FDR>5%).

Focusing on the 21 genes associated with total insulin content (**Figure 4A, Table S4**), we selected five protein-coding genes (**Figure 4B-F**) to test for a causal relationship with insulin content based on the effect size of the association (|effect size|>1.5) and the gene’s expression in human islets (TPM>5) and hESC-β cells (TPM>5). Using EndoC-βH1 cells, we perturbed the expression of these candidate genes by inhibiting the expression of genes positively correlated with insulin content—*CP* and *FOSB*—through CRISPR interference (CRISPRi) and activating the expression genes negatively correlated—*RNASE1, PCSK1N*, and *GSTA2*—through CRISPR activation (CRISPRa; Methods). Prior to testing for effects on insulin content, we confirmed that the expression of all five genes was reduced or activated as expected using real-time quantitative reverse transcription PCR (qRT-PCR; **Figure S5A, 5B;** Methods). For 4/5 of the selected genes, we observed the predicted effect on total insulin content (*P*-value<0.05, unpaired Student’s t-test; **Figure 4G-H**), where both the inhibition of *CP* and the activation of *RNASE1*, *PCSK1N*, and *GSTA2* decreased total insulin content. For *FOSB* inhibition, we observed no effect on total insulin content (*P*-value=0.32, unpaired Student’s t-test), suggesting that the observed correlation between *FOSB* and total insulin content was not driven by a causal relationship, or that EndoC-βH1 cells are not the right model for detecting this effect.

**Figure 4.**
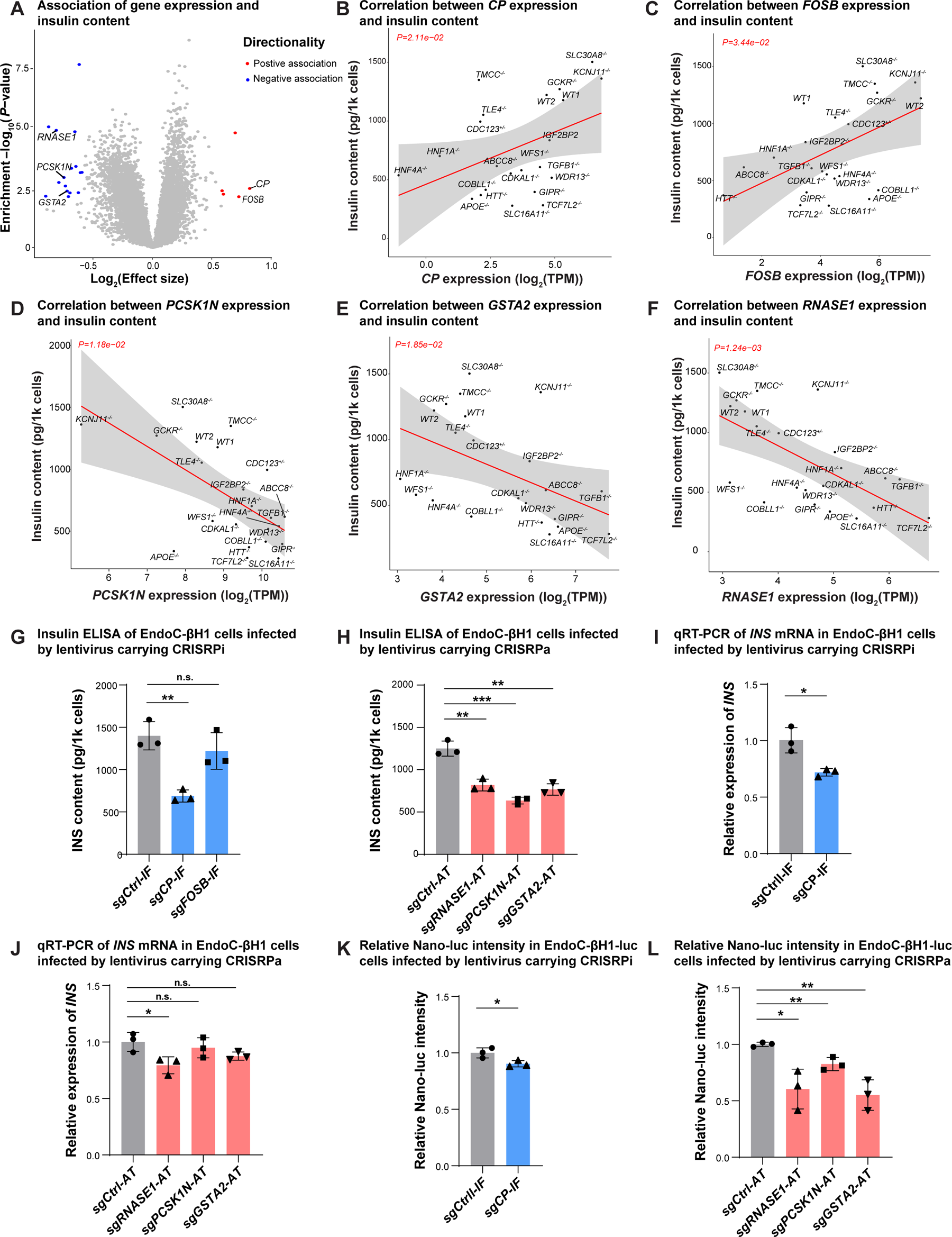
Cellular trait association analysis identifies potential genes controlling insulin content. **(A)** Identification of genes associated with total insulin content in hESC-β cells. A linear regression model was fitted to detect genes associated with total insulin content in purified INS-GFP^+^ cells derived from 20 KO and 2 *WT* hESCs (see Methods). x-axis=log_2_(effect size), which indicates the effect size with directionality, y-axis=−log_10_(*P*-value). Genes associated at FDR<0.05 and |effect size|>1.5 are colored and labeled (negative: blue, positive: red). **(B-F)** Association analysis of total insulin content *in each WT or KO line* with RNA expression of candidate gene *CP* (**B**), *FOSB* (**C**), *PCSK1N* (**D**), *GSTA2* (**E**) and *RNASE1* (**F**). x-axis=log2(TPM), y-axis= total insulin content (pg/1k cells). The solid line and gray area indicate the regression line and 95% confidence interval (CI), respectively. **(G)** Insulin ELISA analysis of total insulin content of EndoC-βH1 cells transfected with lentivirus carrying sgRNAs to transcriptionally inhibit expression of candidate genes *CP* or *FOSB* by CRISPR interference (IF). N=3 biological replicates. **(H)** Insulin ELISA analysis of total insulin content of EndoC-βH1 cells transfected with lentivirus carrying sgRNAs to transcriptionally activate expression of candidate genes *RNASE1, PCSK1N*, or *GSTA2* by CRISPR activation (AT). N=3 biological replicates. **(I)** qRT-PCR to monitor the relative expression of *INS* mRNA in EndoC-βH1 cells transfected with lentivirus carrying sgRNAs to transcriptionally inhibit expression of *CP*. N=3 biological replicates. **(J)** qRT-PCR to monitor the relative expression of *INS* mRNA in EndoC-βH1 cells transfected with lentivirus carrying sgRNAs to transcriptionally activate expression of *RNASE1, PCSK1N*, or *GSTA2*. N=3 biological replicates. **(K)** Relative luciferase intensity of EndoC-luc cells transfected with lentivirus carrying sgRNAs to transcriptionally inhibit expression of *CP*. N=3 biological replicates. Nano-Glo luciferase was cloned into the C-peptide portion of a proinsulin transgene driven by a CMV promoter and Nano-Glo luciferase (Nano-luc) intensity correlated with proinsulin transgene expression and processing. **(L)** Relative luciferase intensity of EndoC-luc cells transfected with lentivirus carrying sgRNAs to transcriptionally activate expression of *RNASE1, PCSK1N*, or *GSTA2*. Nano-luc intensity measures the c-peptide content. N=3 biological replicates. For panels **4G-4L**, data are shown as mean ± SD. *P*-values were calculated by unpaired Student’s t-test. The n.s. indicates a non-significant difference (*P*-value>0.05) and * symbol illustrates the significant difference of each genetic perturbation line compared to the control line. * P < 0.05, ** P < 0.01, ***P < 0.001, ****P < 0.0001.

To better understand the molecular mechanisms underlying the observed effects of *CP, RNASE1*, *PCSK1N*, and *GSTA2* on total insulin content, we conducted similar CRISPR perturbation experiments and measured (i) *INS* transcription and (ii) insulin protein translation/processing in EndoC-βH1-luc cells using the proinsulin Nano-Glo luciferase (Nano-luc) reporter as a readout (Methods). For *INS* transcription, we used qRT-PCR in EndoC-βH1 cells, revealing that decreased *CP* expression and increased *RNASE1* expression resulted in reduced *INS* expression (*P*-value<0.05, unpaired Student’s t-test), while changes in *PCSK1N* and *GSTA2* expression had no effect (**Figure 4I, 4J**). To assess insulin protein translation/processing, we generated EndoC-βH1-luc cells by inserting Nano-Glo luciferase into the C-peptide portion of a proinsulin transgene. Nano-luc could then be released via endogenous proinsulin convertase enzymes,^44^ and its intensity could be used to track the change of proinsulin transgene translation and processing upon CRISPR perturbation. Prior to the Nano-luc assay, we confirmed the reduced expression of *CP* and activated expression of *RNASE1*, *PCSK1N*, and *GSTA2* in EndoC-βH1-luc cells using qRT-PCR (**Figure S5B**; Methods). The Nano-luc assays showed that inhibition of *CP* had a mild effect on insulin protein translation/processing, reducing luciferase intensity by only ∼10% (*P*-value<0.05, unpaired Student’s t test; **Figure 4K**), while activation of *PCSK1N, GSTA2*, and *RNASE1* greatly reduced intracellular Nano-luc production (*P*-value<0.05, unpaired Student’s t-test; **Figure 4L**). Combined, these experiments confirm a causal role of *CP, RNASE1*, *PCSK1N*, and *GSTA2* in regulating insulin content. Our results suggest that *CP* primarily affects *INS* transcription, while *PCSK1N* and *GSTA2* have a greater impact on insulin protein translation/processing. For *RNASE1*, we detected strong effects on both *INS* transcription and insulin protein translation/processing, consistent with the established role of *RNASE1* in moderating mRNA stability.^45^

In addition, from the 35 genes associated with palmitate-induced β cell apoptotic rate (**Figure 5A, Table S4**), we selected five protein-coding genes to test for a causal relationship with palmitate-induced apoptotic rate based on the effect size of the association (|effect size|>1.5) and the gene’s expression in human islets (TPM>5) and hESC-β cells (TPM>5): *TAGLN3*, *ADCYAP1*, *DHRS2*, *CP*, and *SYNPO* (**Figure 5B-F**). Similar to the assessment of insulin content, we used CRISPRa to activate the expression of positively correlated genes—*TAGLN3, ADCYAP1*, and *DHRS2*— and CRISPRi to inhibit the expression of negatively correlated genes—*CP* and *SYNPO*—in EndoC-βH1 (Methods). Prior to conducting apoptosis experiments, we confirmed that the expression of all five genes was reduced or activated as expected using qRT-PCR (**Figure S5C**; Methods). After infecting EndoC-βH1 cells with lentivirus carrying the CRISPR construct and guide, we treated cells with 1 mM of palmitate for 3 days and quantified the apoptotic rate using flow cytometry analysis (Methods). We detected an increased apoptotic rate for *TAGLN3* and *DHRS2* (*P*-value<0.05, unpaired Student’s t-test; **Figure 5G-H**) but found no effects on apoptotic rates for the other genes (*P*-value>0.05, unpaired Student’s t-test; **Figure 5G-J**). Through immunofluorescence staining (Methods), we further confirmed that activation of *TAGLN3* and *DHRS2* led to an increased percentage of cleaved caspase 3+ cells (**Figure 5K-L**). Overall, these findings suggest that *TAGLN3* and *DHRS2* may play a critical role in regulating β-cell survival, while the other three correlated genes—*CP*, *ADCYAP1* and *SYNPO*—that did not exhibit effects on palmitate-induced β-cell apoptosis, may only be involved in the innate β-cell survival response to lipotoxicity but do not exert a direct role to regulate β-cell apoptosis.

**Figure 5.**
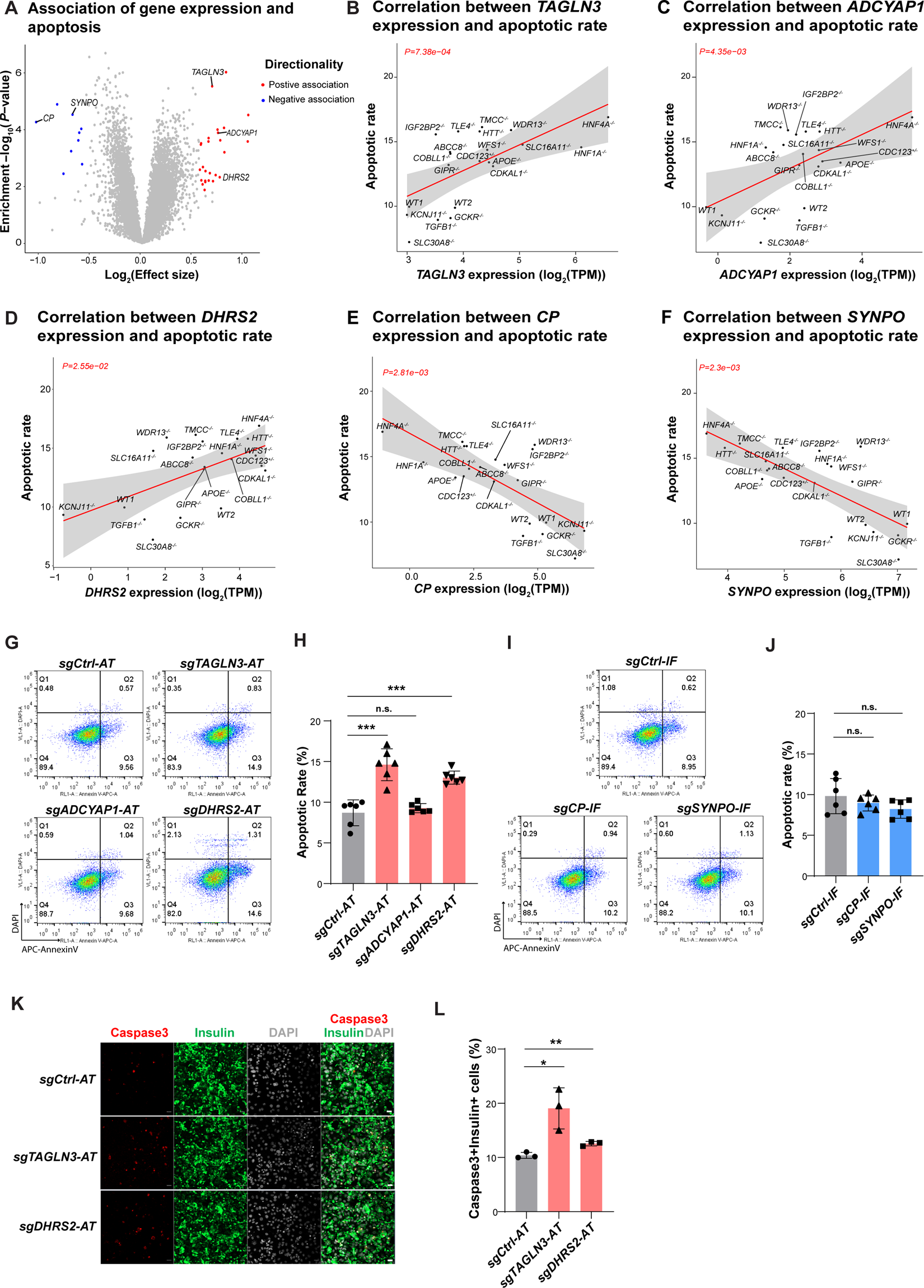
Cellular trait association analysis identifies genes controlling β cell survival. **(A)** Identification of genes correlated with palmitate-induced apoptotic rate in hESC-β cells. A linear regression model was fitted to detect genes associated with palmitate-induced apoptotic rate in INS-GFP^+^ cells derived from 20 KO and 2 *WT* hESCs (see Method). x-axis=log_2_(effect size), y-axis=−log_10_ (*P*-value). Genes associated at FDR<0.05 and |effect size|>1.5 are colored and labeled (negative: blue, positive: red). The apoptotic rate was defined as the percentage of Annexin V+/DAPI-cells. **(B-F)** Linear regression analysis of apoptotic levels *in each WT or KO line* with RNA expression of candidate genes *TAGLN3* (**B**)*, ADCYAP1*(**C**), *DHRS2* (**D**), *CP* (**E**) and *SYNPO* (**f**). The degree of apoptosis in hESC-β cells was assessed after treatment with 1mM palmitate for 3 days. x-axis=log2(TPM), y-axis=apoptotic rate (%). The solid line and gray area indicate the regression line and 95% CI, respectively. **(G-J)** Representative flow cytometry analysis (**G** and **I**) and the quantification of the percentage of AnnexinV+DAPI-cells (**h** and **j**) in genetic perturbed EndoC-βH1 cells after treatment with 1 mM palmitate for 3 days. Positively correlated genes-*TAGLN3*, *ADCYAP1* and *DHRS2*, were transcriptionally activated by CRISPR activation (AT, **G** and **H**) while negatively correlated genes-*CP* and S*YNPO*, were transcriptionally suppressed by CRISPR interference (IF, **I** and **J**). Gating strategy is shown in **Figure S5D**. N=6 biological replicates. **(K** and **L)** Representative Immunofluorescent staining pictures (**K**), and the quantification of the percentage of Cleaved caspase3+Insulin+ cells (**L**), in EndoC-βH1 cells transfected with lentivirus carrying sgRNA to activate *TAGLN3* or *DHRS2*. N=3 biological replicates. Cleaved caspase 3 was presented in red and insulin was presented in green. Nucleuses were stained with DAPI (grey). Scale bar = 20 μm. For panels **5H, 5J** and **5L**, data are shown as mean ± SD. *P*-values were calculated by unpaired Student’s t-test. The n.s. indicates a non-significant difference (*P*-value>0.05) and * symbol illustrates the difference of each genetic perturbation line compared to the control line. * P < 0.05, ** P < 0.01, ***P < 0.001, ****P < 0.0001.

### Analysis of allelic imbalance in accessible chromatin regions identifies a single candidate causal variant at 23 T2D genetic associations

ATAC-seq quantifies the accessibility of genomic regions, for example driven by TF binding, and is capable of capturing instances where one allele is preferentially accessible (e.g., preferentially bound by a TF^46, 47^). Such events can be quantified by measuring the difference in allele counts at heterozygous variants. Since both alleles occur within the same cell and have been exposed to the same experimental conditions, the intra-sample nature of this metric greatly reduces noise and maximizes signal.

Using the chromatin accessibility data generated across the 20 KO lines and two *WT* lines, we quantified allelic imbalance (Methods) across SNPs in 99% credible sets for T2D genetic associations.^5^ We tested for allelic imbalance signatures common across all lines (common effect analysis; Methods) and identified 45 T2D association signals with >=1 SNP that showed allelic imbalance (FDR<5%; **Table S5**). At 18 of those signals, the *INS^GFP/w^*MEL1 cell line was heterozygous at all SNPs in the credible set and only one SNP showed allelic imbalance, which may represent the causal SNP at the T2D genetic association (**Figure 6A**). A possible explanation of these common effects is that they represent shared TF binding events common across all lines that are not disrupted by any of the LoF gene perturbations. However, it is also possible that a causal SNP may not manifest an effect across the majority of lines and rather be limited to a subset of lines or a single line. Therefore, we also tested for allelic imbalance within each line individually (line-specific effect analysis; Methods). At an FDR<5%, we identified 25 signals with >=1 SNP that showed allelic imbalance in >=1 line (**Table S5**). Focusing on the T2D signals where *INS^GFP/w^* MEL1 was heterozygous at all SNPs in the credible set, we found 12 signals where only one SNP showed allelic imbalance, including 5 signals where a single SNP was not identified when considering common allelic imbalance effects across all lines **(Figure S6)**. Of the 18 signals identified previously, 7 signals showed an effect at the individual line level, all at the previously identified SNP. Assuming that the SNP(s) driving the T2D genetic association manifest an effect in hESC-β cells, the 23 SNPs identified from these combined analyses represent strong candidates for being the causal SNP at these 23 T2D association signals where *INS^GFP/w^* MEL1 was heterozygous at all SNPs in the credible set.

**Figure 6.**
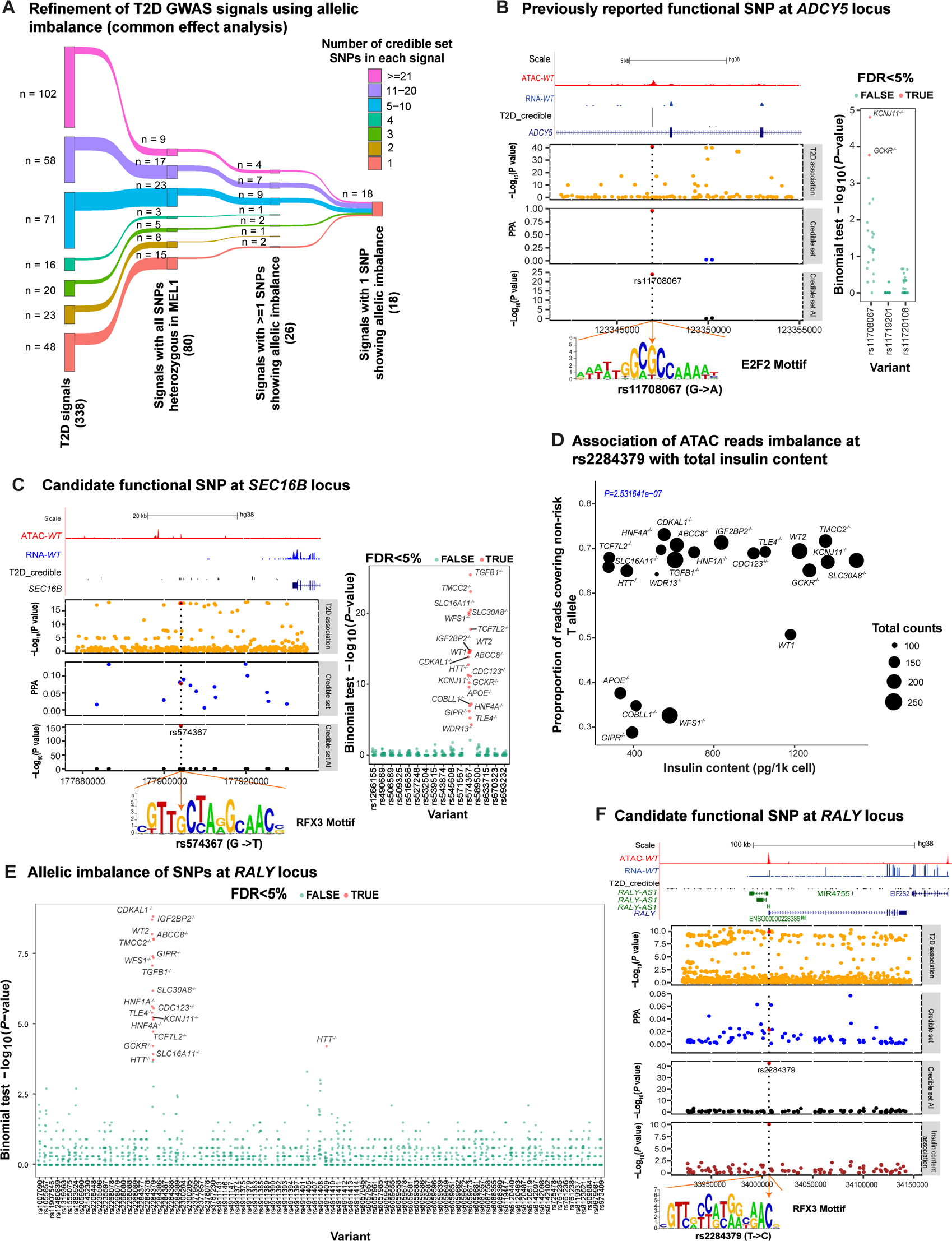
ATAC-seq allelic imbalance analysis nominates functional candidates. **(A)** Refinement of T2D GWAS signals using allelic imbalance analysis (binomial test from the common effect analysis). The *INS^GFP/w^* MEL1 hESC line is heterozygous at 80 credible set SNPs for 338 published T2D association signals^5^. Within this group of 80 SNPs, we identified at least one SNP with allelic imbalance (FDR<5%, Methods) for 26 signals using a common effect approach across all lines. At 18/26 signals, we identified a single SNP with allelic imbalance, and thus likely to be causal SNPS driving each association signal. **(B)** Candidate causal SNP at the *ADCY5* locus reported previously ^93^. Top panel: UCSC browser of ATAC-seq (red) and RNA-seq (blue) reads around the credible set of SNPs in *INS^GFP/w^* MEL1 *WT* hESC-β cells. Other panels: −log10 (*P*-values) from T2D genetic association; Posterior probability of association (PPA) from credible set analysis; −log10(*P*-value) of ATAC-seq allelic imbalance common effect analysis across all lines at credible set SNPs. Dashed vertical blue line represents the candidate functional SNP and corresponds with the position of the disruption (G to A change) in the predicted TFBS motif (yellow arrow). Note, the T2D non-risk allele “G” of rs11708067 is associated with higher ATAC-seq reads (greater chromatin accessibility) and more similar to the consensus sequence of the E2F2 binding motif. Right panel shows the −log10(*P*-value) of the line-specific effect analysis across all credible set SNPs. **(C)** An example of novel candidate functional SNP at the *SEC16B* locus. T2D non-risk allele “G” of rs574367 is associated with higher ATAC-seq reads (greater open chromatin accessibility) and more similar to the consensus sequence of the RFX3 TFBS motif. Right panel shows the −log10(*P*-value) of the line-specific effect analysis across all credible set SNPs. **(D)** Allelic imbalance at rs2284379 is associated with total insulin content. rs2284379 is the only SNP that has allelic imbalance across lines (**Figure 6F**), and the level of imbalance is associated with the total insulin content. x-axis=total insulin content (pg/1k cells) in each knockout or wild type line, y-axis=proportion of ATAC-seq reads associated with the non-risk “T” allele at the position. Each black datapoint represents a KO or WT line and the size of the data points represents the total number of ATAC-seq reads covering the SNP position. **(E)** Allelic imbalance of SNPs at *RALY* locus: Two SNPs, rs2284379 and rs4911409, show evidence of allelic imbalance within at least one line considered individually (FDR<5%). **(F)** Overview of T2D association *P* values, 99% credible set of SNP PPAs, and the common effect allelic imbalance analysis *P* values of the credible set of SNPs at the *RALY* locus. The bottom panel shows allelic imbalance at rs2284379 is associated with total insulin content.

As an example of a signal where we identified a single candidate causal SNP, we highlight a T2D association near *ADCY5* (**Figure 6B**). Within the 99% credible set there are three SNPs, all of which are heterozygous in the *INS^GFP/w^* MEL1 cell line. In our data, we found that rs11708067 exhibited allelic imbalance both in the common effect analysis and in the line-specific effect analysis (FDR<5%; *GCKR^−/−^,* and *KCNJ11^−/−^)*—where the “G” allele, associated with reduced T2D risk, shows increased accessibility. These results comport with a previous study that reports increased H3K27ac ChIP-seq reads from the “G” allele in human islets and increased luciferase activity of the “G” allele in a mouse β cell line.^48^ We performed TF footprint analysis (Methods) and discovered that the rs11708067 overlapped a E2F2 footprint where the “G” allele is predicted to have increased binding. We looked for, but did not find, an association between the chromatin accessibility of the region overlapping rs11708067 and the expression of nearby genes (FDR>5%), making the candidate effector gene at this signal an open question. Nonetheless, these results suggest that the T2D risk allele “A” may contribute to T2D risk by disrupting E2F2 binding.

As another example, we examined the complete credible set of 16 SNPs at a T2D association ∼25kb downstream of *SEC16B* (chr1:177889025), all of which were heterozygous in the *INS^GFP/w^* MEL1 cell line. Of the credible set SNPs, only rs574367 showed allelic imbalance in both common and line-specific effect e analyses (FDR<5%), with an increased proportion of reads from the non-risk “G” allele in 21 lines (**Figure 6C**). We performed TF footprint analysis and found that rs574367 strongly disrupts a predicted binding site for the RFX TF family, previously reported as an important, T2D-relevant regulator of islet gene expression.^42^ We tested for an association between chromatin accessibility of the region overlapping rs11708067 and the expression of nearby genes, but found no association (FDR>5%). A possible mechanism at this signal is that the non-risk “G” allele of rs574367 increases the binding of an RFX family TF; further work will be required to identify the downstream effector gene.

Finally, due to the intra-sample control inherent to allelic imbalance signals, associations between allelic imbalance and a phenotype of interest (e.g., a cellular trait like insulin content or disease status) offers a powerful approach to dissect regulatory effects relevant to phenotype of interest.^49–51^ To identify such effects, we jointly modeled allelic imbalance across all lines, testing for an association with each cellular trait at all T2D 99% credible set SNPs (Methods). We identified two associations (FDR<5%), both with insulin content: rs2284379 at the chr20:3267496 T2D association signal (*RALY)* and rs1800900 at the chr20:57387352 T2D association signal (**Figure 6D; Table S5**).

For one of these signals, located at chr20:32674967, all 95 SNPs in the 99% credible set were heterozygous in the parental *INS^GFP/w^*MEL1 line. In our allelic imbalance analysis, we found that two SNPs, rs2284379 and rs4911408, show evidence of allelic imbalance from the common effect analysis (FDR<5%) and two SNPs, rs2284379 and rs4911409, show evidence of allelic imbalance within at least one line considered individually (FDR<5%, **Figure 6E**). However, when considering the association between allelic imbalance across all lines and total insulin content, only one SNP—rs2284379—shows an association (**Figure 6F**). This association is particularly remarkable, in that the allelic association favors the T allele in the majority of lines, but the C allele in the four lines with the lowest insulin content **(Figure 6D, 6F**). rs2284379 occurs in the first intron of *RALY* and the T2D risk allele, “C”, is predicted to better match the binding site motifs at footprints of RFX3, ZNF737, and MTF1. Given the location of this SNP, we tested for an association between chromatin accessibility of the region overlapping this SNP and *RALY* expression but did not find an association (FDR>5%; Methods). Nonetheless, these data reduce the candidate causal SNP search space by ∼97%, to three SNPs, one of which may perturb total insulin content in hESC-β cells.

## Discussion

We developed an isogenic KO hESC-derived β cell platform to assess the molecular and cellular changes of human β cells carrying LoF mutations of 20 T2D associated genes with strong evidence of being causal. Using this platform, we profiled the effect of each individual gene in β cell differentiation, insulin production, insulin secretion in response to glucose and KCI stimulation, and β cell survival after a lipotoxic exposure, providing new insights on their roles in T2D development. Specifically, compared with *WT* hESCs, 19 KO hESCs showed impairments in at least one of the five β cell traits. Deficiency of *COBLL1*, *GIPR*, *HNF4A*, *TCF7L2*, *TGFB1*, and *TLE4* impaired β cell differentiation (**Figure 1B, 1C** and **Figure S2B**). Previous studies demonstrated mutations in *HNF4A* and *TCF7L2* have an impact on pancreas β-cell development and function,^52–55^ and these hESC experiments demonstrate a similar effect in an *in vitro* system. Similarly, dysregulation of *TGFB1* signaling has also been implicated in β cell development, proliferation, and dedifferentiation.^56^ The mechanisms underlying the relationship of T2D between *COBLL1*, *GIPR* and *TLE4* are not fully understood. Our study suggests that these genes may play a role in diabetes progression through their impact on β cell development.

While we expect some of these genes may affect a cellular phenotype in a similar way due to their transcriptional co-regulation (i.e., *HNF1A* and *HNF4A*, **Figure 1**), we also observed multiple other genes of diverse classes affecting the similar set of cellular traits. For example, loss of *ABCC8, HNF1A, WDR13,* and *HTT* does not affect β cell generation in this system, but impairs four cellular traits including total insulin content, insulin secretion response to glucose or KCl stimulation, and susceptibility to lipotoxicity. Mutations in the genes *ABCC8*^57^, *HNF1A*^58^, and *WDR13*^59^ have been associated with impaired β cell function and insulin secretion, leading to the development of diabetes and other metabolic disorders, such as congenital hyperinsulinism (CHI). Recent studies suggest that mutations in *ABCC8 and HNF1A,* may also contribute to β cell dedifferentiation/transdifferentiation,^58, 60^ which is an adaptive process where mature β cells lose their specialized identity under prolonged metabolic stress.^61^ We found that *ABCC8^−/−^*, *HNF1A^−/−^*, and *WDR13^−/−^* hESC-β cells exhibit decreased total insulin content (**Figure 1D**) along with decreased expression of β-cell genes, including *PAX4* and *NKX6-1* (**Figure S2C**). Meanwhile, for *ABCC8^−/−^*, *HNF1A^−/−^*, and *WDR13^−/−^*hESC-β cells we also observed an upregulation of some marker genes associated with ɑ cells (eg., *ARX, LOXL4*), **δ** cells (eg., *SST, LEPR, ETV1*), **ε** cells (eg., *GHRL, VTN*), PP cells (eg., *PPY*), ductal cells (eg.,*KRT19*, *SOX4*) and acinar cells (eg.,*PRSS1, PRSS2*), further confirming the impaired β cell identity in those KO cells (**Figure S2C**). Those results indicate β cell transdifferentiation might be a mechanism underlying the pathogenesis of diabetes caused by mutations in *ABCC8*, *HNF1A* and *WDR13*. Huntingtin (HTT) is a protein expressed in multiple tissues including pancreatic islets, but that is primarily associated with the development of Huntington’s disease when there is an expansion of its coding region triplet repeat.^62^ The link between the HTT mutation and T2D is still being investigated. Our study demonstrates that HTT is essential for proper β cell function and resistance to lipotoxicity, which is consistent with recent research suggesting that HTT may play a role in the regulation of pancreatic functions and survival.^63–66^

Among 20 T2D associated genes studied, only loss of *HNF4A* caused effects in all five cellular traits (**Figure 1I**). *HNF4A* is an important transcription factor affecting liver development^67^, islet development^53, 68^, and is essential for glucose-stimulated insulin secretion (GSIS) in mouse β cells.^69^ However, its regulatory role in β cell development has been less clear. We demonstrate that KO of *HNF4A* results in major loss of accessible chromatin regions where the HNF4A TFBSs are enriched. Furthermore, HNF4A appears to directly bind to regulatory elements that contribute to T2D risk, as we have shown for the FAIM2 signal. In contrast, *SLC30A8^−/−^* hESCs showed no impairments in any of five cellular traits in terms of pancreatic β cell generation, function, or survival. *SLC30A8* encodes ZnT8, a zinc transporter that mediates zinc uptake into insulin secretory granules in pancreatic islet β cells.^70^ Polymorphisms in *SLC30A8* are linked to T2D risk, with some variants increasing and others decreasing the risk.^70, 71^ However, the exact mechanism by which *SLC30A8* affects T2D risk is not fully understood. Recent studies using knockout models have shown contradictory results,^72–74^ while hPSC-based data mainly suggests that *SLC30A8* may not be essential or may even have a protective role in β cell function.^75, 76^ Our results support the latter notion that ZnT8 is not necessary for β cell function and survival, but further research is needed to fully understand their implications.

In this study, we selected 20 diverse genes for knockout to assess their functional consequences in an isogenic background. This heterogeneity of proposed function of the KO genes provided a novel opportunity to identify common gene expression pathways that correlate with β cell phenotypes. We focused on two phenotypes and identified 21 genes associated with total insulin content and 35 genes related to β cell survival under lipotoxic conditions. In a recent genome-wide pooled CRISPR loss-of-function screen study, Rottner et al. identified 580 genes influencing insulin content in the human β cell line EndoC-βH1.^74^ Out of the 580 screening hits, 154 genes, including *CALCOCO2*, *MAPK1* and *ARF1*, also showed correlations (FDR<5%) with total insulin content in our hESC-β cell system, but none of them overlapped with the top 21 candidate genes (FDR<5% and |effect size|>1.5, **Table S4**). This supports that there are shared insulin regulation pathways within the two human β cell models and highlights the necessity to diversify experimental strategies for optimal prioritization of effector genes. Functional assays in the EndoC-βH1 cells confirmed the regulatory roles of *CP, RNASE1, PCSK1N*, and *GSTA2* in controlling insulin production, and identified *TAGLN3* and *DHRS2* as regulators of β cell survival. *CP* and *GSTA2* encode enzymes that are involved in regulating oxidative stress in cells. Specifically, *CP* encodes for ceruloplasmin, which is involved in copper metabolism and antioxidant defense.^77^ *GSTA2* encodes for glutathione S-transferase A2, which is involved in detoxification and antioxidant processes.^78^ Of particular interest, prior investigations have reported elevated levels of serum ceruloplasmin in individuals with type 2 diabetes relative to healthy controls.^79, 80^ Additionally, reduced expression of *GSTA2* has been observed in human islets treated with palmitate and high glucose, a condition that mimics diabetic conditions.^81^ Our study provides complementary evidence that implicates the involvement of ceruloplasmin and GSTA2 in a compensatory mechanism that underlies β cell dysfunction caused by metabolic stress. *PCSK1N* is an inhibitor of PCSK1, which is the key enzyme controlling the conversion of proinsulin to insulin.^82^ Thus, upregulation of *PCSK1N* might directly affect insulin content by inhibiting the proinsulin to insulin conversion. *RNASE1* encodes an endonuclease that cleaves internal phosphodiester RNA bonds on the 3’-side of pyrimidine bases.^83^ Therefore, *RNASE1* may indirectly regulate insulin production by affecting the stability of *INS* mRNA. Regarding genes associated with apoptosis, *DHRS2* overexpression has been shown to induce apoptosis in certain cancer cells.^84, 85^ Although the mechanism of how *DHRS2* induces apoptosis is not fully understood, it may involve modulating intracellular reactive oxygen species (ROS) levels or regulating apoptosis-related signaling pathways. *TAGLN3* encodes transgelin 3, which has been shown to be involved in astrocyte inflammation.^86^ However, how transgelin 3 regulates β cell survival is still not clear. Overall, our findings suggest that *TAGLN3* and *DHRS2* may have a pro-apoptotic role in β cells, but further research is needed to fully elucidate their function and potential therapeutic implications.

Additional insights were derived by close inspection of allele-specific imbalance in ATAC-seq data at sites where the *INS^GFP/w^* MEL1 parental line was heterozygous. With 20 knockout and two wild type lines, all isogenic, the depth of coverage at an ATAC-seq peak was large enough to detect situations where one allele contributed substantially more than 50% to the reads. We reasoned that such observations likely pointed to a SNP with functional consequences, allowing the potential of narrowing down T2D risk loci harboring multiple SNPs in a credible set to the variant most likely to be causative. With this approach, we were able to pinpoint a single likely functional variant at each of 23 loci. One of them is a previously reported likely functional T2D SNP (rs11708067) near *ADCY5* locus.^87^ Another example allowed identification of a single likely causative SNP out of a credible set of 16 near the *SEC16B* locus. The success of this effort, demonstrating that deep ATAC-seq coverage can discover biologically interesting and GWAS-relevant allele-specific imbalance in a disease-relevant tissue, suggests that future work like this could be usefully done with a larger collection of iPSCs of diverse genotypes.

In summary, we developed an isogenic hESC platform to examine the impact of knocking out 20 T2D associated genes in the human β cell generation, insulin content, glucose and KCl stimulated insulin secretion, and β cell survival. New insights were derived about each of the individual knockout lines, especially *HNF4A^−/−^*, but the molecular comparison also revealed pathways involved in insulin production and apoptosis that would have been difficult to discern by other means. Future work to expand the panel to many more genes relevant to T2D, while maintaining the same standards for cellular phenotyping, is likely to be revealing. One can also readily imagine extrapolating this same platform to the analysis of any other polygenic disorder where there are GWAS signals pointing to risk loci, and relevant tissues can be differentiated from hPSCs and studied by gene expression and chromatin accessibility methods.

## Data and code availability

The RNA-seq, ATAC-seq data, and SNP array genotyping data generated during this study are available at GEO under accession no. GSE228665. Reviewer Token: gvcnyqaqphurzub

## Acknowledgments

This research was supported in part by the United States’ National Institutes of Health grant 1-ZIA-HG000024 (to F.S.C.), National Institute of Diabetes and Digestive and Kidney Diseases (NIDDK) grants (R01 DK124463, R01 DK116075-01A1, R01 DK119667-01A1, and 1U01DK127777-01 to S.C.), and the American Diabetes Association grant (9-22-PDFPM-06 to D.X.). We thank Kevin W. Currin and Cassie Robertson for helpful advice in the data analysis. We thank Efsun Arda for her gracious review and valuable feedback on this manuscript.

## Author Contributions

D.X., S.C., N.N., M.R.E., L.L.B., and F.S.C conceived the project. D.X., D.L.T., N.N., M.R.E., L.L.B., S.C., and F.S.C. designed the experiments. D.X., M.Z., T.Z., and A.Z. performed gene editing and generated isogenic hESCs. D.X. and X.T. performed differentiation and functional evaluation assays. D.X. and J.Z. performed the luciferase reporter assay, collected RNA and DNA samples, and prepared libraries for RNA-seq and ATAC-seq. J.J.V. and A.C.N.C. maintained EndoC-βH1 cells and constructed EndoC-βH1-luc cell lines. D.X., N.N., D.L.T., T.Y., N.S., C.G., A.L., E.M., A.J.S., M.R.E., L.L.B., S.C. and F.S.C. performed research. N.N., D.L.T., T.Y., N.S., C.G., H.J.T., and D.X. performed bioinformatics analyses. N.N., D.X., D.L.T., H.J.T., T.Y., N.S. C.G., L.L.B., S.C. and F.S.C. wrote the paper. S.C, and F.S.C. jointly supervised the study.

## STAR Methods

### Cell lines and culture conditions

We obtained *INS^GFP/w^* MEL-1 (RRID: CVCL_XA16) human embryonic stem cell (hESC) stocks from Dr. Ed Stanley at Monash University. All hESC studies were approved by the Tri-Institutional Embryonic Stem Cell Research Committee (ESCRO).To culture and maintain both wildtype (*WT*) and isogenic (see ***Generation of isogenic hESC lines***) hESCs, we followed a previously described protocol.^94^ We grew hESCs on Matrigel-coated plates in StemFlex medium (Thermo Fisher Scientific) supplemented with 50 µg/mL Normocin (InvivoGen), with medium changed daily and cultures passaged at 1:6-1:10 with ReLeSR (Stem Cell Technologies). We obtained EndoC-βH1 cells (RRID: CVCL_L909) from CNRS, and cultured them in DMEM containing 5.6 mM glucose, 2% BSA (Sigma-Aldrich), 50 µM 2-mercaptoethanol (Thermo Fisher Scientific), 10 mM nicotinamide (Sigma-Aldrich), 5.5 μg/ml transferrin (Sigma-Aldrich), 6.7 ng/ml selenite (Sigma-Aldrich), 100 U/ml penicillin and 100 μg/ml streptomycin. HEK293T cells (purchased from ATCC, CRL-11268) were cultured in DMEM supplemented with 10% FBS (Thermo Fisher Scientific). All cell lines were cultured at 37 °C with 5% CO2 and were tested for mycoplasma contamination every six months using MycoAlertTM PLUS Mycoplasma Detection Kit (Lonza).

### Generation of isogenic hESC lines

To create the isogenic KO hESC lines, we designed short guide RNAs (sgRNAs) targeting exons of 20 genes with varied evidence for T2D (**Table 1**; **Table S1**) using the web resources available at http://chopchop.cbu.uib.no/. We cloned them into the pSpCas9(BB)-2A-Puro (PX459) V2.0 vector (Addgene #62988) according to the instructions described in our previous publication.^95^ All KO lines were generated from INS^GFP/w^ MEL-1. Briefly, MEL-1 cells were dissociated using Accutase (Innovative Cell Technologies) and electroporated (5 × 10^5^ cells per sample) with 4 μg sgRNA-construct plasmids using Human Stem Cell Nucleofector^TM^ solution (Lonza) following manufacturer’s instructions. The cells were then seeded into 2 wells of 24-well plates and cultured in StemFlexTM medium with 10 μM Y-27632. They were switched to StemFlexTM medium with 0.5mg/ml puromycin on the next day and maintained for 2 days. After puromycin selection, hESCs were dissociated into single cells with Accutase and re-plated at a density of 5 cells/well in 96-well plates. 10 μM Y-27632 was added for the first 2 days. 10 days later, individual colonies were picked and re-plated into two wells of 96-well plates. When hESCs reached ∼90% confluence, one well of each clone was analyzed to confirm the indel information of each clone by Sanger sequencing a ∼500 bp window around the Cas9-sgRNA recognition site (**Figure S1**). For biallelic frameshift mutants, we expanded two clones (clones #1 and 2) with either homozygous indel mutations or compound heterozygous indel mutations in each target gene to perform cellular assays. We also expanded two *WT* clonal lines as *WT* controls to account for potential non-specific effects associated with the gene-targeting process.

### Directed differentiation of hESC to β cells

We differentiated hESCs into pancreatic β-like cells using a modified protocol from previous studies.^18, 19, 27^ Briefly, on day 0, we exposed cells to basal medium RPMI 1640 (Corning) supplemented with 1× glutamax (Thermo Fisher), 50 μg/mL normocin, 100 ng/mL Activin A (R&D systems), and 3 μM of CHIR99021 (Cayman Chemical) for 24 hours. On day 1, we changed the medium to basal RPMI 1640 medium supplemented with 1× glutamax, 50 μg/mL normocin, 0.2% FBS (Thermo Fisher Scientific), 100 ng/mL Activin A for 2 days, producing definitive endoderm cells. On day 3, we cultured the definitive endoderm cells in basal MCDB131 supplemented with 1× Glutamax (ThermoFisher Scientific), 1.5 g/L sodium bicarbonate (Sigma-Aldrich), 2% bovine serum albumin (BSA, Lampire),10 mM glucose (Sigma Aldrich), 50 ng/mL FGF7 (Peprotech) and 0.25 mM L-ascorbic acid (Sigma Aldrich) for 2 days to acquire primitive gut tube. On day 5, we induced the cells to differentiate to posterior foregut in basal medium MCDB 131 supplemented with 2% BSA, 2.5 g/L sodium bicarbonate, 1× Glutamax, 10 mM glucose, 0.25 mM L-ascorbic acid, 50 ng/mL FGF-7, 2 μM Retinoic acid (RA; Sigma Aldrich), 100 nM LDN193189 (LDN, Axon Medchem), 1:200 ITS-X (Thermo Fisher Scientific), 200 nM TPPB (Tocris Bioscience) and 0.25 μM SANT-1 (Sigma Aldrich) for 2 days. On day 7, we induced the cells to differentiate to pancreatic endoderm in MCDB 131 medium supplemented with 2% BSA, 2.5 g/L sodium bicarbonate, 1× Glutamax, 10 mM glucose, 0.25 mM L-ascorbic acid, 2 ng/mL of FGF-7, 0.2 μM RA, 200 nM LDN193189, 1:200 ITS-X, 100 nM TPPB and 0.25 μM SANT-1 for 3 days. On day 10, the cells were induced to differentiate to pancreatic endocrine precursors in MCDB 131 medium supplemented with 1.5 g/L sodium bicarbonate, 1×Glutamax, 20 mM glucose at final concentration, 2% BSA, 0.1 μM RA, 100 nM LDN193189, 1:200 ITS-X, 0.25 mM SANT-1, 1 μM T3 hormone (Sigma Aldrich), 10 μM ALK5 inhibitor II (Cayman Chemical), 10 μM zinc sulfate heptahydrate (Sigma Aldrich) and 10 μg/mL of heparin (Sigma Aldrich) for 3 days. On day 13, we exposed cells to MCDB 131 medium supplemented with 1.5 g/L sodium bicarbonate, 1× Glutamax, 20 mM glucose at final concentration, 2% BSA, 100 nM LDN193189, 1:200 ITS-X, 1 μM T3, 10 μM zinc sulfate, 10 μg/mL of heparin, 100 nM gamma secretase inhibitor XX (Millipore) for the 7 days. On day 21, cells were exposed to MCDB 131 medium supplemented with 1.5 g/L sodium bicarbonate, 1× Glutamax, 20 mM glucose, 2% BSA, 1:200 ITS-X, 1 μM T3, 10 μM ALK5 inhibitor II, 10 μM zinc sulfate, 10 μg/mL of heparin, 1 mM N-acetyl cysteine (Sigma Aldrich), 10 μM Trolox (Millipore), 2 μM R428 (MedchemExpress) for 7-15 days. We refreshed the medium every day. Specially, for GSIS and KSIS assay, We dissociated cells at stage 6 using accutase, and seeded them into 96-well U-bottom low attachment plates as described in **Static GSIS and KSIS Assays**. We presented the actual number of biological replicates (n) for each downstream assay in **Table S2**.

### Immunofluorescence staining and confocal microscopy

We fixed cells in 4% paraformaldehyde solution (Thermo Fisher Scientific) for 20 minutes, washed them three times in PBS with 5 minutes incubation for each wash, and blocked and permeabilized cells in a PBS solution containing 5% horse serum and 0.3% Triton X-100 (Sigma Aldrich) for 1 hour at room temperature. Then, we incubated the cells with primary antibodies overnight at 4°C and washed them in PBS with a 5-minute incubation three times. After a 1-hour incubation with fluorescence-conjugated secondary antibodies (Alexafluor, Thermo Fisher Scientific) at room temperature, we washed the cells with PBS three times. The detailed antibody information has been included as **Table S6**. Images in **Figure S2A** were taken by Inverted Microscope/Apotome (Zeiss). Images in **Figure 5K** were taken by LSM 800 confocal microscope (Zeiss) and scored using MetaMorph® image analysis software (Molecular Devices). We calculated mean ± SD for each assay using 3 independent biological replicates, and we present those data in **Figure 5L**.

### Fluorescence-activated cell sorting

We dissociated hESC-derived cells at day 24 into single cells using Accutase and resuspended in PBS supplemented with 0.5% BSA, 300 nM DAPI, and 2 mM EDTA. The Flow Cytometry Core Facility in Weill Cornell Medicine helped conduct the sorting experiments and collect GFP+DAPI-cells by BD FACS Melody™ Cell Sorter. All experiments were performed with >=3 independent replicates. For RNA-seq, we collected 500,000 cells for each replicate. For ATAC-seq or ELISA assay, we collected 50,000 cells per replicate. We present the actual number of biological replicates (n) in **Table S2**.

### Flow cytometry analysis

We dissociated hESC-derived cells or EndoC-βH1 cells using Accutase. To analyze GFP expression, we resuspended the hESC-derived cells in PBS and used them directly for analysis. The gating strategy for the analysis of GFP+ cells is shown in **Figure S2D**. For Annexin V cellular apoptosis analysis, we stained hESC-derived or EndoC-βH1 cells with the APC/Annexin V apoptosis detection Kit (BD Bioscience) and DAPI according to manufacturer’s instructions and analyzed cells using Attune NxT Flow Cytometer (Thermo Fisher Scientific) within 30 minutes. The gating strategy for the analysis of apoptotic rate in hESC-derived β cells and EndoC-βH1 cells is shown in **Figure S2D** and **Figure S5D**, respectively. All experiments were performed with >=3 independent replicates. We present the actual number of biological replicates in **Table S2** and the legends of **Figure 5H** and **5J**.

### Static GSIS and KSIS Assays

We dissociated cells at stage 6 using Accutase and resuspended them in S6 medium supplemented with 10 μM Y-27632 at a final concentration of 300 cells/μl. Using a multichannel pipette and trough, we filled 96-well U-bottom low attachment plates with 100 cell suspensions in each well and spun at 300g for 5 minutes. Cells were aggregated into clusters by incubating for at least 24 hours at 37°C with 5% CO2 and then fed every 48 hours until at least the 8th day at Stage 7. Before the static GSIS/KSIS assays, 6-8 islet-like clusters were combined into one well as one replicate and starved in S7 medium but with 5 mM glucose for 12 hours. Subsequently we removed the medium and washed cell clusters with fresh KRBH Buffer. We then incubated the cells in LG KRBH (with 0.1% BSA and 2 mM glucose) for 1 hour in an air incubator at 37°C. We aspirated the media and replaced it with 200 μL LG KRBH buffer or LG KRBH buffer with combinations of 20 mM glucose, or 30 mM KCl to each well and incubated at 37°C for 1 hour. Plates were spun and the top 120 µl supernatants were collected. The residual medium was removed, and cell clusters of each well were lysed by RIPA buffer (Sigma Aldrich) supplemented with Protease and Phosphatase Inhibitor Cocktail (Thermo Fisher Scientific). We measured insulin content in both supernatant and cell lysis using STELLUX Chemi Human Insulin ELISA Jumbo kit (Alpco). We calculated mean ± SD for each assay using 6-8 independent biological replicates, and we present the actual number of biological replicates in **Table S2**.

### Luciferase Reporter Assay

We based the construction of all luciferase vectors on the pGL4.23[luc2/minP] vector (Promega) which contains a firefly luciferase gene *luc2* under regulation of a TATA-box minimal promoter (minP). From genomic DNA of EndoC-βH1 cells, we cloned the DNA region (723bp, from chr12:49868906 to chr12:49869637) at the locus of T2D_fmap. FAIM2.chr12:50263148. This construct included the “A” allele at rs7132908. We then subcloned this into pGL4.23[luc2/minP] vector (Promega) between the XhoI and Bgl II restriction sites. Using PCR amplification with mutated primers, followed by DpnI digestion and nick ligation in *E. coli* ^96^, we performed site-directed mutation of the plasmid to produce the same vector with the “G” allele at rs7132908. Constructs of all plasmids were confirmed by Sanger sequencing. The sequences of primers used to construct and validate each vector are shown in **Table S6**. For luciferase assays, we seeded EndoC-βH1 cells into 12-well plates at a density of 5.0 × 10^5^ cells/well, cultured those for 48 hours, and then transfected with firefly luciferase reporter vectors. We used a Renilla luciferase vector carrying the SV40 promoter, phRL-SV40 (Promega) as an internal control. We co-transfected cells with firefly luciferase reporters (1 µg/well) and phRL-SV40 (20 ng/well), using Lipofectamine 2000 (Thermo Fisher Scientific), following the manufacturer’s instructions. Transfections were performed in triplicate for experimental group using constructed vectors and in quadruplicate for control group using empty vector. We harvested cells at 48 hours after transfection and lysed them in the passive lysis buffer (Promega). We measured luciferase activity of the lysates with the Dual-Luciferase® Reporter Assay System (Promega) according to the manufacturer’s protocols. We calculated the ratio of firefly/Renilla luciferase activity for each tested enhancer candidate vector and normalized that to the empty vector pGL4.23[luc2/minP] as the final relative luciferase intensity. We calculated mean ± SD for each assay using 3-4 independent biological replicates and we present those data in **Figure 3H**.

### CRISPR perturbation experiments

To perturb the transcriptional expression of candidate genes, we designed two different sgRNAs for each candidate gene, using the web resources available at http://chopchop.cbu.uib.no/. We cloned sgRNAs targeting *RNASE1*, *PCSK1N*, *GSTA2*, *TAGLN3*, *ADCYAP1* and *DHRS2* (sequences of sgRNA targeting regions are listed in **Table S1**) into dSpCas9-VPR vector (Addgene #139090) for gene activation. We cloned sgRNAs targeting *CP*, *FOSB*, and *SYNPO* into the dCas9-KRAB vector (Addgene #139097) according to the previously described instructions ^97^. We produced lentivirus expressing each CRISPRa or CRISPRa system in HEK293T cells, using a second-generation viral packaging system, and used the virus to infect EndoC-βH1 cells or EndoC-luc cells as previously described ^44^. At 48 hours post transduction, we treated cells with 2 µg/mL puromycin for one week to select for infected cells, which were then used for downstream functional assays.

### Generation of EndoC-βH1-luc cells and Nano-luc reporter assay

We produced lentivirus expressing proinsulin-luciferase fusion protein in 10-cm diameter dishes from 80% confluent HEK293T cells, transfected with lentiviral packaging plasmid psPAX2 (Addgene #12260), envelope plasmid pMD2.G (Addgene #12259) and Proinsulin-NanoLuc plasmid (Addgene #62057). We pooled viral supernatant harvested at 48h and 72h post-transfection and concentrated it using Lenti-X Concentrator (Takara) according to the instructions. We added the virus prep to EndoC-βH1 cells in fresh culture medium (see **Cell lines and culture conditions**) with 8 μg/ml Polybrene (Sigma-Aldrich), and spun the cells at 800 x g for 1 hour at 30 °C. After 24 hours in the presence of virus, we placed cells in fresh growth media. Subsequently, we treated the infected EndoC-βH1 cells with 5 µg/mL blasticidin (Thermo Fisher Scientific) for one week to produce the stable EndoC-luc lines. To test if *CP*, *RNASE1*, *PCSK1N* and *GSTA2* have effects on insulin translation/processing, we conducted CRISPR perturbation experiments in EndoC-luc cells (see **CRISPR perturbation experiments**). We dissociated EndoC-luc cells into single cells and counted them by a Countess II Cell Counter (Thermo Fisher Scientific). 10,000 EndoC-βH1-luc cells were lysed in 100 μl passive lysis buffer (Promega) and we then measured intracellular Nano-luc intensity of lysate with the Nano-Glo® Luciferase Assay System (Promega) according to the manufacturer’s protocols. We calculated mean ± SD for each assay using 3 independent biological replicates and we present those data in **Figure 4K-4L**.

### qRT-PCR

We isolated total RNA from EndoC-βH1 cells or EndoC-luc cells using the RNeasy Plus Mini Kit (QIAGEN), quantified RNA with a NanoDrop spectrophotometer (Thermo Fisher Scientific), and synthesized cDNA with a high-capacity reverse transcription kit (Thermo Fisher Scientific). We performed real-time qPCR with a LightCycler 480 (Roche) instrument with LightCycler DNA master SYBR Green I reagents (Roche). Primer sequences specific to *INS*, candidate genes being tested, and the reference gene (*GAPDH*) are listed in **Table S1 and S6**. We determined Delta-delta-cycle threshold (DDCT) relative to the *GAPDH* and control samples. We calculated mean ± SD for each assay using 3 independent biological replicates, and we present those data in **Figure 4I-J** and **Figure S5A-C**.

### *INS^GFP/w^* MEL1 genotyping, quality control, and imputation

We genotyped the parental *INS^GFP/w^* MEL1 hESC line used for generating the isogenic hESC lines using the Infinium Omni2.5Exome-8 BeadChip array v1.3 (Illumina, San Diego, CA) at the NHGRI Genomics Core facility, resulting in a call rate of 99.7% (out of 2,612,357 SNPs). Using novoalign v2.07.11 (http://www.novocraft.com/products/novoalign), we mapped the array probe sequences to the GRCh37 (hg19) genome assembly and filtered variants with ambiguous probe alignments as previously described in Currin et al..^93^ We combined the *INS^GFP/w^* MEL1 genotypes with 15 samples genotyped on the same chip and 2,504 samples from 1000G project phase 3 release ^98^. We removed variants not in the 1000G Phase 3 release panel, with missing genotypes in >1 of the 16 genotyped samples, that are likely palindromic variants with MAF>0.4 in the 16 genotyped samples, or with a genotype distribution that deviates from Hardy-Weinberg equilibrium (*P*-value<1×10^−4^). After filtering the genotypes, we used the remaining 1,589,371 SNPs for genotype imputation on the Michigan TOPmed Server (Minimac v4;^99^). In total, we generated imputed genotypes of all SNPs (r2>0.3) included in the TOPmed panel for the analysis described in **ATAC-seq allelic imbalance analysis**.

### RNA isolation, sequencing, and processing

For the 20 KO and 2 *WT* hESC lines described in ***Generation of isogenic hESC lines***, we selected and expanded clones (see ***hESC procurement, culture, and maintenance***), differentiated hESCs into hESC-β cells (see ***β-cell differentiation protocol of hESCs***), and generated RNA-seq data on the purified hESC-β cells (**Table S7**). We selected a single clone for the KO lines and two clones for the *WT* lines (**Table S2**). For each clone, we performed the differentiation and RNA-seq experiment in 3-4 replicates (**Table S2**). For each replicate, we extracted and purified total RNA from the FACS-sorted hESCs-β cells (see **Table S2**) using the Absolutely RNA Nanoprep kit (Agilent Technologies), quantified with a NanoDrop spectrophotometer (Thermo Fisher Scientific). We used the Weill Cornell Genomics Core to sequence the purified RNA. Briefly, we evaluated the quality of RNA samples using the Agilent bioanalyzer (Agilent Technologies), generated cDNA libraries using TruSeq RNA Sample Preparation (Illumina) and sequenced the cDNA libraries using an Illumina NovaSeq 6000 with 2×51 bp cycles (Illumina). We aligned the processed reads to the GRCh38 genome assembly using STAR v2.73a ^100^ with default parameters and quantified expression levels of Gencode v19 genes (Ensembl release 103) using QoRTs (v1.3.6,^101^; **Table S7**). Finally, we generated a raw read count matrix of gene by library and a normalized mRNA expression matrix of transcripts per million (TPM). On average, we generated 36,586,780 (13,273,689-123,518,245) paired-end reads per library, of which 84.36% uniquely aligned to the genome. Out of the aligned reads, 79.42% were unambiguously assigned to unique genes, emphasizing the quality of these data. For sequencing statistics, see **Table S7**.

### RNA-seq quality control

To assess the reproducibility of RNA-seq data from replicate libraries, we normalized the gene expression using log_2_(counts per million total reads [CPM]) and calculated Pearson correlation of pair-wise replicate libraries. We did not identify any outlier (minimum Pearson’s *r*≥0.95) and used the resulting data for downstream analysis.

To assess the contamination of the total RNA isolated for RNA-seq, we combined the aligned reads across replicate libraries and flagged duplicate reads in the bam files using GATK v4.1.9.0 MarkDuplicates^102^ with default options. Using the “view” function from Samtools v1.9^103, 104^ with option “-F 3840”, we removed duplicate reads as well as those reads that mapped to supplementary/secondary alignments. Finally, using the *INS^GFP/w^* MEL1 genotypes (see ***INS^GFP/w^ MEL1 genotyping, quality control, and imputation***) as the reference panel, we used verifybamID v1.1.1^105^ with options --ignoreRG --precise --self --maxDepth 100 to identify clones with RNA that was likely contaminated (FREEMIX>5%) or did not match reference genotypes (CHIPMIX>5%). We did not identify any problematic clones.

### ATAC-seq library preparation, sequencing, and processing

We used the FACS-sorted hESC-β cells described in ***RNA isolation, sequencing, and processing*** to perform ATAC-seq and prepared samples according to Weill Cornell Medicine Epigenetics Core facility protocol (https://epicore.med.cornell.edu/services.php?option=atacseqdescription#seq). Briefly, we sorted 50,000 INS-GFP+ cells, washed them with 1000 μl of ice-cold PBS, and resuspended the pellets in 25 μl of ice cold 1X ATAC Buffer [20mM Tris-HCl (pH 7.4), 20mM NaCl and 6mM MgCl2]. We incubated the samples for 5 minutes on ice, thoroughly mixed in 25 μl of ice cold ATAC-Detergent-buffer [20mM Tris-HCl (pH 7.4), 20 mM NaCl and 6 mM MgCl2, 0.2% Igepal CA-630 (Sigma Aldrich), 0.2% Tween 20 (Sigma Aldrich) and 0.02% Digitonin (Promega), and continued incubating the samples on ice for another 3 minutes. After incubation, we centrifuged the samples and collected the pellets. Next, we resuspended the pellets in the following transposase mixture: 25 μl 2X TD Buffer (Illumina), 2.5 μl TDE1 (Illumina), 16.5 ul PBS, 0.5 ul Digitonin (1%), 0.5 ul Tween-20 (10%), and 5 ul H_2_O. We incubated the suspended cells at 37°C for 30 minutes in a thermomixer (Benchmark) set to 500 rpm. We added 250 μl of Zymo DNA binding buffer to the suspension and purified the tagmented DNA with Zymo DNA clean and concentrator (Zymo research) according to manufacturer’s instructions. We submitted the samples to the Weill Cornell Medicine Epigenetics Core facility for library preparation according to a previously published method^106^ and NovaSeq SP (800M reads) 2×50 cycles (PE50) sequencing. We trimmed adaptor sequences using CTA (v0.1.2) and aligned the trimmed reads to the GRCh38 genome assembly using BWA-MEM v0.7.17-r1194 with the -M option.^107^ On average, we generated 94,307,193 (70,146,700-111,259,636) reads per library, of which 96.84% aligned to the genome as primary alignments. After removing duplicate reads with GATK v4.1.9.0 MarkDuplicates and filtering for autosomal, properly paired reads with mapping quality ≥30 with samtools v1.9^103, 104^, we retained 63,255,116 uniquely aligned primary reads per library (minimum of 47,329,535) for downstream analyses (**Table S8**).

Using the filtered reads, we called ATAC peaks as described in Rai et al..^108^ Briefly, we converted the aligned BAM files to BED files using the bamtobed function from bedtools v2.26.0^109^ and called peaks using MACS2 v2.2.7.1 ^110^ with options “--nomodel --shift -100 --extsize 200 -B --keep-dup all”, removing candidate peaks that overlap with ENCODE blacklists^111^ and controlling for a false discovery rate (FDR) of 5%. For each hESC-β line (20 KO and 2 *WT*), we merged peaks across replicates and retained peaks present in ≥2 replicates. Next, we created a master set of peaks by merging peaks across all 22 lines, generating 208,945 peaks. Finally, we used this master set of peaks to quantify the number of reads mapping to the peaks within each library using the “multicov” function from bedtools v with the option “-q 30” and the aligned BAM files, generating the accessible chromatin region count matrix.

### ATAC-seq quality control

We used FastQC and MultiQC to generate and aggregate QC metrics across libraries. We evaluated the base quality scores (“per_base_sequence_quality_scores”) and sequence quality scores (“per_sequence_quality_scores”), and detected all samples passed MultiQC thresholds with “pass”. To assess reproducibility across hESC-β line replicates, we normalized the peak count matrix (see ***ATAC-seq library preparation, sequencing, and processing***) using log_2_(CPM) and calculated the pairwise Pearson correlation of the normalized counts between replicates for each hESC-β line. We identified no outlier libraries (Pearson correlation coefficients ≥0.97). Finally, we identified potential contamination or sample swaps in the isogenic hESC lines by combining the filtered, aligned reads (see ***ATAC-seq library preparation, sequencing, and processing***) across replicates (merge function from Samtools v1.9 ^103, 104^) and using verifybamID v1.1.1^105^ with options “--ignoreRG --precise --self --maxDepth 100” on the merged reads and *INS^GFP/w^* MEL1 genotypes (as the reference panel). We identified no contamination (all lines with FREEMIX<5%) or sample swaps (all lines with CHIPMIX<3%).

### Identification of differentially expressed genes and differentially accessible regions in KO hESC-β cells compared to *WT* cells

We tested for differential expression and accessibility in the KO hESC-β cells compared to the *WT* cells (see **Table S2**) using DESeq2 v1.32.0.^112^ For each set of comparisons, we retained features (e.g., genes or chromatin regions) with CPM≥0.5 in ≥50% of the replicate libraries across the KO and *WT* lines (>=3). To identify shared differences between the KO line and both WT lines, we used the Wald test implemented in DESeq2 with default options to compare the KO line against each *WT* line and meta-analyzed the results using the rem_mv function from MetaVolcano v1.10.0^113^ with default parameters. We performed multiple hypothesis correction using the Benjamini-Hochberg procedure (BH;^114^) and considered features with |fold change (FC)|>1.5 and FDR<5% to be differentially expressed or accessible.

To test for differential expression, we used the gene expression matrix described in ***RNA isolation, sequencing, and processing***. To test for differential chromatin accessibility, we used the accessible chromatin region count matrix described in ***ATAC-seq library preparation, sequencing, and processing***.

### T2D effector genes

We downloaded a list of 257 T2D predicted effector genes generated by integrating the results from three different approaches, namely, “Curated T2D effector gene prediction”, “Effector index predictions”, and “Integrated classifier predictions” (https://t2d.hugeamp.org/method.html?trait=t2d&dataset=egls, accessed October 1, 2022) and considered them as the “T2D effector genes”.

### Enrichment of gene sets in KO hESC-β cells compared to *WT* cells

We tested for gene sets enriched in the KO hESC-β cells compared to *WT* cells using fgsea v1.20.0.^115^ Using the differential expression results from each KO line (see ***Identifying differentially expressed genes and differentially accessible regions in KO hESC-β cells compared to WT cells***), we ranked genes by the meta-analysis log_2_(FC) in descending order. We performed gene set enrichment analysis using fgsea with default parameters, the ranked gene list, and gene sets from the Kyoto Encyclopedia of Genes and Genomes (KEGG) database (obtained from the R package msigdbr v7.5.1^116^). We performed multiple hypothesis correction using the BH procedure ^114^ and considered gene sets with FDR<5% as enriched.

### β-cell gene expression specificity scores

We used a single cell RNA-seq dataset of 12 libraries prepared from human pancreatic islets of one donor^37^ to assess islet cell type specific expression of genes. We collected sequence reads of 16,028 cells representing major endocrine and exocrine cell types. We normalized the raw read counts by library size and transcript length for each cell and generated a TPM matrix of cell barcode by gene. Next, we applied CELLEX v1.2.2^117^ to derive cell type specificity scores using the TPM matrix and cell type labels.

### Enrichment of differentially expressed genes in β cell specific genes

We binned protein coding genes into 10 groups with equal number of genes where we have β cell expression specificity scores (>0) using the single cell RNA-seq data from^37^ (see **β-cell gene expression specificity scores**). This approach defines the genes in bin 1 to be highly specific to β cells and those in bin 10 the least specific. We included genes that are expressed ubiquitously (specificity score = 0) into the bin 10. Next, we tested enrichment of differentially expressed genes in each bin per KO line using Fisher’s exact test (fisher.test function of R stats package; v4.1.2). We performed multiple hypothesis correction using the BH procedure^114^ per KO line and considered gene sets with FDR<5% as enriched.

### Association between chromatin accessibility/gene expression and cellular traits

To identify associations with insulin content, apoptotic rate, differentiation efficiency, glucose-induced insulin secretion (GSIS), and KCl-induced insulin secretion (KSIS), we performed differential chromatin accessibility and gene expression analysis across all samples. Since sequencing replicates and phenotypic assay replicates were not paired, we summed the feature (i.e., accessible chromatin regions or genes) reads and averaged the cellular assay results across replicates. We standardized the cellular trait values, removed features with low signal—keeping accessible regions and genes with CPM ≥0.5 in ≥50% of samples—and used DESeq2 v1.32.0^112^ to test for an association between each feature and cellular trait. For each omics feature type and cellular trait pair, we removed tests where the regression was driven by an outlier(s) (minimum Cook’s distance *P*-value<0.01^118^), used the BH procedure^114^ to control for the number of tests, and considered tests with |fold change (FC)|>1.5 and FDR<5% to be associated.

### Enrichment of differentially expressed genes at differentially accessible regions

For each KO line, we tested for enrichment of DEGs nearby DARs (see ***Identifying differentially expressed genes and differentially accessible regions in KO hESC-β cells compared to WT cells***). Briefly, we performed a Fisher’s exact test (fisher.test function of R stats package; v4.1.2) to evaluate the enrichment of DARs out of all ATAC-seq peaks that overlap with DARs where the transcription start site (TSS) is within a specified window size. For TSSs, we used the genomic coordinates defined in the NCBI RefSeq release (NCBI RefSeq table; GRCh38 assembly) from UCSC Genome Browser (https://genome.ucsc.edu/)119. We tested using 5kb, 10kb, 25kb, 50kb, and 100kb window sizes. For each window size analysis, we applied the Benjamini-Hochberg procedure^114^ to correct for multiple hypotheses testing across all KO lines. We considered DARs to be enriched around DEGs at 5% FDR.

### Association of gene expression and chromatin accessibility

Using the paired RNA-seq and ATAC-seq data for each hESC-β replicate, we tested for associations between gene expression and chromatin accessibility. We removed any gene or ATAC peak with CPM≤0.5 in ≥50% of all 67 libraries to focus on shared features across all KO lines and considered gene-peak pairs where the peak was within 50kb of either side of the gene TSS, which was derived from NCBI RefSeq release (NCBI RefSeq table; GRCh38 assembly) from UCSC Genome Browser (https://genome.ucsc.edu/).119 For each gene-peak pair, we used the qtl_test_lmm function from LIMIX v1.0.17^120^ to fit a linear regression to model the inverse-normalized peak counts as the dependent variable and the inverse-normalized gene expression as the independent variable. We controlled for the FDR across all gene-peak tests using the BH procedure.^114^ To explore the effects of different window sizes, we also tested for associations using 5kb, 25kb, and 100kb windows around the gene TSSs to identify gene-peak pairs. For the gene expression and chromatin accessibility data, we used the counts matrices described in **Association between chromatin accessibility/gene expression and cellular trait**.

### Effects of T2D GWAS credible set of SNPs on transcription factor (TF) footprints

In order to assess the effect of SNPs on regulatory elements, we performed a TF footprint analysis in the merged ATAC-seq peaks for each cell line individually. We scanned the peak regions with position weight matrices (PWMs) of the directly determined TF motifs included in Cis-BP v2^121^ using “Find Individual Motif Occurrences” (FIMO) v5.4.1 ^122^ with default options. Next, we used CENTIPEDE v1.2^123^ to call footprints for each FIMO scan result in combination with the corresponding ATAC-seq aligned bam file. This approach allowed us to measure the number of transposase Tn5 integration events at a region ±100bp from each motif occurrence. We defined a motif occurrence to be bound by the respective TF if the CENTIPEDE posterior probability was ≥0.95 and its coordinates were fully contained within an ATAC-seq peak. We further considered any T2D 99% credible set SNP^5^ overlapping such a motif occurrence to be potentially disrupting the binding site of the respective TF.

### Enrichment of differential ATAC-seq peaks in ChromHMM

To investigate the enrichment of DARs in islet regulatory regions, we analyzed the regulatory features defined by ChromHMM^42^. Using the intersect function of bedtools (v2.26.0), we compared DARs identified in “**Identification of differentially expressed genes and differentially accessible regions in KO hESC-β cells compared to WT cells**” with each ChromHMM feature, with a restriction of DARs that overlapped at least 50% with a feature of interest (using option “-f 0.5”). We conducted Fishers’ exact test (fisher.test function of R stats package; v4.1.2) to evaluate the enrichment of suppressed DARs among all DARs that overlap with a feature. We also performed the same test in the other way around - for the enrichment of activated DARs for the set of features. We applied the Benjamini-Hochberg procedure^114^ to correct for multiple hypotheses across all 14 ChromHMM features. We considered a feature of interest in a line to have enriched suppressed/activated DARs at 5% FDR.

### Enrichment of TFBSs in differential ATAC-seq peaks

We tested for enrichment of TF binding site motifs for 677 TFs that are expressed in *WT* line (TPM>0) in the suppressed or activated ATAC-seq peaks in each KO and *WT* line (see **Effects of T2D GWAS credible set of SNPs on transcription factor (TF) footprint and Comparison of chromatin accessibility in KO and *WT* lines)** using “Simple Enrichment Analysis” (SEA) with default options (v5.4.1)^124^. We performed multiple hypothesis correction using the Benjamini-Hochberg procedure^114^ and considered motifs with FDR<5% to be enriched.

### ATAC-seq allelic imbalance analysis

For each line, we filtered duplicate reads, reads identified as secondary alignments, or reads with poor mapping quality (<30) and merged the aligned, paired reads across replicates using the “merge” function from samtools v1.9.^103, 104^ We applied WASP v0.3.4^125^ to quantify allele counts while controlling for mapping biases at heterozygous variants in the *INS^GFP/w^* MEL1 parental line, using only variants with an imputation quality r^2^>0.3. From the allelic counts generated by WASP, we selected T2D 99% credible sets SNPs from Mahajan et al.^5^ and performed a two-sided binomial test (binom.test function in R v4.2.2) in each line across all variants with >=1 total counts. To identify common effects across lines, we performed a meta-analysis using Stouffer’s Z-score method,^126, 127^ weighting the Z-scores from each line by the total read counts overlapping the variant (sumz method from metap R package v1.8). For both line-specific and common effect analyses, we controlled for the number of tests using the BH procedure.

### Association between ATAC-seq allelic imbalance signals with cellular traits

Using the allelic counts generated by WASP (see **ATAC-seq allelic imbalance analysis**), we selected T2D 99% credible sets SNPs from Mahajan et al.^5^ and fit a binomial regression (sm.GLM function with family set to sm.families.Binomial() from the statsmodels Python package v0.13.2) across all lines, testing for an association between allelic imbalance and each cellular trait. For each variant considered for allelic imbalance, we standardized the cellular trait values prior to fitting the regression. We included all variants in the analysis with >=1 count. Finally, we removed associations driven by outliers, dropping those with a minimum Cook’s distance *P*-value<0.01,^118^ and controlled for the number of tests for each cellular trait using the BH procedure.

### Statistical analysis for functional assay

All experiments were performed with >=3 independent replicates unless otherwise specified in the Figure legends. Unless otherwise noted in the rest of the method sections, for comparisons of functional assay results, we calculated mean ± SD for each assay using >=3 independent biological replicates. We included descriptions of each statistical test and the n and P values in each Figure legend and related experimental method sections.

## Supplemental Figures

**Figure S1:** DNA sequence confirmation of frameshift mutations in isogenic KO hESC clones (two clones for each KO line) compared two clones of WT hESCs.

**Figure S2:** Functional characterization of isogenic *WT* and KO hESCs and hESC-β cells.

**Figure S3:** ATAC-seq and RNA-seq of isogenic *WT* and KO hESC-β cells.

**Figure S4:** Characterization of ATAC-seq and RNA-seq of *HNF4A*^−/−^ hESC-β cells.

**Figure S5:** Functional evaluation of candidate genes that are potentially associated with insulin content or β-cell survival.

**Figure S6:** Refinement of T2D GWAS signals using allelic imbalance analysis (binomial test from the line-specific analysis).

## Supplemental Tables

**Table S1:** Sequences of sgRNA targeting regions and primers used in gene KO and perturbation.

**Table S2:** Number of biological replicates used for molecular sequencing and cellular trait functional assays.

**Table S3:** 99% Likely functional T2D credible set of SNPs that disrupt HNF4A binding sites and overlap suppressed ATAC-seq peaks in *HNF4A^−/−^*hESC-β cells.

**Table S4:** Genes associated with insulin content and apoptotic rate across 22 cell lines.

**Table S5:** Summary of ATAC-seq allelic imbalance signatures at heterozygous SNPs in 99% credible sets for T2D association.

**Table S6:** Summary of antibodies and other primers related to STAR methods.

**Table S7:** Summary of RNA-seq sequencing statistics.

**Table S8:** Summary of ATAC-seq sequencing statistics.

