## Supplementary material for "Functional interrogation of twenty type 2 diabetes-associated genes using isogenic hESC-derived β-like cells": Figure S1-S6 and Table S1-S8

### Supplemental information

#### Supplemental Figures

**Figure S1:** DNA sequence confirmation of frameshift mutations in isogenic KO hESC clones (two clones for each KO line) compared two clones of WT hESCs.

**Figure S2:** Functional characterization of isogenic *WT* and KO hESCs and hESC-β cells.

**Figure S3:** ATAC-seq and RNA-seq of isogenic *WT* and KO hESC-β cells.

**Figure S4:** Characterization of ATAC-seq and RNA-seq of *HNF4A*$^{-/-}$ hESC-β cells.

**Figure S5:** Functional evaluation of candidate genes that are potentially associated with insulin content or β-cell survival.

**Figure S6:** Refinement of T2D GWAS signals using allelic imbalance analysis (binomial test from the line-specific analysis).

#### Supplemental Tables

**Table S1:** Sequences of sgRNA targeting regions and primers used in gene KO and perturbation.

**Table S2:** Number of biological replicates used for molecular sequencing and cellular trait functional assays.

**Table S3:** 99% Likely functional T2D credible set of SNPs that disrupt HNF4A binding sites and overlap suppressed ATAC-seq peaks in *HNF4A*$^{-/-}$ hESC-β cells.

**Table S4:** Genes associated with insulin content and apoptotic rate across 22 cell lines.

**Table S5:** Summary of ATAC-seq allelic imbalance signatures at heterozygous SNPs in 99% credible sets for T2D association.

**Table S6:** Summary of antibodies and other primers related to STAR methods.

**Table S7:** Summary of RNA-seq sequencing statistics.

**Table S8:** Summary of ATAC-seq sequencing statistics.

### Figure S1

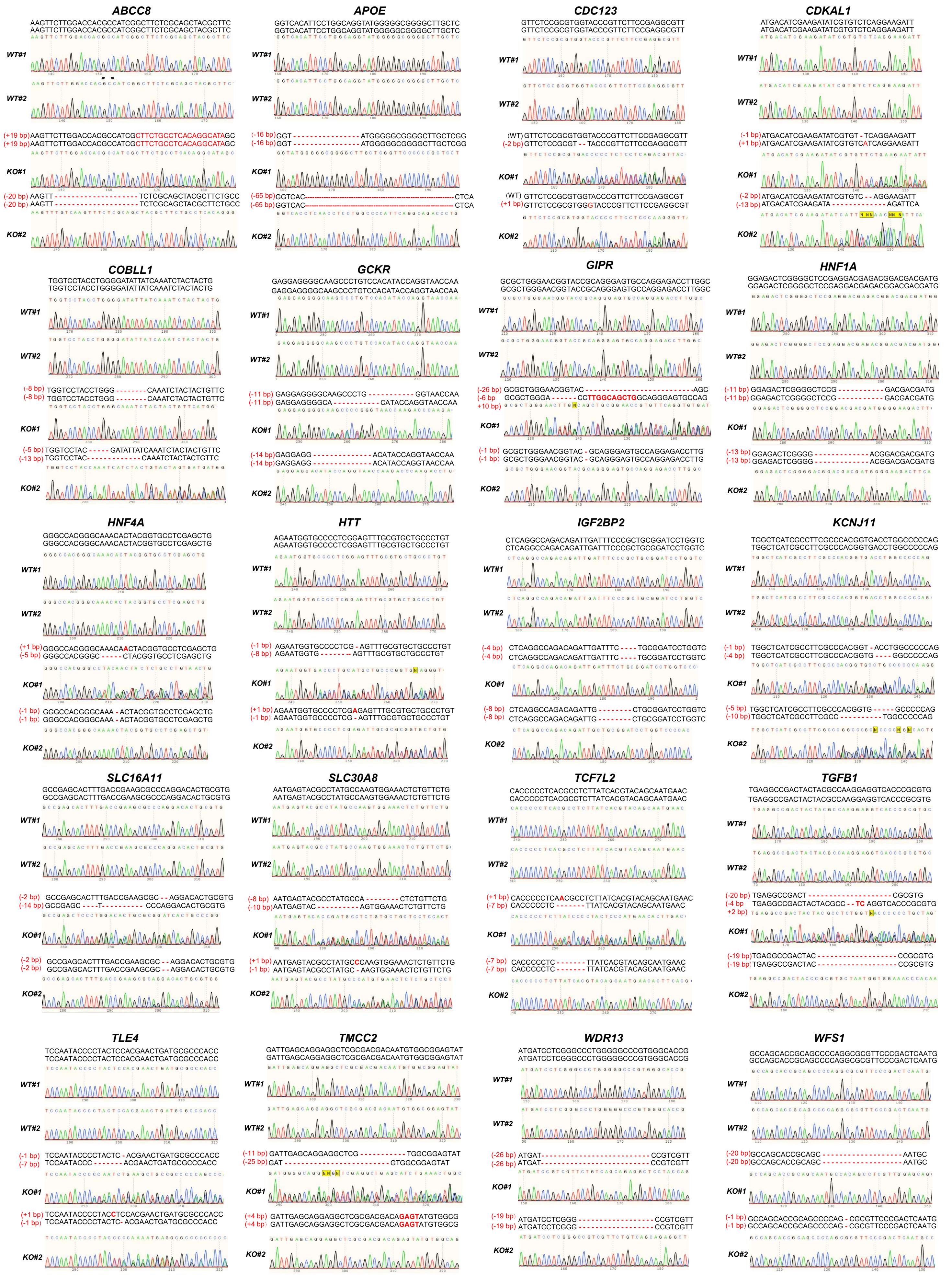
ABCC8
APOE
CDC123
CDKAL1
COBLL1
GCKR
GIPR
HNF1A
HNF4A
HTT
IGF2BP2
KCNJ11
SLC16A11
SLC30A8
TCF7L2
TGFB1
TLE4
TMCC2
WDR13
WFS1

**Figure S1. DNA sequence confirmation of frameshift mutations in isogenic KO hESC clones (two clones for each KO line) compared to two clones of *WT* hESCs.**

#### Figure S2

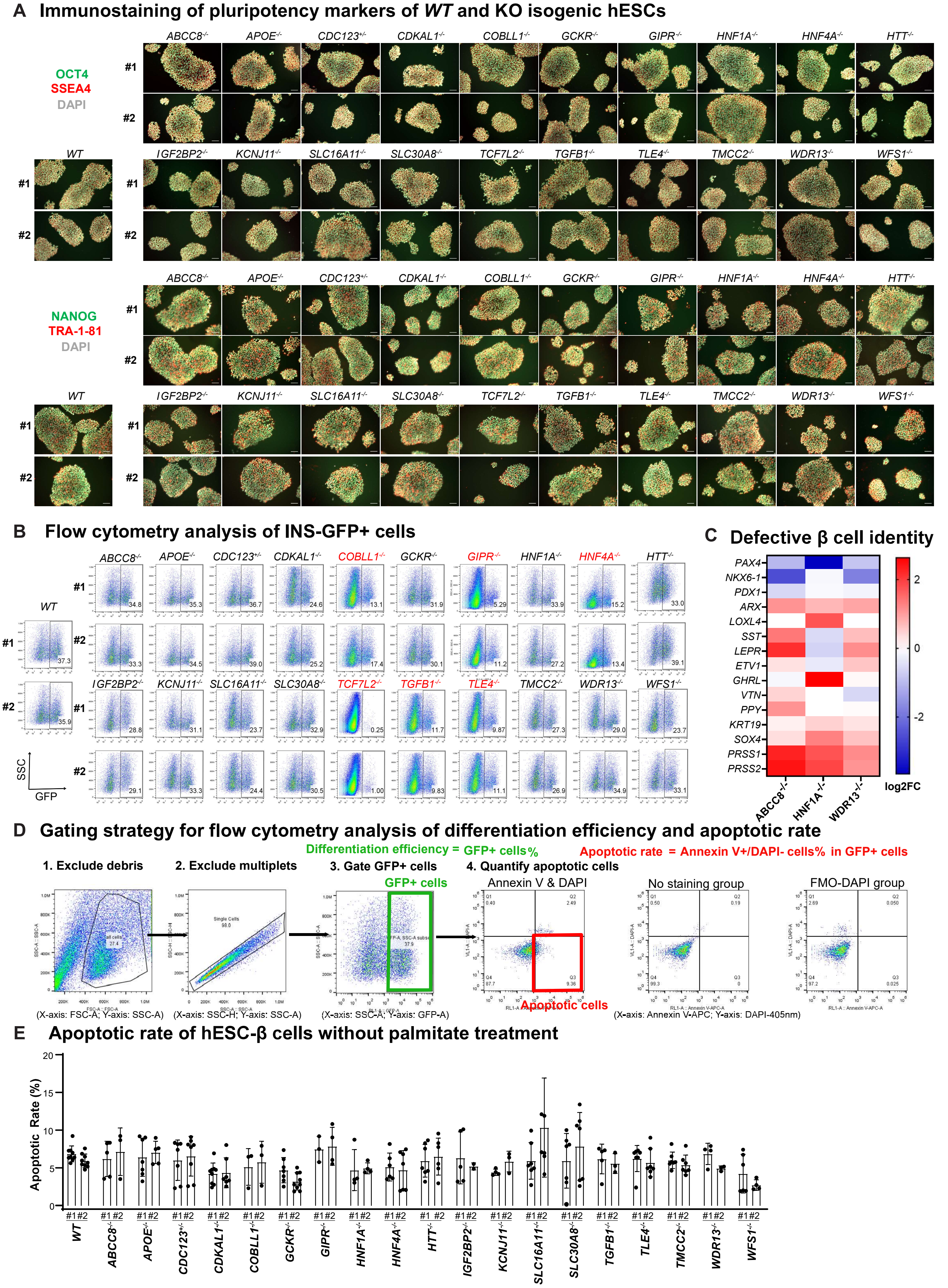
A Immunostaining of pluripotency markers of WT and KO isogenic hESCs
OCT4
SSEA4
DAPI
NANOG
TRA-1-81
DAPI
WT
#1
#2
ABCC8-/- APOE-/- CDC123+/- CDKAL1-/- COBLL1-/- GCKR-/- GIPR-/- HNF1A-/- HNF4A-/- HTT-/-
IGF2BP2-/- KCNJ11-/- SLC16A11-/- SLC30A8-/- TCF7L2-/- TGFB1-/- TLE4-/- TMCC2-/- WDR13-/- WFS1-/-
B Flow cytometry analysis of INS-GFP+ cells
SSC
GFP
WT #1 37.3
WT #2 35.9
#1: ABCC8-/- 34.8, APOE-/- 35.3, CDC123+/- 36.7, CDKAL1-/- 24.6, COBLL1-/- 13.1, GCKR-/- 31.9, GIPR-/- 5.29, HNF1A-/- 33.9, HNF4A-/- 15.2, HTT-/- 33.0
#2: ABCC8-/- 33.3, APOE-/- 34.5, CDC123+/- 39.0, CDKAL1-/- 25.2, COBLL1-/- 17.4, GCKR-/- 30.1, GIPR-/- 11.2, HNF1A-/- 27.2, HNF4A-/- 13.4, HTT-/- 39.1
#1: IGF2BP2-/- 28.8, KCNJ11-/- 31.1, SLC16A11-/- 23.7, SLC30A8-/- 32.9, TCF7L2-/- 0.25, TGFB1-/- 11.7, TLE4-/- 9.87, TMCC2-/- 27.3, WDR13-/- 29.0, WFS1-/- 23.7
#2: IGF2BP2-/- 29.1, KCNJ11-/- 33.3, SLC16A11-/- 24.4, SLC30A8-/- 30.5, TCF7L2-/- 1.00, TGFB1-/- 9.83, TLE4-/- 11.1, TMCC2-/- 26.9, WDR13-/- 34.9, WFS1-/- 33.1
C Defective β cell identity
PAX4
NKX6-1
PDX1
ARX
LOXL4
SST
LEPR
ETV1
GHRL
VTN
PPY
KRT19
SOX4
PRSS1
PRSS2
ABCC8-/-
HNF1A-/-
WDR13-/-
2
0
-2
log2FC
D Gating strategy for flow cytometry analysis of differentiation efficiency and apoptotic rate
Differentiation efficiency = GFP+ cells%
Apoptotic rate = Annexin V+/DAPI- cells% in GFP+ cells
1. Exclude debris
2. Exclude multiplets
3. Gate GFP+ cells
4. Quantify apoptotic cells
GFP+ cells
Annexin V & DAPI
No staining group
FMO-DAPI group
Single Cells
98.0
Apoptotic cells
(X-axis: FSC-A; Y-axis: SSC-A)
(X-axis: SSC-H; Y-axis: SSC-A)
(X-axis: SSC-A; Y-axis: GFP-A)
(X-axis: Annexin V-APC; Y-axis: DAPI-405nm)
E Apoptotic rate of hESC-β cells without palmitate treatment
Apoptotic Rate (%)
20
15
10
5
0
#1 #2
WT ABCC8-/- APOE-/- CDC123+/- CDKAL1-/- COBLL1-/- GCKR-/- GIPR-/- HNF1A-/- HNF4A-/- HTT-/- IGF2BP2-/- KCNJ11-/- SLC16A11-/- SLC30A8-/- TGFB1-/- TLE4-/- TMCC2-/- WDR13-/- WFS1-/-

**Figure S2. Functional characterization of isogenic WT and KO hESCs and hESC-β cells.**

**(A)** Immunostaining of pluripotency markers OCT4 (green, top panel), SSEA4 (red, top panel), NANOG (green, bottom panel), and TRA-1-81 (red, bottom panel) for *WT* and KO isogenic hESC clones (two clones for each line). Nucleuses were stained with DAPI (grey). Scale bar = 100 μm. **(B)** Flow cytometry analysis of INS-GFP+ day 24 cells derived from isogenic *WT* and KO hESCs. **(C)** INS-producing cells derived from *ABCC8*$^{-/-}$, *HNF1A*$^{-/-}$ and *WDR13*$^{-/-}$ hESCs showed defective β cell identity by down-regulated expression of β cell marker genes and upregulated of non-β cell genes. color indicates the $\log_2$(fold change) of specific gene expression in KO hESC-β lines compared with that in *WT* hESC-β cells. **(D)** Gating strategy for flow cytometry analysis of differentiation efficiency by quantifying the GFP+ cells and apoptotic rate by quantifying the percent of apoptotic cells in GFP+ cells. INS-GFP+ cells were gated based on Insulin-GFP expression. Apoptotic cells (Annexin V+DAPI- cells) were gated based on the cell population distribution in the negative control (no staining group) and fluorescence minus one (FMO) control (only DAPI staining, no Annexin V staining). **(E)** Quantification of the percentage of apoptotic (Annexin V+DAPI-) cells in isogenic hESC-derived INS-GFP+ cells in control culture condition. *P*-values calculated by one-way ANOVA followed by Dunnet's test. The gating strategy for analysis of apoptotic rate in hESC-derived β cells is shown in **Figure S2D**. Number of biological replicates was listed in **Table S2**.

Figure S3

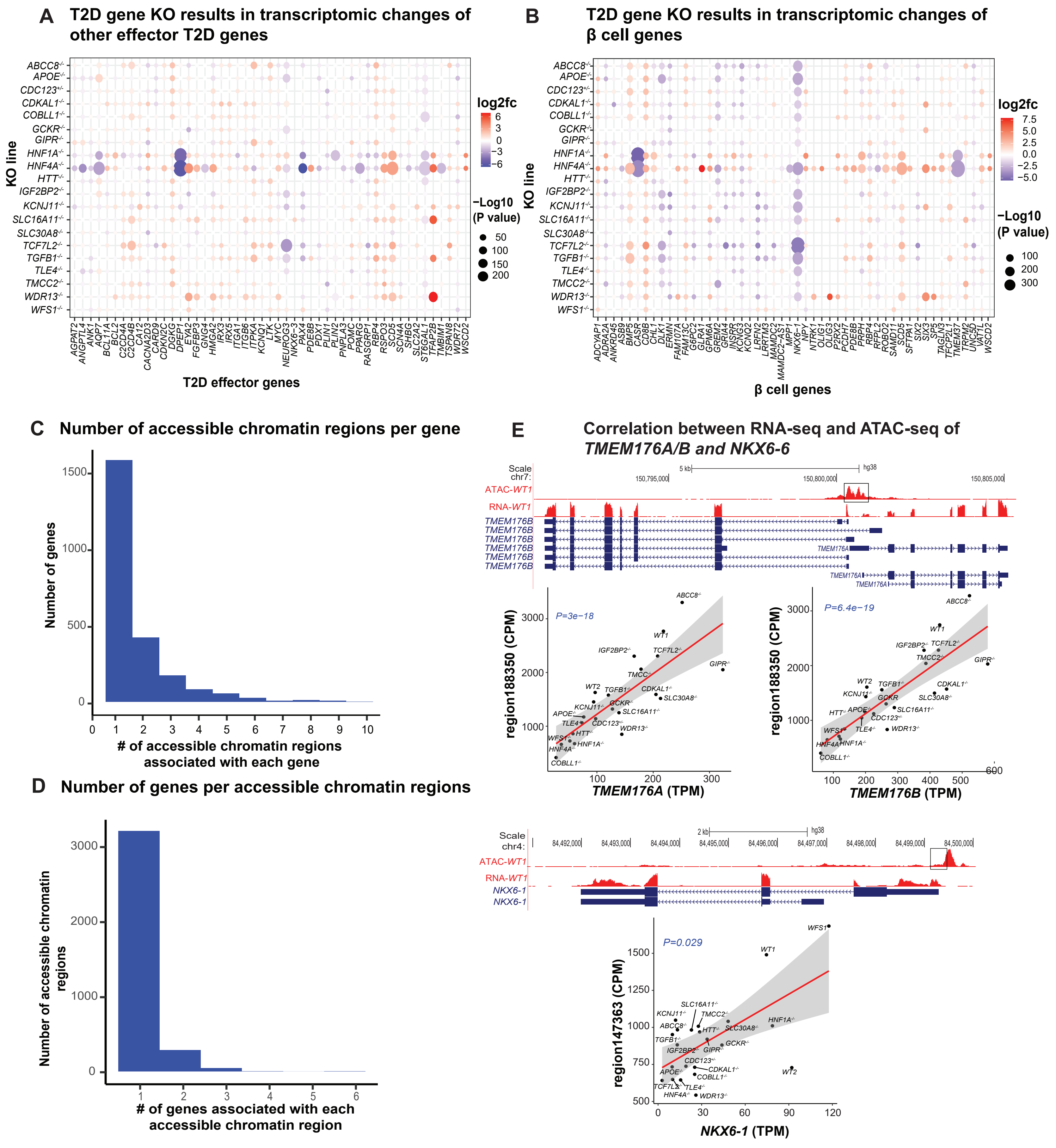
A T2D gene KO results in transcriptomic changes of other effector T2D genes
KO line
T2D effector genes
log2fc
−Log10 (P value)
B T2D gene KO results in transcriptomic changes of β cell genes
KO line
β cell genes
log2fc
−Log10 (P value)
C Number of accessible chromatin regions per gene
Number of genes
### of accessible chromatin regions associated with each gene
D Number of genes per accessible chromatin regions
Number of accessible chromatin regions
### of genes associated with each accessible chromatin region
E Correlation between RNA-seq and ATAC-seq of TMEM176A/B and NKX6-6
ATAC-WT1
RNA-WT1
TMEM176B
TMEM176A
P=3e-18
region188350 (CPM)
TMEM176A (TPM)
P=6.4e-19
TMEM176B (TPM)
NKX6-1
P=0.029
region147363 (CPM)
NKX6-1 (TPM)

**Figure S3. ATAC-seq and RNA-seq of isogenic WT and KO hESC-β cells.**

**(A)** T2D effector genes (x-axis) differentially expressed in >=1 KO line (y-axis). Top 50 genes selected according to the maximum $|\log_2(\text{fold change})|$ across all lines. **(B)** Most specific (specificity bin 1) genes for β cell expression (x-axis) differentially expressed in >=1 KO line. Top 50 genes selected according to the maximum $|\log_2(\text{fold change})|$ across all lines. **(C)** Number of accessible chromatin regions associated with expression of a gene (Methods). x-axis=# of accessible chromatin regions associated with expression of a single gene, y-axis=number of genes. Expression of a gene is associated with as many as 10 different chromatin accessible regions. **(D)** Number of genes associated with each accessible chromatin region (Methods). x-axis=# of genes associated with a chromatin accessible region, y-axis=number of accessible chromatin regions. A single accessible chromatin region can be associated with expression of as many as 6 genes. **(E)** Examples of likely novel regulatory elements that modulate expression of known T2D effector genes *TMEM176A/B* and β cell specific gene *NKX6-1*. Regulatory elements are defined as ATAC-seq peaks associated with expression of nearby genes (see Methods). Boxed region on the UCSC ATAC-seq track (ATAC-*WT1*) in the top panel indicates the associated regulatory region in each plot.

### Figure S4

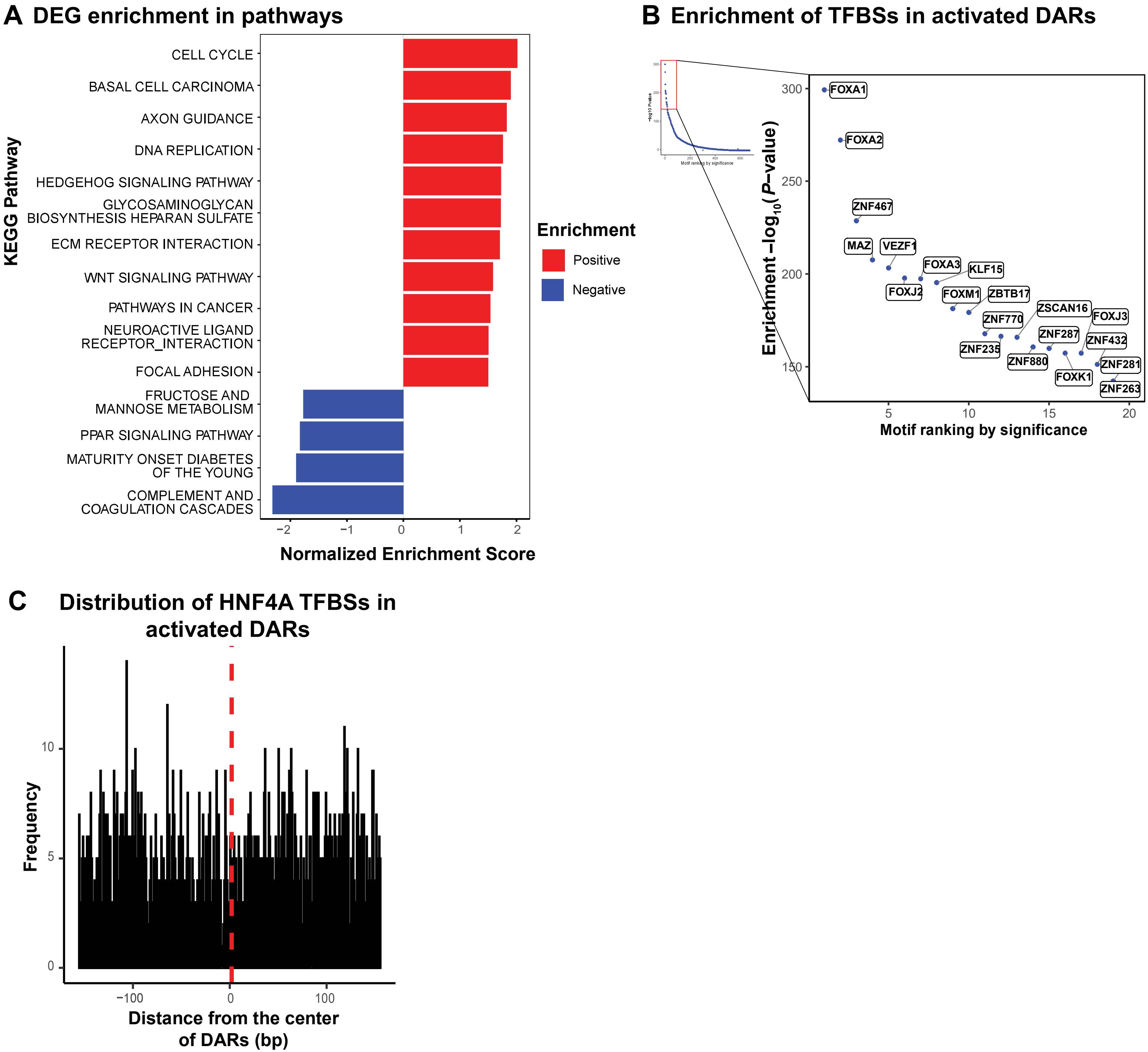
A DEG enrichment in pathways
KEGG Pathway
CELL CYCLE
BASAL CELL CARCINOMA
AXON GUIDANCE
DNA REPLICATION
HEDGEHOG SIGNALING PATHWAY
GLYCOSAMINOGLYCAN BIOSYNTHESIS HEPARAN SULFATE
ECM RECEPTOR INTERACTION
WNT SIGNALING PATHWAY
PATHWAYS IN CANCER
NEUROACTIVE LIGAND RECEPTOR_INTERACTION
FOCAL ADHESION
FRUCTOSE AND MANNOSE METABOLISM
PPAR SIGNALING PATHWAY
MATURITY ONSET DIABETES OF THE YOUNG
COMPLEMENT AND COAGULATION CASCADES
−2
−1
0
1
2
Normalized Enrichment Score
Enrichment
Positive
Negative
B Enrichment of TFBSs in activated DARs
Enrichment −log10(P−value)
300
250
200
150
FOXA1
FOXA2
ZNF467
MAZ
VEZF1
FOXA3
KLF15
FOXJ2
FOXM1
ZBTB17
ZSCAN16
FOXJ3
ZNF770
ZNF287
ZNF432
ZNF235
ZNF880
ZNF281
FOXK1
ZNF263
5
10
15
20
Motif ranking by significance
C Distribution of HNF4A TFBSs in activated DARs
Frequency
10
5
0
−100
0
100
Distance from the center of DARs (bp)

**Figure S4. Characterization of ATAC-seq and RNA-seq of HNF4A-/- hESC-β cells.**

**(A)** KEGG pathways enriched with down-regulated genes in the *HNF4A*$^{-/-}$ versus *WT* hESC-derived INS-GFP+ cells (FDR < 0.05). x-axis = normalized enrichment scores per GSEA (Methods), y-axis = KEGG pathways. Up-regulated genes were associated with enriched processes not clearly relevant to diabetes (e.g., "cell cycle", "DNA replication", "basal cell carcinoma"). **(B)** Enrichment of TFBS motifs in activated ATAC-seq peaks in *HNF4A*$^{-/-}$ hESC-β cells. Right panel shows the top 20 most enriched TFBSs. x-axis = ranking of enriched TFBSs (enrichment in decreasing order from left to right), y-axis = -log10(*P*-value) by MEME simple enrichment analysis (SEA) (Methods). **(C)** Relative distance of HNF4A TFBSs from the center of activated DARs in the purified *HNF4A*$^{-/-}$ hESC-β cells in comparison to *WT* hESC-β cells. TFBS motif abundance was generated by scanning 150bp flanking regions around centers of all activated DARs using MEME fimo (Methods). x-axis=distance of the TFBS motif from the center of DARs, y-axis= frequency of the motif occurrences.

#### Figure S5

**A** qRT-PCR analysis of expression change of insulin content associated genes in EndoC-βH1 cells

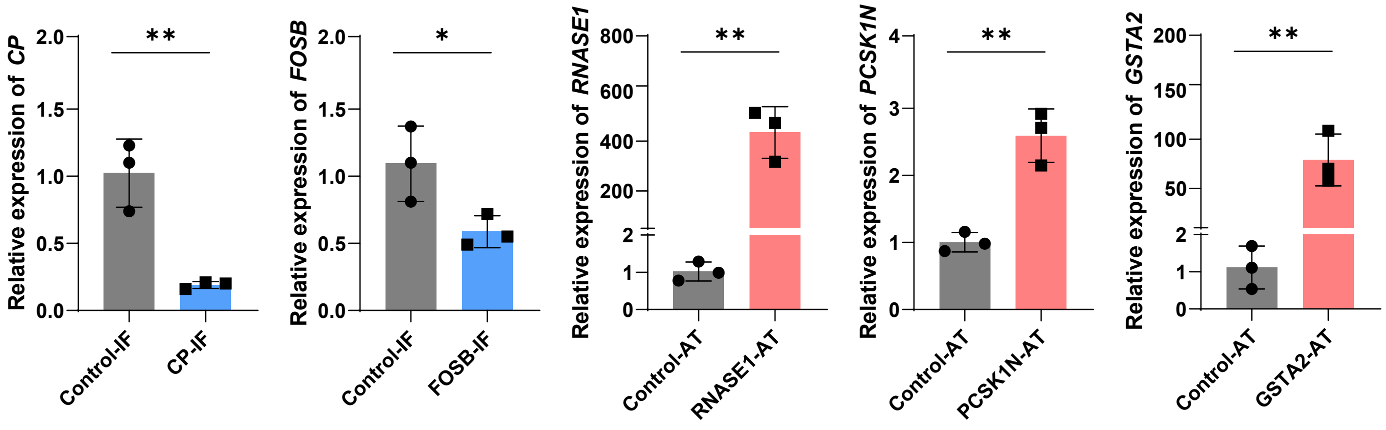


**B** qRT-PCR analysis of expression change of insulin content associated genes in EndoC-βH1-luc cells

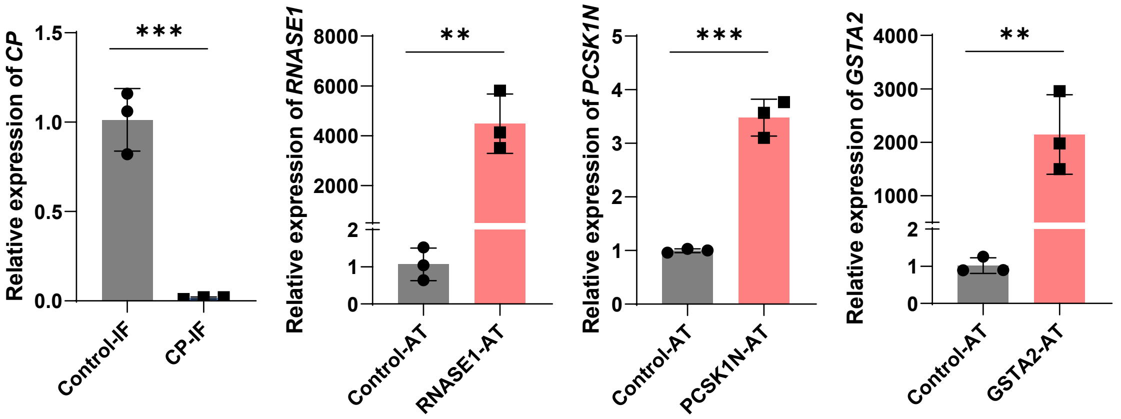


**C** qRT-PCR analysis of expression change of apoptosis associated genes in EndoC-βH1 cells

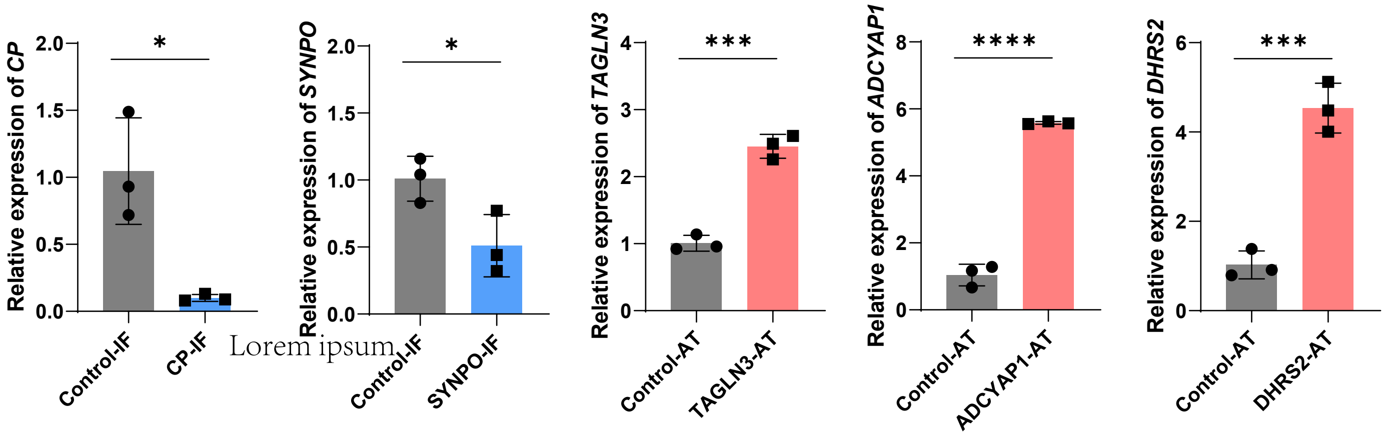


**D** Gating strategy for flow cytometry analysis of apoptotic rate in EndoC-βH1 cells

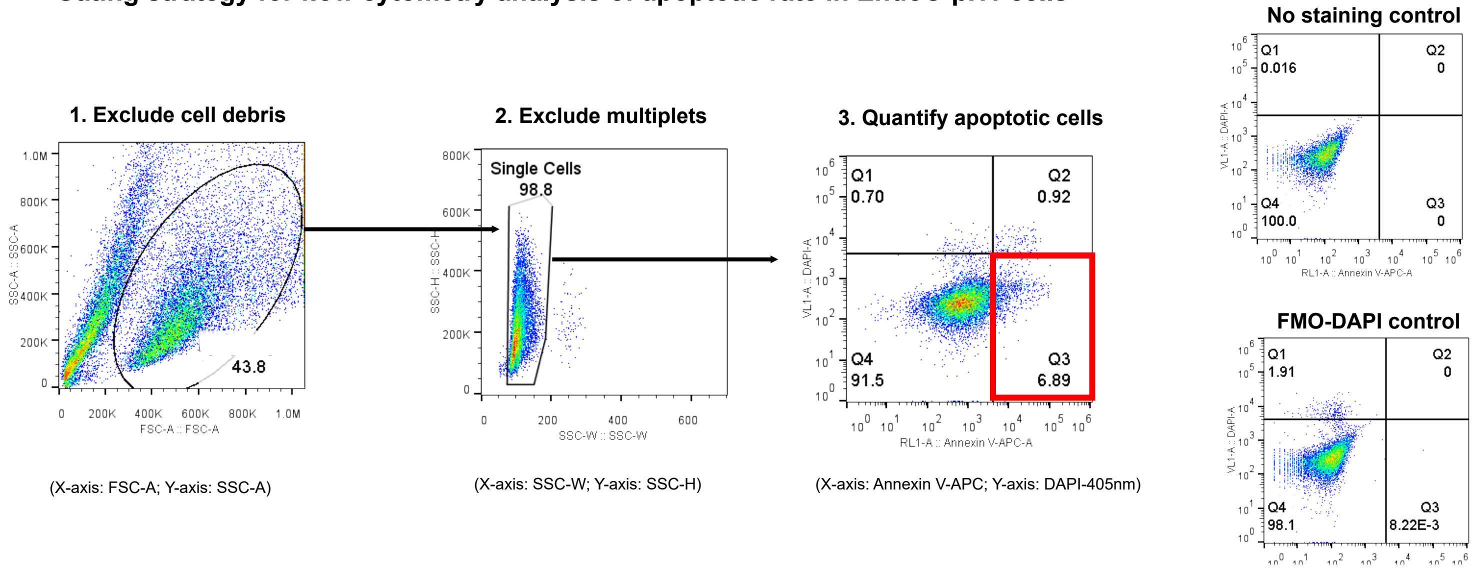


(X-axis: Annexin V-APC; Y-axis: DAPI-405nm)

**Figure S5. Functional evaluation of candidate genes that are potentially associated with insulin content or β-cell survival.**

**(A)** qRT-PCR analysis of EndoC-βH1 cells transfected with lentivirus carrying CRISPRi or CRISPRa to perturb expression of candidate genes that are potentially associated with insulin content. *CP* and *FOSB* were transcriptionally suppressed by CRISPR interference (IF) while *RNASE1*, *PCSK1N*, and *GSTA2* were transcriptionally activated by CRISPR activation (AT). N=3 biological replicates. **(B)** qRT-PCR analysis of EndoC-βH1-luc cells transfected with lentivirus carrying sgRNAs to transcriptionally inhibit expression of *CP* by CRISPRi, or sgRNAs to activate expression of *RNASE1, PCSK1N*, and *GSTA2* by CRISPRa. N=3 biological replicates. **(C)** qRT-PCR analysis of EndoC-βH1 cells transfected with lentivirus carrying CRISPRi or CRISPRa to perturb expression of candidate genes that are potentially associated with palmitate-induced apoptotic rate. *CP* and *SYNPO* were transcriptionally suppressed by CRISPR interference (IF) while *TAGLN3*, *ADCYAP1* and *DHRS2* were transcriptionally activated by CRISPR activation (AT). N=3 biological replicates. **(D)** Gating strategy for flow cytometry analysis of apoptotic rate in EndoC-βH1 cells by quantifying the percent of apoptotic (Annexin V+DAPI-) cells in all cells. Apoptotic cells (Annexin V+DAPI- cells) were gated based on the cell population distribution in the negative control (no staining group) and fluorescence minus one (FMO) control (only DAPI staining, no Annexin V staining). For panels **A-C**, data are shown as mean ± SD. *P*-values were calculated by unpaired Student's t-test. The * symbol illustrates the difference of each genetic perturbation line compared to the control line. * $P < 0.05$, ** $P < 0.01$, ***$P < 0.001$, ****$P < 0.0001$.

Figure S6

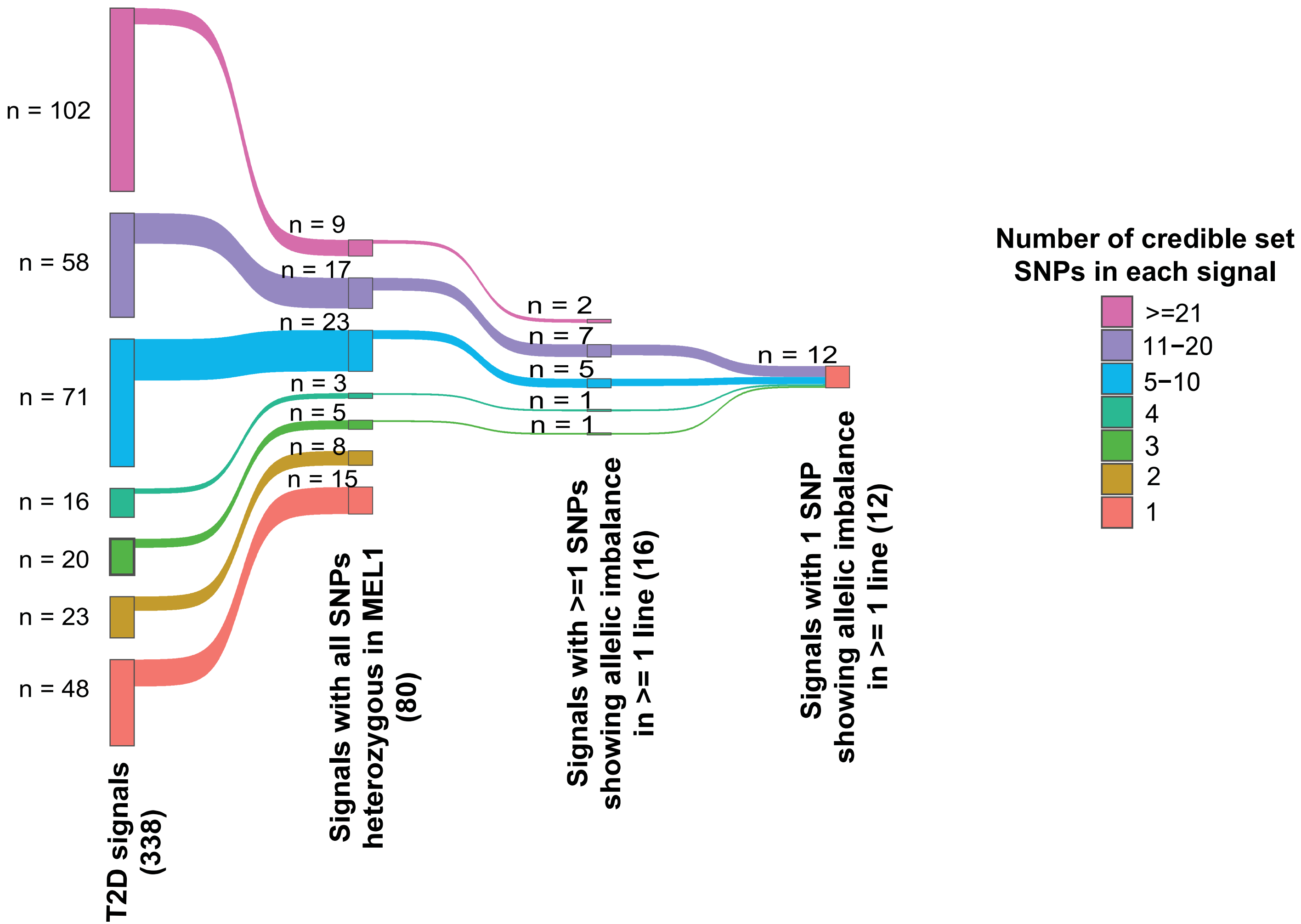
n = 102
n = 58
n = 71
n = 16
n = 20
n = 23
n = 48
n = 9
n = 17
n = 23
n = 3
n = 5
n = 8
n = 15
n = 2
n = 7
n = 5
n = 1
n = 1
n = 12
T2D signals (338)
Signals with all SNPs heterozygous in MEL1 (80)
Signals with >=1 SNPs showing allelic imbalance in >= 1 line (16)
Signals with 1 SNP showing allelic imbalance in >= 1 line (12)
Number of credible set SNPs in each signal
>=21
11−20
5−10
4
3
2
1

**Figure S6. Refinement of T2D GWAS signals using allelic imbalance analysis (binomial test from the line-specific analysis).**

Similar to **Figure 6A**, the $INS^{GFP/w}$ MEL1 hESC line is heterozygous at 80 credible set SNPs for 338 published T2D association signals[5]. Within this group of 80 SNPs, we identified at least one SNP with allelic imbalance for 16 signals using a line-specific effect analysis approach. At 12/16 signals, we identified a single SNP with allelic imbalance, and thus likely to be causal SNPS.

**Table S1**: Sequences of sgRNA targeting regions and primers used in gene KO and perturbation.

| Gene | CRSPR/Cas9 type | sgRNA target region (Recognition sequence + PAM) | Genetic consequence | Validation | | |
|---|---|---|---|---|---|---|
| | | | | Method | Primer | |
| *ABCC8* | *CRISPR/Cas9* | GCGTAGCTGCGAGAAGCCG ATGG | Frame-shifting knockout | DNA sequencing | Forward | TACACACCCAGGCACGCTTATT |
| | | | | | Reverse | TACACGGGCGGGTAAAACAAGC |
| *APOE* | *CRISPR/Cas9* | TGCGTTGCTGGTCACATTCC TGG | Frame-shifting knockout | DNA sequencing | Forward | TGGTAAATGTGCTGGGATTAGGC |
| | | | | | Reverse | GTCAAATCGCTGTTCAGAGCCAG |
| *CDC123* | *CRISPR/Cas9* | TCGGAAGAACGGGTACCACG CGG | Frame-shifting knockout | DNA sequencing | Forward | TCCTCCGCTTCCTTTTCTCG |
| | | | | | Reverse | AAACCGTGGTACCCGTTCTTCCGAC |
| *CDKAL1* | *CRISPR/Cas9* | CATCGAAGATATCGTGTCTC AGG | Frame-shifting knockout | DNA sequencing | Forward | TTGTTGAAGAAATAGTTCCTGTTGA |
| | | | | | Reverse | AAAGCTTACCTGTCACTTGGTG |
| *COBLL1* | *CRISPR/Cas9* | GTAGATTTGATAATATCCCCA GG | Frame-shifting knockout | DNA sequencing | Forward | ATCCTAATTCGACCATCCAACAC |
| | | | | | Reverse | ACGAGATATGAAAGGGTTCCAAA |
| *GCKR* | *CRISPR/Cas9* | GGTTACCTGGTATGTGGACA GGG | Frame-shifting knockout | DNA sequencing | Forward | GTGCAGGCTGAGTTTCTGGTGTG |
| | | | | | Reverse | AAGGAGGACGATGAATGGGGGAG |
| *GIPR* | *CRISPR/Cas9* | GCGCTGGGAACGGTACCGC AGGG | Frame-shifting knockout | DNA sequencing | Forward | AACAACACGCAACAGACCCTCAA |
| | | | | | Reverse | TCAAGGTTTATTTCTGTGCAGGC |
| *HNF1A* | *CRISPR/Cas9* | TCGGGGCTCCGAGGACGAG ACGG | Frame-shifting knockout | DNA sequencing | Forward | AAGGAGTTTGGTTTGTGTCTGCC |
| | | | | | Reverse | ACTAGGCTAAAGGCTGAAGGTCA |
| *HNF4A* | *CRISPR/Cas9* | CGGGCCACGGGCAAACACTA CGG | Frame-shifting knockout | DNA sequencing | Forward | CACTGAGTGGGGAGGTGATG |
| | | | | | Reverse | GCTCAATAAACGGCAGCCAG |
| *HTT* | *CRISPR/Cas9* | GGCAGCACGCAAACTCCGAG GGG | Frame-shifting knockout | DNA sequencing | Forward | GGAAATGCTGTTTGGTAGACGAT |
| | | | | | Reverse | ACATGGATTAGTGCTATGAGGGA |
| *IGF2BP2* | *CRISPR/Cas9* | GACAGATTGATTTCCCGCTG CGG | Frame-shifting knockout | DNA sequencing | Forward | TTTCTGACGAAGCTCCCCTG |
| | | | | | Reverse | TCCTGTTGAAAGAAGCCACTG |
| *KCNJ11* | *CRISPR/Cas9* | CGCCTTCGCCCACGGTGACC TGG | Frame-shifting knockout | DNA sequencing | Forward | ACAAGAACATCCGGGAGCAG |
| | | | | | Reverse | GTCACCCACACGTAGCATGA |
| *SLC16A11* | *CRISPR/Cas9* | GCACTTTGACCGAAGCGCCC AGG | Frame-shifting knockout | DNA sequencing | Forward | CCGGGAGTTCCAGCCCTA |
| | | | | | Reverse | TGCCCTTTCTCAGGTTGTCG |
| *SLC30A8* | *CRISPR/Cas9* | ATGAGTACGCCTATGCCAAG TGG | Frame-shifting knockout | DNA sequencing | Forward | TAGCGGTTCCAAGTATGGGC |
| | | | | | Reverse | TGCATTGTACAAAACACTCCCT |
| *TCF7L2* | *CRISPR/Cas9* | TACGTGATAAGAGGCGTGAG GGG | Frame-shifting knockout | DNA sequencing | Forward | GCCAGGCTTAGTGGGGAAAT |
| | | | | | Reverse | AACCACCCCAAAGAGCTTCC |
| *TGFB1* | *CRISPR/Cas9* | GGCCGACTACTACGCCAAGG AGG | Frame-shifting knockout | DNA sequencing | Forward | GACTATCGACATGGAGCTGGTGA |
| | | | | | Reverse | CTCCTCTCCAAGACCAGACACCT |

| | | | | | | | |
|---|---|---|---|---|---|---|---|
| *TLE4* | *CRISPR/Cas9* | GCGCATCAGTTCGTGGAGTA GGG | | Frame-shifting knockout | DNA sequencing | Forward | TCTGTTTCTCTTGGGGCAGG |
| | | | | | | Reverse | AAATAGGCCCCTCTGCCTCT |
| *TMCC2* | *CRISPR/Cas9* | GGAGGCTCGCGACGACAAT GTGG | | Frame-shifting knockout | DNA sequencing | Forward | GGCTGTCAGGGATACTGCTCTTG |
| | | | | | | Reverse | GGTAGTGCTCCAGCTTCTTGTGC |
| *WDR13* | *CRISPR/Cas9* | TGACAGAACGACGGTGCCCA CGG | | Frame-shifting knockout | DNA sequencing | Forward | CAGAACTCGTGAGGAAGCCACTG |
| | | | | | | Reverse | AAACATGATACATGCCCGCAAAG |
| *WFS1* | *CRISPR/Cas9* | ATTGAGTCGGGAACGCGCCT GGG | | Frame-shifting knockout | DNA sequencing | Forward | TGCCTCCCTCTGCTTTTCTG |
| | | | | | | Reverse | GGAGCTGCACAATGCTGAAC |
| *RNASE1* | *CRISPR/dCas9 -VPR* | *sgRNA1* | TCTGTATAAGGTCC ACACCCCGG | Transcriptional activation | qRT-PCR | Forward | CCTGGTAGATGTCCAGAATGTC |
| | | *sgRNA2* | ATAGGAAGAAAGAC TAACGTAGG | | | Reverse | TGCTGGAGTTGCTCTTGTAG |
| *PCSK1N* | *CRISPR/dCas9 -VPR* | *sgRNA1* | TGCGCCTGCGTCG GTCGGGTGGG | Transcriptional activation | qRT-PCR | Forward | AGGCCGAACGTCAGGAG |
| | | *sgRNA2* | GTCGCCATGGAGA TGCCCAAAGG | | | Reverse | CCAGAGCCGGATCAGAGTT |
| *GSTA2* | *CRISPR/dCas9 -VPR* | *sgRNA1* | TAACTTGACCCTTC TTTCAGTGG | Transcriptional activation | qRT-PCR | Forward | ACGGACAAGACTACCTTGTTG |
| | | *sgRNA2* | CTCATGTCTTAGAA TCCAGTAGG | | | Reverse | GGAAGCTGGAAATAAGGCTAGAG |
| *TAGLN3* | *CRISPR/dCas9 -VPR* | *sgRNA1* | ATTCCCGACAGATG GACAGTTGG | Transcriptional activation | qRT-PCR | Forward | CATACCCAAGATCTCAGAGTCAAA |
| | | *sgRNA2* | TGAGGTAACAACTG TCTGGCAGG | | | Reverse | CTGAAAGATGTCGGTGGTTCT |
| *ADCYAP1* | *CRISPR/dCas9 -VPR* | *sgRNA1* | GTGGCCGTAAGTC CACCCGGAGG | Transcriptional activation | qRT-PCR | Forward | CGCCCACGGGATCCTTAAC |
| | | *sgRNA2* | GTTTGTAGACGCGC AGAACCAGG | | | Reverse | AGCGACTGCAGGTGCTTC |
| *DHRS2* | *CRISPR/dCas9 -VPR* | *sgRNA1* | CAACTTGGGAGGA AACCTAGGGG | Transcriptional activation | qRT-PCR | Forward | CGTCGACTTCCTGGTGTG |
| | | *sgRNA2* | TCTCCAAGTCCCCA CCAGATTGG | | | Reverse | CCAGATCTGCTCACTGGTC |
| *CP* | *CRISPR/dCas9 -KRAB* | *sgRNA1* | TTGGAATTATTGAA ACGACTTGG | Transcriptional supression | qRT-PCR | Forward | AGGCCCTTTATCTTCAGTACAC |
| | | *sgRNA2* | TTACGAGAACTTAA CTGAATTGG | | | Reverse | CTTTATCTCCAGTTTCAGCTTTGAT |
| *FOSB* | *CRISPR/dCas9 -KRAB* | *sgRNA1* | AGTGCGGGACTCG ATTTGGCGGG | Transcriptional supression | qRT-PCR | Forward | GGAACCAGCTACTCCACAC |
| | | *sgRNA2* | GGACACGCGGAAC CAAGACTTGG | | | Reverse | CAGGCCCACTGGTAGTTC |
| *SYNPO* | *CRISPR/dCas9 -KRAB* | *sgRNA1* | AGGGTGTGGTGTC CGGCTACAGG | Transcriptional supression | qRT-PCR | Forward | AAACCCAACCAGAACCTCTC |
| | | *sgRNA2* | AAGAAACACTGTGC AGTCCGGGG | | | Reverse | GGCTTGAAGACTCGATGACATA |

**Table S2:** Number of biological replicates used for molecular sequencing and cellular trait functional assays.

| KO/WT | Clone | Genetic sequencing assay | | Functional assay | | | | | |
|---|---|---|---|---|---|---|---|---|---|
| | | RNA-seq | ATAC-seq | β cell differentiation (Figure 1C) | Total intracellular insulin content (Figure 1D) | Glucose stimulated insulin secretion (Figure 1E) | KCl stimulated insulin secretion (Figure 1F) | Apoptotic rate: With Palmitate (Figure 1H) | Apoptotic rate: No Palmitate (Figure S2E) |
| *ABCC8-/-* | clone #1 | 3 | 3 | 3 | 3 | 7 | 6 | 3 | 5 |
| | clone #2 | N.A. | N.A. | 3 | 3 | 7 | 7 | 3 | 3 |
| *APOE-/-* | clone #1 | 3 | 3 | 3 | 3 | 8 | 8 | 3 | 7 |
| | clone #2 | N.A. | N.A. | 3 | 3 | 8 | 8 | 3 | 5 |
| *CDC123+/-* | clone #1 | 3 | 3 | 3 | 3 | 7 | 8 | 3 | 7 |
| | clone #2 | N.A. | N.A. | 3 | 3 | 6 | 8 | 3 | 8 |
| *CDKAL1-/-* | clone #1 | 3 | 3 | 3 | 3 | 8 | 8 | 3 | 9 |
| | clone #2 | N.A. | N.A. | 3 | 3 | 7 | 8 | 3 | 7 |
| *COBLL1-/-* | clone #1 | 3 | 3 | 3 | 3 | 8 | 7 | 3 | 3 |
| | clone #2 | N.A. | N.A. | 3 | 3 | 8 | 8 | 3 | 3 |
| *GCKR-/-* | clone #1 | 3 | 3 | 3 | 3 | 8 | 7 | 3 | 7 |
| | clone #2 | N.A. | N.A. | 3 | 3 | 8 | 7 | 3 | 9 |
| *GIPR-/-* | clone #1 | 3 | 3 | 3 | 3 | 8 | 8 | 3 | 3 |
| | clone #2 | N.A. | N.A. | 3 | 3 | 8 | 8 | 3 | 4 |
| *HNF1A-/-* | clone #1 | 3 | 3 | 3 | 3 | 7 | 8 | 3 | 4 |
| | clone #2 | N.A. | N.A. | 3 | 3 | 8 | 6 | 3 | 4 |
| *HNF4A-/-* | clone #1 | 3 | 3 | 3 | 3 | 8 | 8 | 3 | 6 |
| | clone #2 | N.A. | N.A. | 3 | 3 | 8 | 8 | 3 | 7 |
| *HTT-/-* | clone #1 | 3 | 3 | 3 | 3 | 8 | 8 | 3 | 7 |
| | clone #2 | N.A. | N.A. | 3 | 3 | 8 | 8 | 3 | 6 |
| *IGF2BP2-/-* | clone #1 | 3 | 3 | 3 | 3 | 8 | 7 | 3 | 5 |
| | clone #2 | N.A. | N.A. | 3 | 3 | 8 | 7 | 3 | 2 |

| | | | | | | | | | |
|---|---|---|---|---|---|---|---|---|---|
| *KCNJ11-/-* | clone #1 | 3 | 3 | 3 | 3 | 7 | 8 | 3 | 4 |
| | clone #2 | N.A. | N.A. | 3 | 3 | 5 | 6 | 3 | 3 |
| *SLC16A11-/-* | clone #1 | 3 | 3 | 3 | 3 | 6 | 7 | 3 | 7 |
| | clone #2 | N.A. | N.A. | 3 | 3 | 8 | 8 | 3 | 7 |
| *SLC30A8-/-* | clone #1 | 3 | 3 | 3 | 3 | 7 | 8 | 3 | 7 |
| | clone #2 | N.A. | N.A. | 3 | 3 | 6 | 7 | 3 | 7 |
| *TCF7L2-/-* | clone #1 | 3 | 3 | 3 | 3 | N.A. | N.A. | N.A. | N.A. |
| | clone #2 | N.A. | N.A. | 3 | 3 | N.A. | N.A. | N.A. | N.A. |
| *TGFB1-/-* | clone #1 | 3 | 3 | 3 | 3 | 7 | 8 | 3 | 5 |
| | clone #2 | N.A. | N.A. | 3 | 3 | 8 | 8 | 3 | 3 |
| *TLE4-/-* | clone #1 | 3 | 3 | 3 | 3 | 8 | 7 | 3 | 7 |
| | clone #2 | N.A. | N.A. | 3 | 3 | 8 | 8 | 3 | 7 |
| *TMCC2-/-* | clone #1 | 3 | 3 | 3 | 3 | 8 | 7 | 3 | 7 |
| | clone #2 | N.A. | N.A. | 3 | 3 | 7 | 7 | 3 | 7 |
| *WDR13-/-* | clone #1 | 3 | 3 | 3 | 3 | 6 | 8 | 3 | 4 |
| | clone #2 | N.A. | N.A. | 3 | 3 | 8 | 6 | 3 | 3 |
| *WFS1-/-* | clone #1 | 4 | 4 | 3 | 3 | 6 | 7 | 3 | 6 |
| | clone #2 | N.A. | N.A. | 3 | 3 | 6 | 6 | 3 | 4 |
| *WT1* | clone #1 | 3 | 3 | 6 | 6 | 8 | 7 | 6 | 9 |
| *WT2* | clone #1 | 3 | 3 | 6 | 6 | 8 | 8 | 6 | 9 |

**Supplementary Table 3:** 99% credible set of SNPs that overlap suppressed DARs in *HNF4A*$^{-/-}$ and disrupt HNF4A TFBSs.

| chrome | SNP_start | SNP_end | rsid | PPA in 99% credible set | GWAS trait and signal | # of credible set SNPs | Protein coding genes (DEGs)in 200kb region centered around the SNP |
|---|---|---|---|---|---|---|---|
| 10 | 112739855 | 112739856 | rs34033101 | 0.001141876 | T2D_fmap.TCF7L2.chr10:115247447 | 18 | *VTI1A* |
| 12 | 49869364 | 49869365 | rs7132908 | 0.916715738 | T2D_fmap.FAIM2.chr12:50263148 | 2 | *BCDIN3D,FAIM2,AQP2,AQP6* |

**Supplementary Table 4:** Genes associated with insulin content and apoptotic rate across 22 cell lines.

| Genes associated with insulin content | | | | | | |
|---|---|---|---|---|---|---|
| **EnsemblID** | **Symbol** | **Log2FC** | **StdErr** | **MeanCP10K** | **P-Value** | **FDR** |
| ENSG00000142910 | *TINAGL1* | -0.6202 | 0.1073 | 8.08E-09 | 0.0958 | 3.06E-05 |
| ENSG00000075673 | *ATP12A* | -0.8773 | 0.1815 | 4.59E-06 | 0.1539 | 0.0010 |
| ENSG00000129538 | *RNASE1* | -0.8131 | 0.1730 | 6.54E-06 | 0.0979 | 0.0012 |
| ENSG00000132329 | *RAMP1* | -0.6565 | 0.1426 | 7.55E-06 | 0.0992 | 0.0013 |
| ENSG00000140090 | *SLC24A4* | 0.6931 | 0.1638 | 8.49E-06 | 0.2301 | 0.0014 |
| ENSG00000139219 | *COL2A1* | -0.6479 | 0.1758 | 0.0003 | 4.7639 | 0.0070 |
| ENSG00000137573 | *SULF1* | -0.6855 | 0.1769 | 0.0003 | 0.1589 | 0.0076 |
| ENSG00000172116 | *CD8B* | -0.5972 | 0.1682 | 0.0005 | 0.0872 | 0.0090 |
| ENSG00000141933 | *TPGS1* | -0.6101 | 0.1603 | 0.0005 | 0.0867 | 0.0090 |
| ENSG00000102109 | *PCSK1N* | -0.7487 | 0.1977 | 0.0008 | 3.2214 | 0.0118 |
| ENSG00000243449 | *C4orf48* | -0.7914 | 0.2119 | 0.0013 | 0.0626 | 0.0153 |
| ENSG00000244067 | *GSTA2* | -0.7361 | 0.2426 | 0.0019 | 0.2241 | 0.0185 |
| ENSG00000047457 | *CP* | 0.8153 | 0.3075 | 0.0024 | 0.5669 | 0.0211 |
| ENSG00000221843 | *C2orf16* | 0.5826 | 0.2199 | 0.0031 | 0.1488 | 0.0241 |
| ENSG00000166126 | *AMN* | -0.7220 | 0.2314 | 0.0032 | 0.3672 | 0.0244 |
| ENSG00000204983 | *PRSS1* | -0.6273 | 0.2279 | 0.0037 | 0.2435 | 0.0268 |
| ENSG00000141433 | *ADCYAP1* | -0.6980 | 0.2117 | 0.0038 | 0.0814 | 0.0271 |
| ENSG00000225783 | *MIAT* | 0.5939 | 0.2165 | 0.0044 | 1.7289 | 0.0294 |
| ENSG00000197106 | *SLC6A17* | -0.9021 | 0.2636 | 0.0054 | 0.1127 | 0.0333 |
| ENSG00000129951 | *PLPPR3* | -0.7109 | 0.2169 | 0.0056 | 0.0886 | 0.0341 |
| ENSG00000125740 | *FOSB* | 0.7237 | 0.2849 | 0.0056 | 0.7248 | 0.0344 |
| **Genes associated with apoptotic rate** | | | | | | |
| **EnsemblID** | **Symbol** | **Log2FC** | **StdErr** | **MeanCP10K** | **P-Value** | **FDR** |
| ENSG00000137573 | *SULF1* | 0.8446 | 0.1541 | 2.89E-07 | 0.1589 | 0.0006 |
| ENSG00000144834 | *TAGLN3* | 0.7045 | 0.1415 | 1.00E-06 | 0.1223 | 0.0007 |
| ENSG00000141469 | *SLC14A1* | -0.8586 | 0.1906 | 4.94E-06 | 0.0928 | 0.0016 |
| ENSG00000171992 | *SYNPO* | -0.7039 | 0.1642 | 1.22E-05 | 2.0761 | 0.0023 |
| ENSG00000197106 | *SLC6A17* | 1.0690 | 0.2269 | 1.26E-05 | 0.1127 | 0.0023 |
| ENSG00000047457 | *CP* | -1.0711 | 0.2750 | 2.35E-05 | 0.5669 | 0.0028 |
| ENSG00000129951 | *PLPPR3* | 0.8242 | 0.1945 | 3.92E-05 | 0.0886 | 0.0036 |
| ENSG00000180875 | *GREM2* | -0.6154 | 0.1522 | 4.27E-05 | 0.0656 | 0.0037 |
| ENSG00000178531 | *CTXN1* | 0.7589 | 0.1770 | 4.58E-05 | 0.1389 | 0.0039 |
| ENSG00000140090 | *SLC24A4* | -0.6359 | 0.1700 | 5.99E-05 | 0.2301 | 0.0043 |
| ENSG00000141433 | *ADCYAP1* | 0.7639 | 0.1930 | 6.04E-05 | 0.0814 | 0.0043 |
| ENSG00000138829 | *FBN2* | 0.6687 | 0.1716 | 9.65E-05 | 0.0839 | 0.0055 |
| ENSG00000277586 | *NEFL* | 0.7864 | 0.1993 | 0.0001 | 0.1194 | 0.0060 |
| ENSG00000171388 | *APLN* | -0.6452 | 0.1636 | 0.0001 | 0.0710 | 0.0060 |

| | | | | | | |
|---|---|---|---|---|---|---|
| ENSG00000133083 | *DCLK1* | 1.0634 | 0.2788 | 0.0001 | 0.1081 | 0.0063 |
| ENSG00000170390 | *DCLK2* | 0.6673 | 0.1660 | 0.0001 | 0.1485 | 0.0063 |
| ENSG00000146122 | *DAAM2* | 0.5932 | 0.1484 | 0.0002 | 0.3494 | 0.0066 |
| ENSG00000184347 | *SLIT3* | 0.5905 | 0.1580 | 0.0002 | 0.1687 | 0.0070 |
| ENSG00000225783 | *MIAT* | -0.7177 | 0.2076 | 0.0003 | 1.7289 | 0.0089 |
| ENSG00000132688 | *NES* | 0.8282 | 0.2301 | 0.0003 | 0.1169 | 0.0093 |
| ENSG00000153902 | *LGI4* | -0.6072 | 0.1750 | 0.0010 | 0.1028 | 0.0147 |
| ENSG00000113657 | *DPYSL3* | 0.6117 | 0.1878 | 0.0012 | 0.4707 | 0.0167 |
| ENSG00000131747 | *TOP2A* | 0.5807 | 0.1911 | 0.0016 | 0.1309 | 0.0185 |
| ENSG00000170419 | *VSTM2A* | 0.6374 | 0.2128 | 0.0018 | 0.2943 | 0.0198 |
| ENSG00000132470 | *ITGB4* | 0.6816 | 0.2144 | 0.0021 | 0.0843 | 0.0218 |
| ENSG00000224057 | *EGFR-AS1* | -0.7951 | 0.2612 | 0.0022 | 0.2188 | 0.0222 |
| ENSG00000243449 | *C4orf48* | 0.7399 | 0.2146 | 0.0025 | 0.0626 | 0.0235 |
| ENSG00000100867 | *DHRS2* | 0.7808 | 0.2225 | 0.0030 | 0.2777 | 0.0255 |
| ENSG00000135046 | *ANXA1* | 0.5955 | 0.1980 | 0.0040 | 0.0646 | 0.0296 |
| ENSG00000101144 | *BMP7* | 0.6701 | 0.2298 | 0.0041 | 0.1307 | 0.0298 |
| ENSG00000143320 | *CRABP2* | 0.6347 | 0.2257 | 0.0042 | 0.1128 | 0.0303 |
| ENSG00000112414 | *ADGRG6* | 0.7131 | 0.2905 | 0.0043 | 0.1518 | 0.0307 |
| ENSG00000131409 | *LRRC4B* | 0.6354 | 0.2278 | 0.0046 | 0.1229 | 0.0319 |
| ENSG00000165655 | *ZNF503* | 0.6080 | 0.2255 | 0.0057 | 0.0783 | 0.0358 |
| ENSG00000137801 | *THBS1* | 0.6285 | 0.2314 | 0.0093 | 0.3226 | 0.0470 |

**Supplementary Table 5:** Summary of ATAC-seq allelic imbalance signatures at heterozygous SNPs in 99% credible sets for T2D association.

| Allelic imbalance analysis using binomial test (common effect and line-specific analyses) | | | | | | | | | |
|---|---|---|---|---|---|---|---|---|---|
| **Genetic association credible set signal** | **Variant** | **Variant location** | **Reference allele** | **Alternate allele** | **Genetic association PPA** | **Number of lines with allelic imbalance (FDR<5%)** | **Common effect analysis (/Meta-analysis) P-value** | **Common effect analysis (/Meta-analysis) FDR<5%** | **Fraction of credible set variants that are heterozygous** |
| T2D_fmap.ADCY5.chr3:123065778 | rs11708067 | chr3:123346931 | A | G | 0.95945 | 2 | 1.419E-24 | TRUE | 1.00 |
| T2D_fmap.AUTS2.chr7:69055951 | rs6974687 | chr7:70044199 | T | C | 0.00035 | 1 | 5.278E-04 | TRUE | 0.94 |
| T2D_fmap.BOP1.chr8:145972670 | rs2721138 | chr8:144524754 | T | C | 0.01151 | 1 | 3.630E-03 | FALSE | 0.96 |
| T2D_fmap.BOP1.chr8:145972670 | rs2954658 | chr8:144792397 | C | T | 0.00121 | 0 | 1.132E-11 | TRUE | 0.96 |
| T2D_fmap.BOP1.chr8:145972670 | rs2955195 | chr8:144763807 | C | T | 0.00055 | 0 | 1.384E-04 | TRUE | 0.96 |
| T2D_fmap.BOP1.chr8:145972670 | rs2958521 | chr8:144798574 | C | G | 0.00101 | 0 | 6.689E-08 | TRUE | 0.96 |
| T2D_fmap.BOP1.chr8:145972670 | rs4637890 | chr8:144786875 | G | A | 0.00099 | 4 | 1.391E-25 | TRUE | 0.96 |
| 2D_fmap.C2CD4A-C2CD4B.chr15:6239160 | rs7163757 | chr15:62099409 | C | T | 0.22696 | 1 | 2.366E-07 | TRUE | 1.00 |
| T2D_fmap.CCND2.chr12:4384696 | 12:4384696_C_T | chr12:4275530 | C | T | 1.00000 | 0 | 1.413E-08 | TRUE | 1.00 |
| 2D_fmap.CDKN2A-CDKN2B.chr9:2213398 | rs10757282 | chr9:22133985 | T | C | 0.95095 | 0 | 5.319E-07 | TRUE | 1.00 |
| 2D_fmap.CDKN2A-CDKN2B.chr9:2213409 | rs10811660 | chr9:22134069 | G | A | 0.30622 | 0 | 1.158E-07 | TRUE | 1.00 |
| 2D_fmap.CDKN2A-CDKN2B.chr9:2213409 | rs10811661 | chr9:22134095 | T | C | 0.69337 | 0 | 3.937E-11 | TRUE | 1.00 |
| T2D_fmap.CEBPB.chr20:48832020 | rs6122889 | chr20:50216686 | T | A | 0.04191 | 1 | 2.725E-03 | FALSE | 1.00 |
| T2D_fmap.CEP68.chr2:65284231 | rs2302647 | chr2:65056040 | G | A | 0.02197 | 0 | 9.703E-04 | TRUE | 1.00 |
| T2D_fmap.CEP68.chr2:65666674 | rs1807337 | chr2:65436033 | G | C | 0.01040 | 0 | 1.480E-04 | TRUE | 1.00 |
| T2D_fmap.CILP2-TM6SF2.chr19:19379549 | rs56255430 | chr19:19367068 | A | C | 0.05617 | 0 | 1.674E-10 | TRUE | 1.00 |
| T2D_fmap.CLEC14A.chr14:38803756 | rs7145965 | chr14:38335229 | G | A | 0.00272 | 0 | 2.005E-04 | TRUE | 1.00 |
| T2D_fmap.CRY2.chr11:45858584 | rs1401419 | chr11:45858188 | T | C | 0.00794 | 1 | 2.161E-01 | FALSE | 1.00 |
| T2D_fmap.CYTH1.chr17:76792179 | rs1384367 | chr17:78733675 | C | A | 0.02177 | 0 | 7.983E-05 | TRUE | 0.96 |
| T2D_fmap.ETS1.chr11:128235252 | rs1125251 | chr11:128361671 | T | C | 0.16156 | 7 | 4.717E-44 | TRUE | 1.00 |
| T2D_fmap.ETS1.chr11:128235252 | rs1125253 | chr11:128361574 | T | G | 0.11613 | 4 | 1.563E-33 | TRUE | 1.00 |

| | | | | | | | | | |
|---|---|---|---|---|---|---|---|---|---|
| T2D_fmap.EYA2.chr20:45596378 | rs6063046 | chr20:46967739 | A | G | 0.37716 | 1 | 6.185E-06 | TRUE | 0.82 |
| T2D_fmap.FARSA-ZNF799.chr19:13038415 | rs2974752 | chr19:12945743 | G | A | 0.01821 | 0 | 6.818E-10 | TRUE | 1.00 |
| T2D_fmap.FBRSL1.chr12:133069698 | 12:133066392_C_A | chr12:132489806 | C | A | 0.08668 | 0 | 3.715E-07 | TRUE | 0.86 |
| T2D_fmap.GNAS.chr20:57387352 | rs1800900 | chr20:58840055 | A | G | 0.04849 | 1 | 1.520E-08 | TRUE | 0.75 |
| T2D_fmap.GPSM1.chr9:139247229 | rs74604683 | chr9:136352777 | C | T | 0.74102 | 0 | 2.773E-24 | TRUE | 1.00 |
| T2D_fmap.GRB10.chr7:50809085 | rs56037894 | chr7:50718556 | G | A | 0.05195 | 0 | 4.762E-09 | TRUE | 1.00 |
| T2D_fmap.HIVEP2.chr6:143058692 | rs9390022 | chr6:142735419 | T | C | 0.00516 | 0 | 2.081E-09 | TRUE | 0.88 |
| T2D_fmap.INS-IGF2-KCNQ1.chr11:2235129 | rs1548432 | chr11:2310317 | G | A | 0.00049 | 1 | 2.645E-29 | TRUE | 0.11 |
| T2D_fmap.INS-IGF2-KCNQ1.chr11:2235129 | rs800140 | chr11:2312445 | A | G | 0.00278 | 2 | 2.893E-41 | TRUE | 0.11 |
| T2D_fmap.INS-IGF2-KCNQ1.chr11:2364549 | rs1548432 | chr11:2310317 | G | A | 0.00091 | 1 | 2.645E-29 | TRUE | 0.10 |
| T2D_fmap.INS-IGF2-KCNQ1.chr11:2364549 | rs179432 | chr11:2532323 | G | A | 0.00023 | 0 | 8.065E-17 | TRUE | 0.10 |
| T2D_fmap.INS-IGF2-KCNQ1.chr11:2364549 | rs800140 | chr11:2312445 | A | G | 0.01390 | 2 | 2.893E-41 | TRUE | 0.10 |
| T2D_fmap.INS-IGF2-KCNQ1.chr11:2375458 | rs800140 | chr11:2312445 | A | G | 0.00659 | 2 | 2.893E-41 | TRUE | 0.14 |
| T2D_fmap.INS-IGF2-KCNQ1.chr11:2799679 | rs2237884 | chr11:2778449 | T | C | 0.99996 | 0 | 3.052E-04 | TRUE | 1.00 |
| T2D_fmap.ITPR2.chr12:26474867 | 12:26472562_A_G | chr12:26319629 | A | G | 0.10745 | 1 | 6.319E-15 | TRUE | 1.00 |
| T2D_fmap.KLHDC5.chr12:27964996 | 12:27962103_C_T | chr12:27809170 | C | T | 0.07055 | 1 | 8.858E-17 | TRUE | 1.00 |
| T2D_fmap.KSR2.chr12:118412373 | 12:118410302_G_T | chr12:117972497 | G | T | 0.09793 | 7 | 1.170E-55 | TRUE | 0.71 |
| T2D_fmap.MAP3K11.chr11:65326154 | rs11227217 | chr11:65539931 | C | T | 0.02347 | 0 | 7.449E-04 | TRUE | 1.00 |
| T2D_fmap.MAP3K11.chr11:65326154 | rs12789028 | chr11:65558683 | G | A | 0.59053 | 17 | 8.011E-102 | TRUE | 1.00 |
| T2D_fmap.MHCregion.chr6:32439077 | rs1049213 | chr6:32659996 | A | G | 0.00070 | 0 | 8.734E-04 | TRUE | 0.59 |
| T2D_fmap.MNX1.chr7:156992461 | rs34498858 | chr7:157138643 | T | C | 0.00961 | 0 | 2.037E-08 | TRUE | 0.99 |
| T2D_fmap.MPHOSPH9-ZNF664.chr12:124545435 | 12:124560457_C_T | chr12:124075910 | C | T | 0.01152 | 0 | 1.063E-03 | TRUE | 0.93 |
| T2D_fmap.PIM3.chr22:50356302 | rs36138276 | chr22:49983919 | G | A | 0.02336 | 1 | 3.696E-07 | TRUE | 1.00 |
| T2D_fmap.PIM3.chr22:50356302 | rs56150947 | chr22:49960271 | G | A | 0.13048 | 2 | 1.283E-30 | TRUE | 1.00 |
| T2D_fmap.PVT1.chr8:129569999 | rs10113762 | chr8:128539956 | T | C | 0.00073 | 18 | 4.353E-105 | TRUE | 1.00 |
| T2D_fmap.PVT1.chr8:129569999 | rs57593539 | chr8:128555269 | G | A | 0.00252 | 0 | 1.077E-10 | TRUE | 1.00 |

| | | | | | | | | | |
|---|---|---|---|---|---|---|---|---|---|
| T2D_fmap.PVT1.chr8:129569999 | rs6651252 | chr8:128554935 | T | C | 0.00327 | 0 | 2.054E-14 | TRUE | 1.00 |
| T2D_fmap.PVT1.chr8:129569999 | rs6651253 | chr8:128555046 | G | C | 0.00352 | 3 | 2.160E-26 | TRUE | 1.00 |
| T2D_fmap.PVT1.chr8:129569999 | rs7824269 | chr8:128540098 | C | T | 0.12298 | 20 | 2.707E-159 | TRUE | 1.00 |
| T2D_fmap.PXK.chr3:58338809 | rs17355543 | chr3:58306573 | C | T | 0.00305 | 3 | 2.784E-21 | TRUE | 0.98 |
| T2D_fmap.PXK.chr3:58338809 | rs6779083 | chr3:58452408 | T | C | 0.00236 | 0 | 1.060E-13 | TRUE | 0.98 |
| T2D_fmap.RALY.chr20:32674967 | rs2284379 | chr20:34006836 | T | C | 0.02255 | 18 | 4.355E-43 | TRUE | 1.00 |
| T2D_fmap.RALY.chr20:32674967 | rs4911408 | chr20:34112760 | G | A | 0.02643 | 0 | 5.637E-04 | TRUE | 1.00 |
| T2D_fmap.RALY.chr20:32674967 | rs4911409 | chr20:34122904 | G | A | 0.00138 | 1 | 3.404E-02 | FALSE | 1.00 |
| T2D_fmap.SCD5.chr4:83587562 | rs10471048 | chr4:82666409 | G | C | 0.74719 | 11 | 2.560E-68 | TRUE | 1.00 |
| T2D_fmap.SEC16B.chr1:177889025 | rs574367 | chr1:177904075 | G | T | 0.07784 | 21 | 3.094E-155 | TRUE | 1.00 |
| T2D_fmap.SLC9B1.chr4:103725894 | rs223333 | chr4:102868653 | C | A | 0.00145 | 0 | 6.683E-10 | TRUE | 0.96 |
| T2D_fmap.SLC9B1.chr4:103725894 | rs223387 | chr4:102828359 | C | G | 0.00113 | 0 | 1.269E-05 | TRUE | 0.96 |
| T2D_fmap.SLC9B1.chr4:103725894 | rs223390 | chr4:102827640 | A | G | 0.00113 | 0 | 2.006E-04 | TRUE | 0.96 |
| T2D_fmap.SLC9B1.chr4:103725894 | rs223465 | chr4:102780174 | A | C | 0.00901 | 0 | 1.063E-03 | TRUE | 0.96 |
| T2D_fmap.SPRY2.chr13:80707429 | rs1215468 | chr13:80133294 | A | G | 0.89537 | 14 | 2.198E-87 | TRUE | 1.00 |
| T2D_fmap.SPRY2.chr13:80707429 | rs12428731 | chr13:80135460 | G | T | 0.07049 | 0 | 2.559E-08 | TRUE | 1.00 |
| T2D_fmap.STEAP1.chr7:89800241 | rs10257064 | chr7:90173282 | G | A | 0.08038 | 0 | 1.867E-16 | TRUE | 0.89 |
| T2D_fmap.STRBP.chr9:126015103 | rs1147322 | chr9:122913322 | A | G | 0.00390 | 0 | 1.098E-12 | TRUE | 0.96 |
| T2D_fmap.STRBP.chr9:126015103 | rs7044347 | chr9:123126007 | A | G | 0.00237 | 0 | 5.785E-07 | TRUE | 0.96 |
| T2D_fmap.TLE1.chr9:83998346 | rs10780510 | chr9:81344496 | T | C | 0.00567 | 21 | 0.000E+00 | TRUE | 1.00 |
| T2D_fmap.UBAP2.chr9:34074476 | rs12001437 | chr9:34074478 | T | C | 0.86633 | 0 | 1.371E-12 | TRUE | 0.70 |
| T2D_fmap.VWA5B1.chr1:20729451 | rs10916784 | chr1:20402958 | G | C | 0.67786 | 0 | 1.605E-04 | TRUE | 1.00 |
| T2D_fmap.ZMIZ1.chr10:80943841 | rs703977 | chr10:79184473 | T | G | 0.23905 | 22 | 5.760E-184 | TRUE | 1.00 |
| T2D_fmap.ZMIZ1.chr10:80943841 | rs703980 | chr10:79184084 | G | A | 0.27318 | 0 | 2.582E-06 | TRUE | 1.00 |

**Allelic imbalance association with intra-cellular total insulin content**

| **Genetic association credible set signal** | **Variant** | **Variant location** | **Referen ce alelle** | **Altern at e allele** | **Genetic association PPA** | **Phenotype** | **P-value** | **Standa rd error** | **Coeffi cient** | **Fraction of credible set variants that are heterozygous** |
|---|---|---|---|---|---|---|---|---|---|---|
| T2D_fmap.RALY.chr20:32674967 | rs2284379 | chr20:34006836 | T | C | 0.02255366 | insulin_content | 9.71E-11 | 0.03350 005 | 0.217 | 1.00 |
| T2D_fmap.GNAS.chr20:57387352 | rs1800900 | chr20:58840055 | A | G | 0.04849008 | insulin_content | 1.14E-05 | 0.06583 747 | 0.289 | 0.75 |

**Table S6.** Summary of antibodies and other primers related to STAR methods.

| **Antibodies used for Immunocytochemistry** | | | | | | | |
|---|---|---|---|---|---|---|---|
| **Usage** | **Antibody** | **Clone #** | **Host** | **Catalog #** | **Vendor** | **Dilution** | **Validation** |
| Immunostaining | Oct-4A (C30A3) Rabbit mAb | C30A3 | Rabbit | 2840 | Cell Signaling Technologies | 1:500 | https://www.cellsignal.com/products/primary-antibodies/oct-4a-c30a3-rabbit-mab/2840 |
| Immunostaining | Nanog (D73G4) XP® Rabbit mAb | D73G4 | Rabbit | 4903 | Cell Signaling Technologies | 1:500 | https://www.cellsignal.com/products/primary-antibodies/nanog-d73g4-xp-rabbit-mab/4903 |
| Immunostaining | SSEA4 (MC813) Mouse mAb | MC813 | Mouse | 4755 | Cell Signaling Technologies | 1:500 | https://www.cellsignal.com/products/primary-antibodies/ssea4-mc813-mouse-mab/4755 |
| Immunostaining | TRA-1-81 Mouse mAb | TRA-1-81 | Mouse | 4745 | Cell Signaling Technologies | 1:500 | https://www.cellsignal.com/products/primary-antibodies/tra-1-81-tra-1-81-mouse-mab/4745 |
| Immunostaining | Polyclonal Guinea Pig Anti-Insulin | Polyclonal | Guinea Pig | A0564 | Dako | 1:500 | https://www.citeab.com/antibodies/3382917-a0564-insulin |
| Immunostaining | Purified Rabbit Anti- Active Caspase-3 | C92-605 | Rabbit | 559565 | BD bioscience | 1:1000 | https://www.bdbiosciences.com/en-us/products/reagents/flow-cytometry-reagents/research-reagents/single-color-antibodies-ruo/purified-rabbit-anti-active-caspase-3.559565 |
| Immunostaining | Alexa Fluor 488 AffiniPure Donkey Anti-Guinea Pig IgG (H+L) | Polyclonal | Donkey | 706-545-148 | Jackson ImmunoResearch Labs | 1:1000 | https://www.jacksonimmuno.com/catalog/products/706-545-148 |
| Immunostaining | Donkey anti-Rabbit IgG (H+L) Highly Cross-Adsorbed Secondary Antibody, Alexa Fluor™ Plus 488 | Polyclonal | Donkey | A32790 | Thermo Fisher Scientific | 1:1,000 | https://www.thermofisher.com/antibody/product/Donkey-anti-Rabbit-IgG-H-L-Highly-Cross-Adsorbed-Secondary-Antibody-Polyclonal/A32790 |
| Immunostaining | Donkey anti-Mouse IgG (H+L) Highly Cross-Adsorbed Secondary Antibody, Alexa Fluor 594 | Polyclonal | Donkey | A-21203 | Thermo Fisher Scientific | 1:1,000 | https://www.thermofisher.com/antibody/product/Donkey-anti-Mouse-IgG-H-L-Highly-Cross-Adsorbed-Secondary-Antibody-Polyclonal/A-21203 |
| Immunostaining | Donkey anti-Rabbit IgG (H+L) Secondary Antibody, Alexa Fluor 594 conjugate | Polyclonal | Donkey | A-21207 | Thermo Fisher Scientific | 1:1,000 | https://www.thermofisher.com/antibody/product/Donkey-anti-Rabbit-IgG-H-L-Highly-Cross-Adsorbed-Secondary-Antibody- |

| | | | | | | | |
|---|---|---|---|---|---|---|---|
| | | | | | | | Polyclonal/A-21207 |

**Sequences of primers used in luciferase reporter assay**

| **Locus** | **SNP** | **Position** | **Region** | **Size** | **Primers for PCR amplication in cloning into PGL4.23 vectors** | | **Primers for PCR amplication in site-directed mutation** | |
|---|---|---|---|---|---|---|---|---|
| T2D_fmap.FAIM2. chr12:50263148 | rs7132908 | chr12:49869365 | 49868906 to 49869637 | 723 bp | Forward | CCGCTCGAGTCTGCAATGTTTCTTGAAAGCTGG | Forward | TTAGGGACTCTGGGCTGAGTGAGTGGCCAGTGAA |
| | | | | | Reverse | GGAAGATCTAAGGCTGAGTGAGGTGTGATT | Reverse | CTCAGCCCAGAGTCCCTAAGTGCTCCCCCA |
| | rs3205718 | chr12:49868026 | 49867719 to 49868368 | 650 bp | Forward | CCGCTCGAG TTA CAG CTT GTG ACA CCT GC | Forward | CTGTCCCTCCTTCCCCAAGCTTTTGGCACTCAGC |
| | | | | | Reverse | GGAAGATCTGGG TTT CTT GCT GTC TGG AAT | Reverse | AAGCTTGGGGAAGGAGGGACAGGTCCTCGG |

**Sequences of primers used in qRT-PCR**

| **Gene** | **Primers** | | **Gene** | **Primers** | |
|---|---|---|---|---|---|
| *INS* | Forward | AAGCGTGGCATTGTGGAA | *GAPDH* | Forward | CTGGGCTACACTGAGCACC |
| | Reverse | CTGCGTCTAGTTGCAGTAGTT | | Reverse | AAGTGGTCGTTGAGGGCAATG |

**Table S7.** Summary of RNA-seq sequencing statistics.

| Gene KO/WT | Library | Length of read | Mean insert size | Total number of read pairs | Number of uniquely mapped read pairs | % of uniquely mapped read pairs | Number of read pairs mapped to ambiguous genes | Number of read pairs mapped to unique genes | Number of read pairs unique gene CDS | Number of read pairs to unique gene UTRs | Number of read pairs not mapped to genes |
|---|---|---|---|---|---|---|---|---|---|---|---|
| ABCC8 | ABCC8_C1 | 51 | 413.02 | 39,125,068 | 32,099,466 | 82.04 | 1,635,543 | 26,644,529 | 19,641,387 | 7,003,142 | 1,730,808 |
| ABCC8 | ABCC8_C2 | 51 | 370.11 | 46,074,721 | 39,627,998 | 86.01 | 2,029,379 | 33,151,691 | 23,623,991 | 9,527,700 | 1,998,210 |
| ABCC8 | ABCC8_C3 | 51 | 324.02 | 39,329,205 | 34,512,150 | 87.75 | 1,486,723 | 28,081,180 | 19,074,070 | 9,007,110 | 2,263,971 |
| APOE | APOE_C1 | 51 | 477.51 | 73,732,437 | 60,257,740 | 81.72 | 2,452,222 | 46,319,766 | 33,365,281 | 12,954,485 | 4,508,507 |
| APOE | APOE_C2 | 51 | 442.65 | 98,792,082 | 83,781,201 | 84.81 | 2,970,771 | 62,503,547 | 41,104,045 | 21,399,502 | 10,872,994 |
| APOE | APOE_C3 | 51 | 415.5 | 123,518,245 | 106,818,854 | 86.48 | 3,910,565 | 79,955,120 | 54,757,233 | 25,197,887 | 12,471,354 |
| CDC123 | CDC123_C1 | 51 | 425.22 | 19,134,954 | 15,592,020 | 81.48 | 649,246 | 12,155,366 | 8,388,068 | 3,767,298 | 1,143,889 |
| CDC123 | CDC123_C2 | 51 | 407.62 | 31,375,050 | 26,221,976 | 83.58 | 1,208,668 | 20,566,672 | 14,834,053 | 5,732,619 | 1,694,116 |
| CDC123 | CDC123_C3 | 51 | 497.06 | 34,918,064 | 27,408,895 | 78.49 | 1,141,090 | 21,966,777 | 14,563,691 | 7,403,086 | 2,060,717 |
| CDKAL1 | CDKAL1_C1 | 51 | 397.04 | 26,447,298 | 21,774,389 | 82.33 | 1,041,179 | 17,646,121 | 12,410,312 | 5,235,809 | 1,375,271 |
| CDKAL1 | CDKAL1_C2 | 51 | 437.12 | 24,568,438 | 19,482,482 | 79.30 | 808,570 | 14,567,313 | 10,216,823 | 4,350,490 | 1,663,007 |
| CDKAL1 | CDKAL1_C3 | 51 | 429.11 | 22,568,102 | 18,463,733 | 81.81 | 833,044 | 14,240,474 | 10,029,188 | 4,211,286 | 1,325,038 |
| COBLL1 | COBLL1_C1 | 51 | 289.39 | 31,855,193 | 28,273,905 | 88.76 | 1,307,593 | 23,570,601 | 15,845,032 | 7,725,569 | 1,903,129 |
| COBLL1 | COBLL1_C2 | 51 | 288.81 | 34,401,707 | 30,317,837 | 88.13 | 1,357,726 | 25,153,567 | 16,810,076 | 8,343,491 | 2,105,048 |
| COBLL1 | COBLL1_C3 | 51 | 292.28 | 29,405,949 | 25,885,114 | 88.03 | 1,163,849 | 21,328,858 | 14,215,129 | 7,113,729 | 1,991,400 |
| GCKR | GCKR_C1 | 51 | 535.44 | 32,959,628 | 25,538,090 | 77.48 | 1,057,020 | 19,841,520 | 12,473,461 | 7,368,059 | 2,523,097 |
| GCKR | GCKR_C2 | 51 | 380.72 | 31,727,046 | 27,559,082 | 86.86 | 1,127,803 | 21,578,010 | 13,738,462 | 7,839,548 | 2,586,775 |
| GCKR | GCKR_C3 | 51 | 415.17 | 57,310,328 | 49,025,813 | 85.54 | 1,998,094 | 38,742,318 | 25,081,750 | 13,660,568 | 4,323,298 |
| GIPR | GIPR_C2 | 51 | 413.81 | 44,316,684 | 36,971,926 | 83.43 | 2,064,156 | 31,632,585 | 23,377,737 | 8,254,848 | 2,110,871 |
| GIPR | GIPR_C3 | 51 | 389.01 | 37,942,377 | 31,251,533 | 82.37 | 1,688,901 | 26,757,490 | 19,113,339 | 7,644,151 | 1,808,753 |
| GIPR | GIPR_C4 | 51 | 370.44 | 45,686,270 | 40,040,236 | 87.64 | 2,056,525 | 34,069,365 | 24,660,614 | 9,408,751 | 2,194,313 |
| HNF1A | HNF1A_C3 | 51 | 338.61 | 48,247,328 | 42,114,057 | 87.29 | 1,771,164 | 32,311,811 | 22,335,058 | 9,976,753 | 4,302,374 |
| HNF1A | HNF1A_C4 | 51 | 340.58 | 50,126,517 | 43,543,799 | 86.87 | 2,063,575 | 34,755,721 | 25,006,126 | 9,749,595 | 3,302,457 |
| HNF1A | HNF1A_S32 | 51 | 427.77 | 15,022,142 | 12,350,958 | 82.22 | 528,633 | 9,191,724 | 6,253,105 | 2,938,619 | 1,147,253 |
| HNF4A | HNF4A_C1 | 51 | 334.21 | 32,796,350 | 27,361,543 | 83.43 | 1,239,339 | 22,691,414 | 15,047,698 | 7,643,716 | 1,832,351 |
| HNF4A | HNF4A_C2 | 51 | 317.59 | 26,974,594 | 22,885,380 | 84.84 | 1,010,489 | 18,955,514 | 12,482,123 | 6,473,391 | 1,572,350 |
| HNF4A | HNF4A_C3 | 51 | 301.09 | 32,468,568 | 28,166,831 | 86.75 | 1,269,985 | 23,311,460 | 15,610,644 | 7,700,816 | 1,863,564 |
| HTT | HTT_C1 | 51 | 495.74 | 26,291,340 | 20,347,947 | 77.39 | 724,115 | 13,992,303 | 9,584,240 | 4,408,063 | 1,515,450 |
| HTT | HTT_C2 | 51 | 356.36 | 42,049,315 | 36,570,699 | 86.97 | 1,551,585 | 28,374,066 | 20,300,837 | 8,073,229 | 2,667,167 |
| HTT | HTT_C3 | 51 | 543.85 | 25,688,479 | 19,435,066 | 75.66 | 756,810 | 14,061,723 | 9,724,753 | 4,336,970 | 1,274,802 |
| IGF2BP2 | IGF2BP2_C1 | 51 | 603.7 | 27,931,924 | 21,017,715 | 75.25 | 848,859 | 15,948,623 | 10,826,029 | 5,122,594 | 2,210,729 |

| | | | | | | | | | | | |
|---|---|---|---|---|---|---|---|---|---|---|---|
| IGF2BP2 | IGF2BP2_C2 | 51 | 351.9 | 46,353,135 | 41,542,696 | 89.62 | 1,980,481 | 33,682,057 | 25,022,618 | 8,659,439 | 2,861,043 |
| IGF2BP2 | IGF2BP2_C3 | 51 | 405.84 | 41,073,687 | 35,253,249 | 85.83 | 1,559,919 | 28,028,021 | 19,998,671 | 8,029,350 | 2,855,413 |
| KCNJ11 | KCNJ11_C1 | 51 | 408.95 | 26,428,415 | 24,064,105 | 91.05 | 1,093,866 | 18,815,066 | 12,415,911 | 6,399,155 | 3,066,299 |
| KCNJ11 | KCNJ11_C2 | 51 | 388.66 | 35,433,392 | 31,903,914 | 90.04 | 1,212,584 | 24,809,579 | 15,315,579 | 9,494,000 | 4,148,825 |
| KCNJ11 | KCNJ11_C3 | 51 | 368.47 | 26,860,192 | 24,435,376 | 90.97 | 849,920 | 18,566,598 | 11,079,650 | 7,486,948 | 3,516,360 |
| SLC16A11 | SLC16A11_C1 | 51 | 317.78 | 30,899,338 | 26,300,457 | 85.12 | 1,383,888 | 22,127,971 | 14,944,899 | 7,183,072 | 1,394,087 |
| SLC16A11 | SLC16A11_C2 | 51 | 344.21 | 35,903,241 | 30,167,372 | 84.02 | 1,441,757 | 24,835,095 | 16,646,986 | 8,188,109 | 1,823,741 |
| SLC16A11 | SLC16A11_C3 | 51 | 324.04 | 27,744,938 | 23,621,599 | 85.14 | 1,107,571 | 19,380,914 | 13,073,331 | 6,307,583 | 1,518,564 |
| SLC30A8 | SLC30A8_C2 | 51 | 332.89 | 26,213,702 | 23,814,529 | 90.85 | 1,154,818 | 19,663,043 | 13,355,203 | 6,307,840 | 1,959,644 |
| SLC30A8 | SLC30A8_C3 | 51 | 343.24 | 22,969,930 | 20,716,842 | 90.19 | 888,253 | 16,321,126 | 10,190,935 | 6,130,191 | 2,427,235 |
| SLC30A8 | SLC30A8_C1 | 51 | 336.31 | 13,273,689 | 11,835,069 | 89.16 | 562,686 | 9,556,426 | 6,358,205 | 3,198,221 | 961,353 |
| TCF7L2 | TCF7L2_C1 | 51 | 436.55 | 27,924,258 | 23,375,588 | 83.71 | 1,126,335 | 19,042,854 | 13,531,867 | 5,510,987 | 1,572,477 |
| TCF7L2 | TCF7L2_C2 | 51 | 470.06 | 30,007,140 | 24,660,650 | 82.18 | 1,201,641 | 19,997,975 | 14,433,700 | 5,564,275 | 1,630,096 |
| TCF7L2 | TCF7L2_C3 | 51 | 399.29 | 42,971,956 | 37,674,163 | 87.67 | 1,872,003 | 30,762,871 | 22,049,055 | 8,713,816 | 2,522,449 |
| TGFB1 | TGFB1_C1 | 51 | 319.24 | 34,634,797 | 29,645,560 | 85.59 | 1,302,099 | 24,280,788 | 15,779,043 | 8,501,745 | 1,945,134 |
| TGFB1 | TGFB1_C2 | 51 | 298.01 | 35,439,096 | 31,269,651 | 88.23 | 1,337,159 | 25,646,838 | 17,143,366 | 8,503,472 | 2,167,315 |
| TGFB1 | TGFB1_C3 | 51 | 307.91 | 34,214,591 | 30,232,479 | 88.36 | 1,260,879 | 24,084,909 | 15,846,641 | 8,238,268 | 2,110,702 |
| TLE4 | TLE4_C1 | 51 | 555.38 | 30,063,849 | 23,292,392 | 77.48 | 903,735 | 18,547,195 | 12,673,469 | 5,873,726 | 1,737,055 |
| TLE4 | TLE4_C2 | 51 | 417.57 | 27,887,792 | 23,594,093 | 84.60 | 994,964 | 18,730,321 | 13,264,585 | 5,465,736 | 2,164,152 |
| TLE4 | TLE4_C3 | 51 | 616.83 | 29,207,573 | 21,849,935 | 74.81 | 901,947 | 17,054,533 | 12,122,901 | 4,931,632 | 2,002,297 |
| TMCC2 | TMCC2_C1 | 51 | 629.14 | 34,605,417 | 25,258,650 | 72.99 | 1,220,668 | 20,381,461 | 13,935,840 | 6,445,621 | 2,061,682 |
| TMCC2 | TMCC2_C2 | 51 | 442.36 | 25,536,821 | 21,497,172 | 84.18 | 1,024,980 | 16,729,006 | 11,481,531 | 5,247,475 | 1,897,764 |
| TMCC2 | TMCC2_C4 | 51 | 381.46 | 39,243,173 | 33,599,217 | 85.62 | 1,375,907 | 26,682,200 | 18,433,772 | 8,248,428 | 2,654,700 |
| WDR13 | WDR13_C1 | 51 | 298.45 | 39,211,900 | 34,376,186 | 87.67 | 1,580,387 | 26,794,103 | 17,666,388 | 9,127,715 | 2,422,635 |
| WDR13 | WDR13_C2 | 51 | 312.04 | 25,176,517 | 21,901,105 | 86.99 | 989,636 | 16,904,583 | 11,157,051 | 5,747,532 | 1,701,167 |
| WDR13 | WDR13_C3 | 51 | 326.35 | 27,080,315 | 23,177,684 | 85.59 | 1,057,926 | 18,052,323 | 11,813,653 | 6,238,670 | 1,625,020 |
| WFS1 | WFS1_C1 | 51 | 434.13 | 53,971,639 | 45,719,750 | 84.71 | 1,830,626 | 34,560,228 | 23,983,444 | 10,576,784 | 4,352,011 |
| WFS1 | WFS1_C2 | 51 | 436.27 | 46,757,818 | 39,227,852 | 83.90 | 1,755,824 | 30,232,336 | 21,069,048 | 9,163,288 | 3,331,852 |
| WFS1 | WFS1_C3 | 51 | 443.59 | 46,213,412 | 38,700,448 | 83.74 | 1,646,810 | 30,291,725 | 21,471,293 | 8,820,432 | 2,942,798 |
| WFS1 | WFS1_C4 | 51 | 402.36 | 52,985,376 | 45,131,696 | 85.18 | 1,951,892 | 35,072,485 | 24,756,066 | 10,316,419 | 3,567,266 |
| WT1 | WT1_C1 | 51 | 356.37 | 49,692,924 | 44,916,279 | 90.39 | 1,882,582 | 36,004,156 | 24,123,113 | 11,881,043 | 4,582,661 |
| WT1 | WT1_C2 | 51 | 387.93 | 18,452,014 | 15,183,336 | 82.29 | 626,673 | 12,157,013 | 7,836,170 | 4,320,843 | 1,486,275 |
| WT1 | WT1_C3 | 51 | 335.54 | 18,973,643 | 16,788,681 | 88.48 | 658,413 | 13,473,941 | 8,696,751 | 4,777,190 | 1,726,271 |
| WT2 | WT2_C1 | 51 | 539.33 | 24,359,019 | 18,897,791 | 77.58 | 915,341 | 14,680,292 | 9,818,925 | 4,861,367 | 1,714,231 |
| WT2 | WT2_C2 | 51 | 602.48 | 29,416,012 | 22,087,915 | 75.09 | 1,100,926 | 17,161,546 | 11,949,152 | 5,212,394 | 1,982,131 |
| WT2 | WT2_C3 | 51 | 470.46 | 41,348,172 | 34,135,512 | 82.56 | 1,733,572 | 27,352,431 | 19,048,280 | 8,304,151 | 2,736,812 |

**Supplementary Table 8:** Summary of ATAC-seq sequencing statistics.

| Gene KO/WT | Library | Length of read | Total number of reads | Number of reads mapped as primary alignment | % of reads mapped as primary alignment | Number of reads after duplicate removal | % of reads after duplicate removal |
|---|---|---|---|---|---|---|---|
| ABCC8 | ABCC8_C1 | 51 | 84,428,344 | 83,882,470 | 99.35 | 58,362,561 | 69.58 |
| ABCC8 | ABCC8_C2 | 51 | 84,981,754 | 84,372,690 | 99.28 | 60,375,891 | 71.56 |
| ABCC8 | ABCC8_C3 | 51 | 92,365,854 | 91,707,736 | 99.29 | 64,079,663 | 69.87 |
| APOE | APOE_C1 | 51 | 92,482,266 | 91,912,312 | 99.38 | 62,513,569 | 68.01 |
| APOE | APOE_C2 | 51 | 90,183,048 | 89,722,530 | 99.49 | 61,730,024 | 68.80 |
| APOE | APOE_C3 | 51 | 98,890,206 | 98,399,412 | 99.5 | 64,356,927 | 65.40 |
| CDC123 | CDC123_C1 | 51 | 94,143,948 | 93,589,984 | 99.41 | 61,048,151 | 65.23 |
| CDC123 | CDC123_C2 | 51 | 79,103,680 | 78,755,050 | 99.56 | 54,994,150 | 69.83 |
| CDC123 | CDC123_C3 | 51 | 95,747,730 | 95,039,495 | 99.26 | 65,654,387 | 69.08 |
| CDKAL1 | CDKAL1_C1 | 51 | 91,445,694 | 90,765,371 | 99.26 | 57,490,009 | 63.34 |
| CDKAL1 | CDKAL1_C2 | 51 | 105,343,018 | 104,661,690 | 99.35 | 67,554,777 | 64.55 |
| CDKAL1 | CDKAL1_C3 | 51 | 99,901,910 | 99,137,487 | 99.23 | 65,145,199 | 65.71 |
| COBLL1 | COBLL1_C1 | 51 | 92,751,776 | 92,014,103 | 99.2 | 65,467,979 | 71.15 |
| COBLL1 | COBLL1_C2 | 51 | 94,289,620 | 93,671,619 | 99.34 | 70,059,095 | 74.79 |
| COBLL1 | COBLL1_C3 | 51 | 97,161,006 | 96,507,335 | 99.33 | 70,707,272 | 73.27 |
| GCKR | GCKR_C1 | 51 | 89,829,616 | 89,143,543 | 99.24 | 64,492,630 | 72.35 |
| GCKR | GCKR_C2 | 51 | 98,819,882 | 98,076,511 | 99.25 | 65,619,773 | 66.91 |
| GCKR | GCKR_C3 | 51 | 70,146,700 | 69,670,342 | 99.32 | 48,786,771 | 70.03 |
| GIPR | GIPR_C1 | 51 | 91,952,726 | 91,287,496 | 99.28 | 65,788,147 | 72.07 |
| GIPR | GIPR_C2 | 51 | 88,665,104 | 87,969,138 | 99.22 | 62,124,342 | 70.62 |
| GIPR | GIPR_C3 | 51 | 102,548,446 | 101,785,491 | 99.26 | 66,294,589 | 65.13 |
| HNF1A | HNF1A_C1 | 51 | 77,221,758 | 76,806,490 | 99.46 | 54,892,213 | 71.47 |
| HNF1A | HNF1A_C2 | 51 | 91,286,992 | 90,583,759 | 99.23 | 61,497,732 | 67.89 |
| HNF1A | HNF1A_C3 | 51 | 85,069,568 | 84,041,319 | 98.79 | 60,011,452 | 71.41 |
| HNF4A | HNF4A_C1 | 51 | 96,061,418 | 95,351,953 | 99.26 | 68,469,375 | 71.81 |
| HNF4A | HNF4A_C2 | 51 | 76,864,132 | 76,368,432 | 99.36 | 56,199,464 | 73.59 |
| HNF4A | HNF4A_C3 | 51 | 88,456,956 | 87,835,017 | 99.3 | 62,083,968 | 70.68 |
| HTT | HTT_C1 | 51 | 106,885,050 | 106,123,147 | 99.29 | 69,646,168 | 65.63 |
| HTT | HTT_C2 | 51 | 86,846,052 | 86,165,091 | 99.22 | 60,961,911 | 70.75 |
| HTT | HTT_C3 | 51 | 86,801,332 | 86,116,683 | 99.21 | 58,522,115 | 67.96 |
| IGF2BP2 | IGF2BP2_C1 | 51 | 96,122,018 | 87,037,132 | 90.55 | 55,635,285 | 63.92 |
| IGF2BP2 | IGF2BP2_C2 | 51 | 97,705,956 | 92,469,307 | 94.64 | 61,991,102 | 67.04 |
| IGF2BP2 | IGF2BP2_C3 | 51 | 90,773,484 | 87,534,715 | 96.43 | 60,723,875 | 69.37 |

| | | | | | | | |
|---|---|---|---|---|---|---|---|
| KCNJ11 | KCNJ11_C1 | 51 | 106,823,510 | 84,486,042 | 79.09 | 55,832,225 | 66.08 |
| KCNJ11 | KCNJ11_C2 | 51 | 87,667,388 | 69,285,153 | 79.03 | 47,329,535 | 68.31 |
| KCNJ11 | KCNJ11_C3 | 51 | 106,185,520 | 83,822,266 | 78.94 | 55,393,994 | 66.09 |
| SLC16A11 | SLC16A11_C1 | 51 | 100,710,958 | 99,650,662 | 98.95 | 72,490,542 | 72.74 |
| SLC16A11 | SLC16A11_C2 | 51 | 105,204,762 | 104,286,227 | 99.13 | 50,055,213 | 48.00 |
| SLC16A11 | SLC16A11_C3 | 51 | 88,899,644 | 88,199,697 | 99.21 | 54,804,980 | 62.14 |
| SLC30A8 | SLC30A8_C1 | 51 | 92,904,418 | 80,585,742 | 86.74 | 60,644,141 | 75.25 |
| SLC30A8 | SLC30A8_C2 | 51 | 111,259,636 | 109,542,495 | 98.46 | 78,127,390 | 71.32 |
| SLC30A8 | SLC30A8_C3 | 51 | 93,137,096 | 92,446,580 | 99.26 | 66,483,877 | 71.92 |
| TCF7L2 | TCF7L2_C1 | 51 | 98,044,262 | 97,297,470 | 99.24 | 63,515,363 | 65.28 |
| TCF7L2 | TCF7L2_C2 | 51 | 96,203,106 | 90,557,715 | 94.13 | 62,184,287 | 68.67 |
| TCF7L2 | TCF7L2_C3 | 51 | 83,998,954 | 80,999,882 | 96.43 | 56,812,697 | 70.14 |
| TGFB1 | TGFB1_C1 | 51 | 107,917,412 | 97,518,275 | 90.36 | 70,326,665 | 72.12 |
| TGFB1 | TGFB1_C2 | 51 | 109,928,808 | 106,328,317 | 96.72 | 77,789,593 | 73.16 |
| TGFB1 | TGFB1_C3 | 51 | 106,106,190 | 95,041,428 | 89.57 | 66,093,081 | 69.54 |
| TLE4 | TLE4_C1 | 51 | 89,286,458 | 88,851,103 | 99.51 | 64,477,298 | 72.57 |
| TLE4 | TLE4_C2 | 51 | 99,666,218 | 99,000,672 | 99.33 | 67,314,520 | 67.99 |
| TLE4 | TLE4_C3 | 51 | 91,671,578 | 90,970,424 | 99.24 | 63,105,936 | 69.37 |
| TMCC2 | TMCC2_C1 | 51 | 92,375,892 | 91,868,148 | 99.45 | 69,186,781 | 75.31 |
| TMCC2 | TMCC2_C2 | 51 | 90,516,734 | 89,980,452 | 99.41 | 69,169,679 | 76.87 |
| TMCC2 | TMCC2_C3 | 51 | 96,133,986 | 95,389,505 | 99.23 | 71,595,979 | 75.06 |
| WDR13 | WDR13_C1 | 51 | 97,942,750 | 92,631,394 | 94.58 | 67,848,492 | 73.25 |
| WDR13 | WDR13_C2 | 51 | 93,595,892 | 90,832,341 | 97.05 | 67,403,984 | 74.21 |
| WDR13 | WDR13_C3 | 51 | 105,001,164 | 102,019,262 | 97.16 | 74,485,109 | 73.01 |
| WFS1 | WFS1_C1 | 51 | 92,832,248 | 92,204,937 | 99.32 | 59,552,823 | 64.59 |
| WFS1 | WFS1_C2 | 51 | 99,044,522 | 98,392,975 | 99.34 | 69,768,631 | 70.91 |
| WFS1 | WFS1_C3 | 51 | 93,177,732 | 92,342,097 | 99.1 | 67,871,821 | 73.50 |
| WFS1 | WFS1_C4 | 51 | 95,658,164 | 94,914,478 | 99.22 | 63,493,097 | 66.90 |
| WT1 | WT1_C1 | 51 | 98,474,116 | 92,450,733 | 93.88 | 62,788,140 | 67.92 |
| WT1 | WT1_C2 | 51 | 99,365,880 | 88,227,375 | 88.79 | 59,530,270 | 67.47 |
| WT1 | WT1_C3 | 51 | 100,514,354 | 91,419,141 | 90.95 | 61,235,380 | 66.98 |
| WT2 | WT2_C1 | 51 | 81,751,718 | 77,885,940 | 95.27 | 56,885,084 | 73.04 |
| WT2 | WT2_C2 | 51 | 98,837,102 | 94,216,444 | 95.32 | 68,156,352 | 72.34 |
| WT2 | WT2_C3 | 51 | 92,466,766 | 89,368,187 | 96.65 | 63,059,282 | 70.56 |
